# Access to unexplored regions of sequence space in directed enzyme evolution *via* insertion/deletion mutagenesis

**DOI:** 10.1101/790014

**Authors:** Stephane Emond, Maya Petek, Emily Kay, Brennen Heames, Sean Devenish, Nobuhiko Tokuriki, Florian Hollfelder

## Abstract

Insertions and deletions (InDels) are frequently observed in natural protein evolution, yet their potential remains untapped in laboratory evolution. Here we introduce a transposon mutagenesis approach (TRIAD) to generate libraries of random variants with short in-frame InDels, and screen TRIAD libraries to evolve a promiscuous arylesterase activity in a phosphotriesterase. The evolution exhibits features that are distinct from previous point mutagenesis campaigns: while the *average activity* of TRIAD variants is more deleterious, a *larger proportion* has successfully adapted for the new activity, exhibiting different functional profiles: (*i*) both strong and weak trade-off in original vs promiscuous activity are observed; (*ii*) trade-off is more severe (10- to 20-fold increased *k*_cat_/*K*_M_ in arylesterase with ∼100-fold decreases in the original phosphotriesterase activity) and (*iii*) improvements show up in *k*_cat_ rather than K_M_, suggesting novel adaptive solution. These distinct features make TRIAD an alternative to widely used point mutagenesis, providing access to functional innovations and traversing unexplored fitness landscape regions.

## INTRODUCTION

Directed evolution aims at identifying proteins with new functional traits by mimicking the natural process of genetic variation through mutations, followed by selection of improved variants. A major challenge for the success of this approach remains that only a very small fraction of the theoretically possible sequence space is accessible experimentally during any screening or selection process, so the type of library determines the success of directed evolution and the features of the functional proteins arising from such protein engineering. Expanding the diversity and the quality of gene libraries has been a major research focus to increase the chances of identifying new variants with desired functions. So far, most directed evolution (and, more generally, protein engineering) experiments have been performed using point substitutions for gene diversification, while <u>in</u>sertions or <u>del</u>etions (InDels) remain an overlooked source of variation despite their frequent and functionally beneficial occurrence in natural protein evolution ^1^. Combinatorial approaches to incorporate InDels at predefined positions, based on phylogenetic and/or structural analyses, have been developed to alter catalytic specificities of enzymes ^2^ or to improve the binding affinities of engineered antibodies^3^. While several methods for incorporating InDels randomly within a gene of interest have been developed, they show many limitations in terms of library quality and diversity. Most of these approaches generate frame-shifting InDels at high frequency (>66%) (*e.g.*, using error-prone DNA polymerases ^4, 5^, terminal deoxynucleotidyl transferase ^6^, exonucleases ^7, 8^, tandem duplication insertion ^9^ or truncation ^10^) and result in libraries that mostly consist of non-functional variants, which must be removed by high-throughput selection or screening. Methods based on the use of engineered transposons are designed to avoid frameshifts but so far have been limited to the generation of deletions ^11, 12^ or insertions of fixed length and defined sequences ^13^.

In the present work, a strategy for random introduction of single short in-frame InDels of one, two or three nucleotide triplets (± 3, 6 or 9 bp) into a given DNA sequence (dubbed TRIAD: **<u>T</u>**ransposition-based **<u>R</u>**andom **<u>I</u>**nsertion **<u>A</u>**nd **<u>D</u>**eletion mutagenesis) was established and validated by generating libraries of InDel variants of *Brevundimonas diminuta* phosphotriesterase (*wt*PTE), a highly efficient enzyme hydrolyzing the pesticide paraoxon ^14^ with promiscuous esterase and lactonase activities ^15^. The resulting TRIAD libraries were used to investigate the fitness effects of InDels on *wt*PTE and compare it to that of substitutions. Moreover, screening these libraries for improved arylesterase activity revealed several hits that would have been inaccessible using traditional and widely used point substitution mutagenesis approaches, demonstrating that the introduction of InDels can harvest functional diversity in previously unexplored regions of protein sequence space.

## RESULTS

### A strategy for creation of random InDel libraries

TRIAD consists of a single transposition reaction followed by successive cloning steps for the generation of deletions or insertions (Figure 1; see also Supplementary Figure S1 for a more detailed illustration). TRIAD’s first step is an *in vitro* Mu transposition reaction ^16^ that ultimately determines the location of the forthcoming single InDel event in each variant. The reaction is performed using engineered mini-Mu transposons, dubbed TransDel and TransIns (Supplementary Figure S2A), that are inserted randomly within the target DNA sequence during the first step of TRIAD, resulting in the generation of transposon insertion libraries. The ends of TransDel and TransIns were designed to bring about deletion and insertion libraries, respectively. TransDel is functionally equivalent to the previously described MuDel transposon ^11^ with recognition sites for the type IIS restriction enzyme MlyI at both ends. The positioning of MlyI sites within TransDel enables the deletion of 3 bp at random positions within the target sequence upon MlyI digestion and self-ligation (Figure 2A), as previously described ^11^. This strategy was extended to the generation of longer contiguous deletions (*i.e.*, -6 and -9 bp) with a second stage, involving the insertion and subsequent MlyI-mediated removal of custom-made cassettes (dubbed Del2 and Del3; Figure 1A and Figure 2A). For the generation of insertions, a new transposon, TransIns, was designed as – in contrast to TransDel – an *asymmetric* transposon (Figure 1B and Figure 2B), bearing different end sequences (NotI on one end and MlyI on the other). The latter site marks subsequent insertion sites for the ligation of custom-made shuttle cassettes: Ins1, Ins2 and Ins3 carrying one, two and three randomized nucleotide triplets, respectively. Further digestion using a type IIS restriction enzyme (AcuI) removes the shuttle sequence but leaves triplet insertions behind (Figure 1B and Figure 2B).

**Figure 1.**
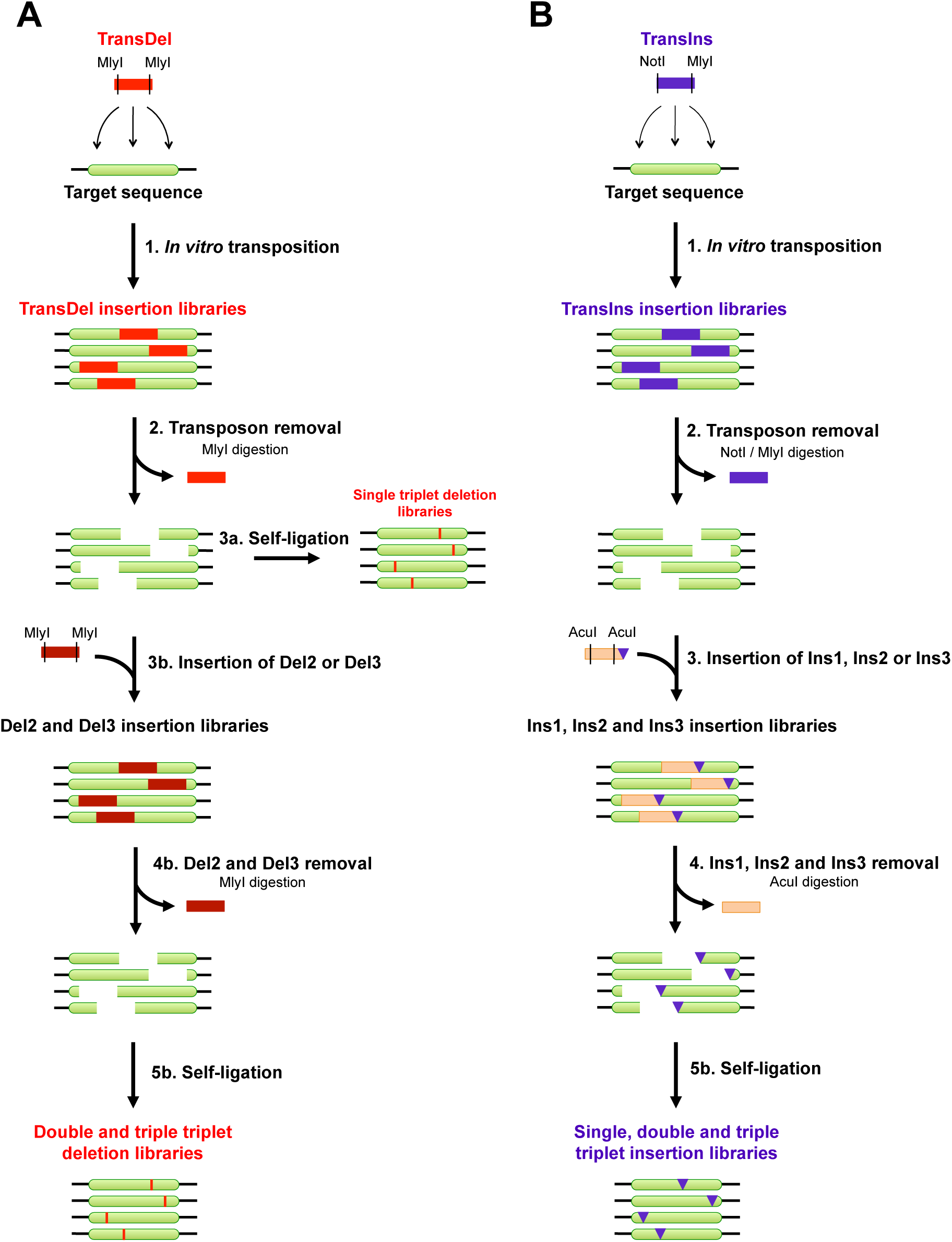
Schematic outline of TRIAD. **(A) Generation of deletion libraries.** *Step 1*: The TransDel insertion library is generated by *in vitro* transposition of the engineered transposon TransDel into the target sequence. *Step 2*: *Mly*I digestion removes TransDel together with 3 bp of the original target sequence and generate a single break per variant. *Step 3a*: self-ligation results in the reformation of the target sequence minus 3 bp, yielding a library of single variants with a deletion of one triplet ^11^. *Step 3b*: DNA cassettes dubbed Del2 and Del3 are then inserted between the break in the target sequence to generate Del2 and Del3 insertion libraries. *Step 4b*: *Mly*I digestion removes Del2 and Del3 together with 3 and 6 additional bp of the original target sequence, respectively. *Step 5b*: self-ligation results in the reformation of the target sequence minus 6 and 9 bp, yielding libraries of single variants with a deletion of 2 and 3 triplets, respectively. Deletions are indicated by red vertical lines. **(B) Generation of insertion libraries.** Step 1: The TransIns insertion library is generated by *in vitro* transposition of the engineered transposon TransIns into the target sequence. Step 2: digestion by *Not*I and *Mly*I removes TransIns. Step 3: DNA cassettes dubbed Ins1, Ins 3 and Ins3 (with respectively 1, 2 and 3 randomized NNN triplets at one of their extremities; indicated by purple triangles) are then inserted between the break in the target sequence to generate the corresponding Ins1, Ins2 and Ins3 insertion libraries. Step 4: *Acu*I digestion and 3’-end digestion by the Klenow fragment remove the cassettes, leaving the randomized triplet(s) in the original target sequence. Step 5: Self-ligation results in the reformation of the target sequence plus 3, 6 and 9 random bp, yielding libraries of single variants with an insertion of 1, 2 and 3 triplets, respectively.

**Figure 2.**
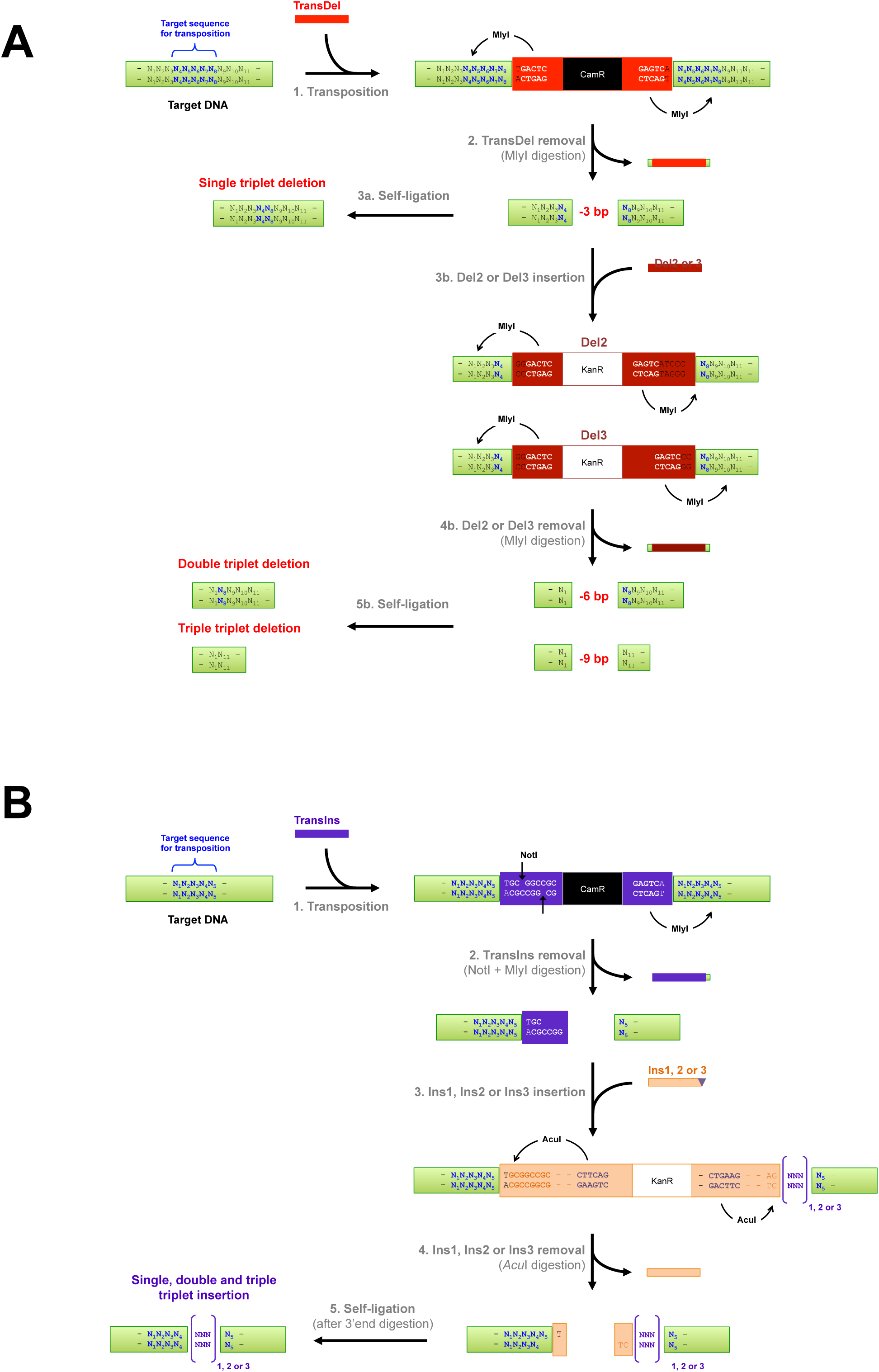
Mechanism for the generation of InDels using TRIAD. **(A) Generation of single, double and triple triplet nucleotide deletions.** Step 1. Two MlyI recognition sites (5’GAGTC(N)_5_↓) are positioned at each end of TransDel, 1 bp away from the site of transposon insertion. Transposition with TransDel results in the duplication of 5 bp (N_4_N_5_N_6_N_7_N_8_) of the target DNA at the insertion point. TransDel carries a selection marker (resistance gene against chloramphenicol; CamR) enabling the recovery of *in vitro* transposition products after transformation into *E. coli*. Step 2. MlyI digestion removes TransDel together with 8 bp of the target DNA (4 bp at each end), leaving blunt ends and resulting in the removal of a contiguous 3 bp sequence from the target DNA (N_5_N_6_N_7_). Step 3a. Self-ligation reforms the target DNA minus 3 bp, as previously described ^11^. Step 3b. Alternatively, blunt-ended cassettes Del2 or Del3 are ligated into the gap left upon TransDel removal for the generation of 6 and 9 bp deletions, respectively. Both Del2 and Del3 also contain two MlyI recognition sites advantageously positioned towards the ends of the cassettes. These cassettes also contain a different marker than TransDel (resistance gene against kanamycin; KanR) to avoid cross-contamination. Step 4b. MlyI digestion removes Del2 and Del3 together with respectively 3 and 6 additional bp of the original target DNA. In the case of Del2, MlyI digestion results in the removal of a 3 bp sequence (N_2_N_3_N_4_) on one side of the cassette. In the case of Del3, MlyI digestion results in the removal of two 3 bp sequence (N_2_N_3_N_4_) on both side of the cassette (N_2_N_3_N_4_ and N_8_N_9_N_10_). Step 5b. Self-ligation reforms the target DNA minus 6 or 9 bp. **(B) Generation of single, double and triple randomized triplet nucleotide insertions.** Step 1. TransDel is an asymmetric transposon with MlyI at one end and NotI at the other end. Both recognition sites are positioned 1bp away from TransIns insertion site. Upon transposition, 5 bp (N_1_N_2_N_3_N_4_N_5_) of the target DNA are duplicated at the insertion point of TransIns. Step 2. Double digestion with NotI and MlyI results in the removal of TransIns. Digestion with MlyI removes TransIns with 4 bp (N_1_N_2_N_3_N_4_) of the duplicated sequence at the transposon insertion site. Digestion with NotI leaves a 5’, 4-base cohesive overhang. Step 3. DNA cassettes Ins1, Ins2 and Ins3 (Ins1/2/3) carrying complementary ends are ligated in the NotI/MlyI digested TransIns insertion site. Ins1, Ins2 and Ins3 carry respectively 1, 2 and 3 randomized bp triplets at their blunt-ended extremities ([NNN]_1,2 or 3_; indicated in purple). Ins1/2/3 contain two AcuI recognition sites (5’CTGAAG(16/14)) strategically positioned towards their ends. One site is located so that AcuI will cleave at the point where the target DNA joins Ins1/2/3. The other site is positioned so that AcuI will cut inside Ins1/2/3 to leave the randomized triplet(s) with the target DNA. Step 4. Digestion with AcuI removes Ins1/2/3 leaving 3’, 2-base overhangs with the target DNA (*i.e.*, 5’N_5_T on one end and 5’TC on the end carrying the randomized triplet(s)). Digestion with the Large Klenow fragment generates blunt ends by removing the overhangs. This step also enables to discard the extra nucleotide (N_5_) from the sequence duplicated during the transposition. Step 5. Self-ligation reforms the target DNA with one, two or three randomized nucleotide triplets.

### Generation of random InDel libraries by TRIAD

To validate TRIAD, we generated InDel libraries from the gene encoding a highly expressed variant of phosphotriesterase (*wt*PTE) that had been previously used as starting point to generate an efficient arylesterase by laboratory evolution ^17, 18^. To enable TRIAD, any recognition sequences for MlyI, NotI and AcuI in the target sequence or the plasmid containing the target sequence must be removed. A synthetic gene encoding *wt*PTE (Supplementary Figure S3) as well as dedicated cloning vectors (namely pID-T7 and pID-Tet; Supplementary Figure S4 and Supplementary Methods) were therefore designed and assembled prior to the construction of libraries. The generation of transposition insertion libraries with TransDel or TransIns was performed as previously described ^19^. Briefly, *in vitro* transposition was performed to integrate TransDel or TransIns (∼1 kbp) within the plasmid (pID-Tet, ∼2.7 kbp) carrying the *wt*PTE synthetic gene (∼1 kbp). Transformation of the transposition products into electrocompetent *E. coli* yielded >30,000 colonies per transposition reaction. The fraction of transformants with the transposon inserted into *wt*PTE (∼27% of the entire plasmid length) corresponds to >8,000 colonies, corresponding to >8-fold coverage of possible insertion sites (∼1,000) within *wt*PTE. The transposon is inserted randomly throughout the entire plasmid, so the fragments corresponding to the target sequence carrying the transposon (∼2 kbp) were isolated by restriction digestion and subcloned back into intact pID-Tet, thereby generating TransDel and TransIns transposition libraries. At this stage, transformation of these libraries into *E. coli* typically yielded >10^6^ colonies, maintaining oversampling of transposon insertion sites without skewing the distribution due to sampling.

The TransDel and TransIns insertion libraries were then used as starting material to generate six independent libraries of *wt*PTE InDel variants: three deletion (−3, -6 and -9 bp) and three insertion libraries (+3, +6 and +9 bp). Without taking into account potential redundancy in the target DNA sequence, the maximal theoretical diversity of TRIAD libraries is a product of the number of positions (∼1000 bp for *wt*PTE) and the diversity introduced at each position: one deletion of each length for deletion libraries and the diversity of randomized triplets (64^1^, 64^2^ and 64^3^ for one, two or three NNN triplets) for insertion libraries. Therefore, the maximal theoretical diversity for *wt*PTE is ∼1000 variants in each deletion library and 6.4×10^4^, ∼4.1×10^6^, and ∼2.6×10^8^ for +3, +6 and +9 bp insertion libraries (see Supplementary Figure 5). However, depending on the sequence context, two or more neighbouring events may result in identical final DNA sequence, which reduces the accessible theoretical diversity. Theoretical diversities at the protein level are further reduced due to codon degeneracy and occurrence of stop codons as a result of certain InDels (Supplementary Figures S5B and S5C). Practically, the size of our libraries was limited by transformation efficiency, achieving > 10^6^ variants upon transformation into *E. coli.* Therefore, all deletions as well as +3 bp insertions were oversampled such that the library diversity was maintained between transformations, while the diversity of sampled transposition sites was maintained in larger +6 bp and +9 bp insertion libraries, with only a fraction of theoretical library diversity generated from the outset.

### Quality assessment of TRIAD libraries

The quality of the TRIAD libraries was assessed with Sanger sequencing to obtain long read accurate information, as well as deep next-generation sequencing to quantify the library sizes, distribution and diversity of InDels over the target sequence, and the transposition bias. All 121 Sanger-sequenced variants displayed only a single modification from the initial transposon insertion, without any incidental mutations, and 90 among them showed anticipated in-frame InDel mutations (see Supplementary Results 1.3 for details). We then obtained a next-generation sequencing dataset containing ∼1×10^6^ total 75-bp reads per deletion library and >3×10^6^ reads per insertion library (Supplementary Methods 2.3 - 2.7; Supplementary Figure S6 and Supplementary Table S4). In all libraries, the targeted in-frame InDels were found in high abundance, reaching more than 10^5^ variants detected by deep sequencing in the most diverse +6 bp and +9 bp libraries (> 10^3^ unique deletions and > 10^5^ unique insertions overall; Table 1). In agreement with Sanger sequencing of individual variants (Supplementary Table S1, Supplementary Results 1.3), frameshifts were rare in the -3 bp deletion library (4%) and more frequent (>20%) in the others (Supplementary Table S4B). Analysis of -3 bp and +3 bp libraries showed that TransDel and TransIns insertion accessed 85% and 95% of all possible DNA positions, respectively.

**Table 1.**
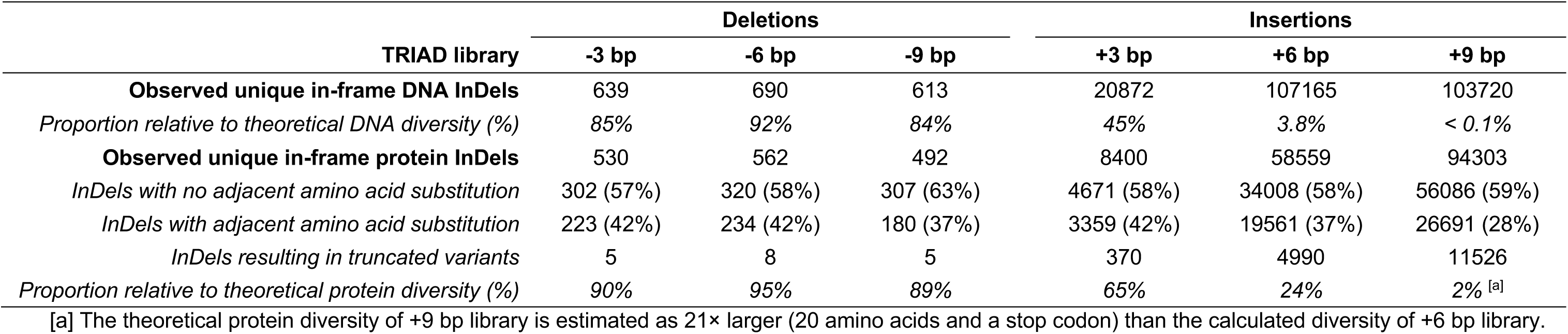
Mutagenesis efficiency of TRIAD analysed by deep sequencing. Unique in-frame InDels (*i.e.*, InDels of multiple of three nucleotides) were counted both at the DNA and the protein level. Adjacent amino acid substitutions and truncations (resulting from the occurrence of stop codons) may occur depending on the insertion point of the transposon, resulting in a lower value for the number of observed unique protein InDels. The proportion relative to the theoretical diversity accessible from the *wt*PTE sequence (both at the DNA and the protein level) was calculated as the ratio between the number of unique in-frame InDels observed by deep sequencing and the theoretical diversity for a given TRIAD library (see Supplementary Figure S5).

| TRIAD library | Deletions |  |  | Insertions |  |  |
| --- | --- | --- | --- | --- | --- | --- |
|  | -3 bp | -6 bp | -9 bp | +3 bp | +6 bp | +9 bp |
| **Observed unique in-frame DNA InDels** | 639 | 690 | 613 | 20872 | 107165 | 103720 |
| *Proportion relative to theoretical DNA diversity (%)* | 85% | 92% | 84% | 45% | 3.8% | < 0.1% |
| **Observed unique in-frame protein InDels** | 530 | 562 | 492 | 8400 | 58559 | 94303 |
| *InDels with no adjacent amino acid substitution* | 302 (57%) | 320 (58%) | 307 (63%) | 4671 (58%) | 34008 (58%) | 56086 (59%) |
| *InDels with adjacent amino acid substitution* | 223 (42%) | 234 (42%) | 180 (37%) | 3359 (42%) | 19561 (37%) | 26691 (28%) |
| *InDels resulting in truncated variants* | 5 | 8 | 5 | 370 | 4990 | 11526 |
| *Proportion relative to theoretical protein diversity (%)* | 90% | 95% | 89% | 65% | 24% | 2% <sup>[a]</sup> |

[a] The theoretical protein diversity of +9 bp library is estimated as 21× larger (20 amino acids and a stop codon) than the calculated diversity of +6 bp library.

Previous analysis of Mu transposon target site preference ^20^ suggests a strong preference for pyrimidines in position 2 and purines in position 4 of the 5 bp transposition site, based on 806 observed transpositions. By contrast, we observed similar frequencies for most deletions, with 52% of all detected deletions having between 10 and 99 reads per variant, and only 11% of all deletions (Supplementary Table S7) occurring more frequently across all three libraries combined (200 reads or more per variant; see distribution in Supplementary Figure S7). We extracted the weakly preferred transposition sequence to be 5’N-Py-G/C-Pu-N (see insert in Figure 3A; Supplementary Table S5) and we conclude that the sequence bias of Mu transposons is less pronounced than previously thought ^20, 21^.

**Figure 3.**
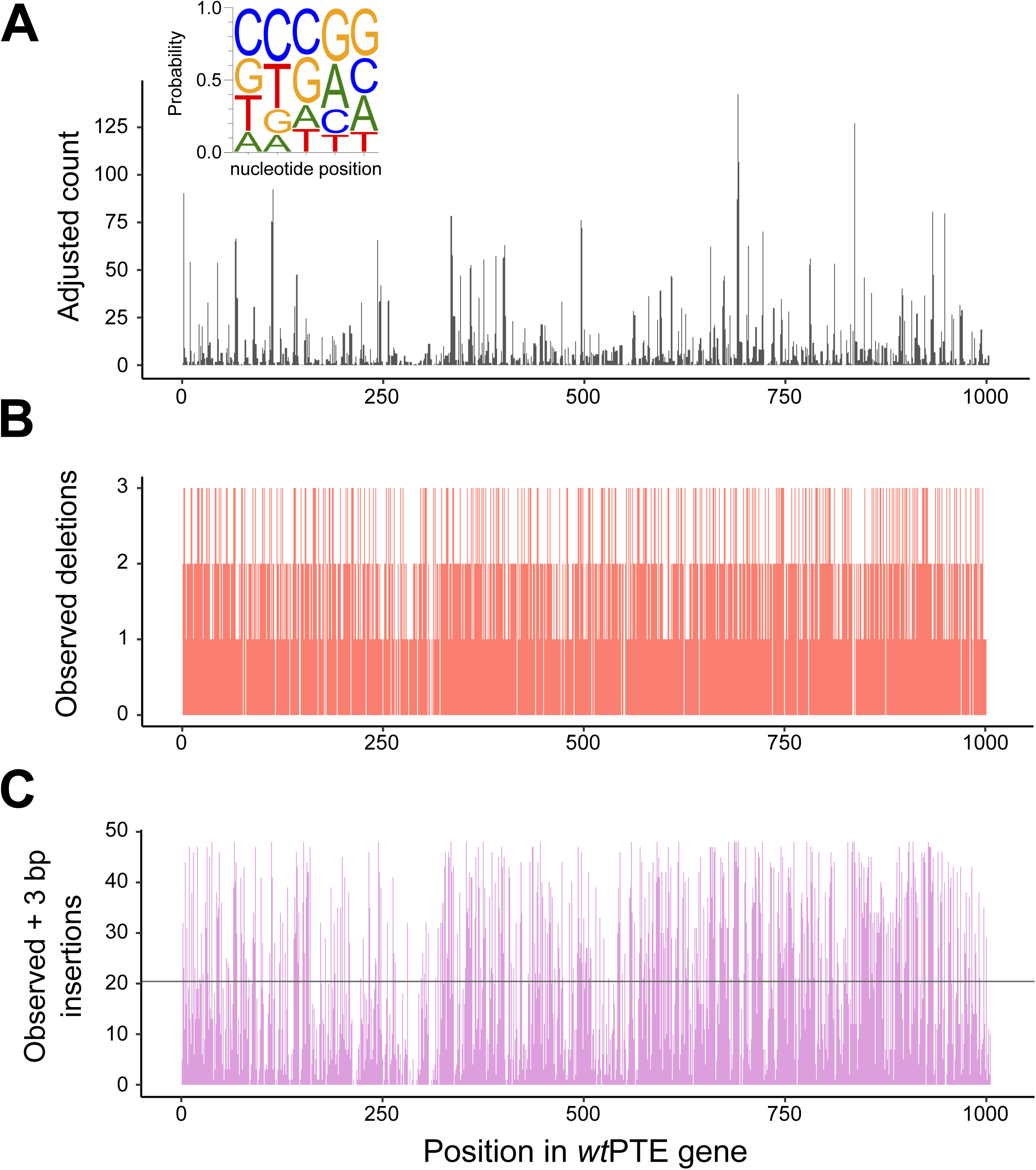
Mutagenesis efficiency of TRIAD. The composition of InDel libraries in the *wt*PTE gene was determined by deep sequencing and validated using Sanger sequences from randomly chosen variants. **(A)** Relative frequency of TransDel transposon insertion across *wt*PTE, derived from -3 bp deletions observed in deep sequencing and normalized for sequencing depth and InDel redundancy in DNA sequence (see Supplementary Methods 2.7). Insert: Relative site preference converted into the TransDel consensus insertion site. **(B)** Distribution and number of detected distinct DNA deletions in -3, -6 and -9 bp libraries combined per *wt*PTE position. **(C)** Distribution and number of observed +3 bp mutations per DNA position in +3 bp library, compared to the median 20.85 variants per position (horizontal line). Due to varying InDel redundancy depending on sequence context, the theoretical DNA diversity per position is between 42 and 48 variants (see Supplementary Figure S5). Analogous plots for +6 and +9 bp libraries are shown in Supplementary Figure S8.

Good coverage of possible positions in the insertion libraries translates into high diversity at most positions in *wt*PTE (Figure 3C; Supplementary Figure S8A): 10 or more distinct DNA insertions were observed at between 66% (+3 bp) and 80% (+6 and +9 bp libraries) of positions; furthermore, 100 or more insertions were detected in 34% (+6 bp) and 31% (+9 bp) of positions (Supplementary Figure S8B). While insertion libraries were sequenced with a higher loading onto the flow cell, this was still insufficient to fully capture the diversity in the +6 bp and +9 bp libraries (24% and 2% at the protein level, respectively; Table 1), where each variant was observed only once or twice (Supplementary Figure S7), and so the true diversity may be higher. When the transposition event does not align with codon boundaries, the resulting InDels exhibits an adjacent amino acid substitution: on protein level, an average of 39% of the InDels observed in the deep sequencing dataset of *wt*PTE variants exhibited such substitutions (Table 1). No significant bias was observed in the nucleotide composition of the in-frame insertions (Supplementary Figure S9), indicating that TRIAD generates diverse insertion variants.

Our quality assessment of the TRIAD libraries shows that - beyond a weak bias during transposon insertion - TRIAD libraries show excellent coverage of > 85% of positions in the DNA sequence of *wt*PTE. These results provide evidence that the TRIAD approach leads to large and diverse libraries of InDel variants through a set of straightforward cloning procedures that spanned just over 5 days (Supplementary Figure S1).

### Comparison of fitness effects between InDels and point substitutions

To compare the distribution of fitness effects of InDels *vs.* point substitutions, the levels of native phosphotriesterase (PTE; substrate: paraoxon; Figure 4) and promiscuous arylesterase (AE; substrate: 4-nitrophenyl butyrate, 4-NPB; Figure 4) activities were determined for several hundred *wt*PTE variants from each TRIAD library and from a trinucleotide substitution library (Figure 5; Supplementary Tables S8-9). Considering *wt*PTE is an evolutionarily “optimized” enzyme as a phosphotriesterase (based on the observation that it is operating near the diffusion limit for its native activity ^14^), it is to be expected that very few mutations would be beneficial and that InDels are more deleterious than point substitutions overall. This expectation is underlined by the observation that 83% of deletions and 77% of insertions are strongly deleterious (<0.1 PTE activity), compared to only 24% in the substitution library (Figure 5A). The average fitness change similarly favours substitutions and is an order of magnitude more deleterious for InDels (Figure 5C). However, of 485 screened deletions and 351 insertions, a total of 12 were beneficial (>1.5-fold PTE activity increase) against a background of already high catalytic efficiency. By contrast, no beneficial substitutions were found amongst the 342 substitutions screened. Similar frequencies were observed with respect to deleterious fitness changes induced by InDels *versus* point substitutions in *wt*PTE’s promiscuous arylesterase activity, with 76% of deletions and 62% of insertions strongly deleterious in comparison to only 19% of substitutions (Figure 5B; Supplementary Table S8). The frequency of InDels beneficial for arylesterase activity was found to be at least 3-fold higher than that of beneficial substitutions (6% and 7.7% for deletions and insertions, respectively *versus* 1.8% for substitutions; Figure 5B).

**Figure 4.**
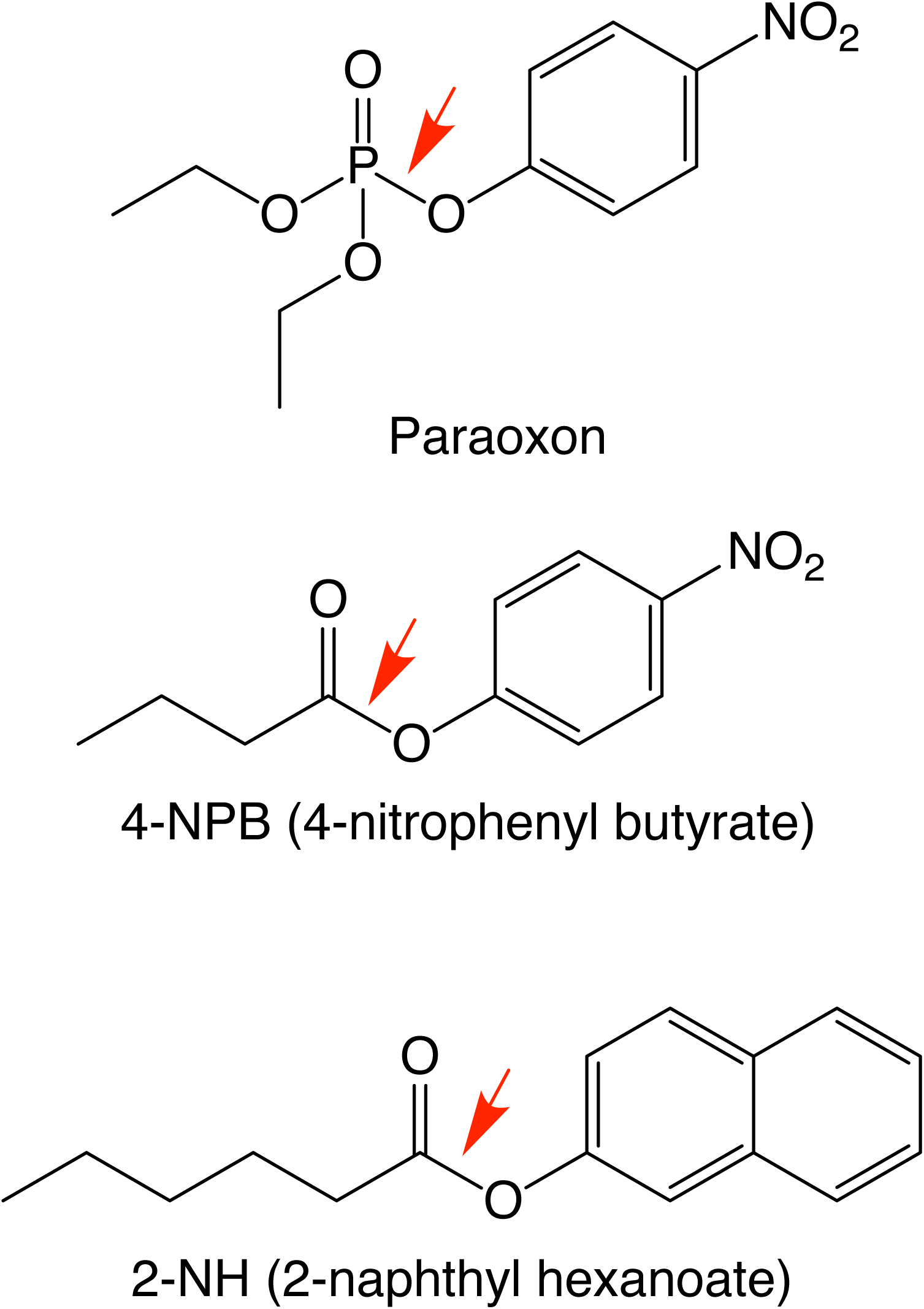
Structures of substrates. *wt*PTE catalyzes the hydrolysis of paraoxon (native substrate) and possess promiscuous activity against arylester substrates, *e.g.* 4-nitrophenyl butyrate and 2-naphthyl hexanoate.

**Figure 5.**
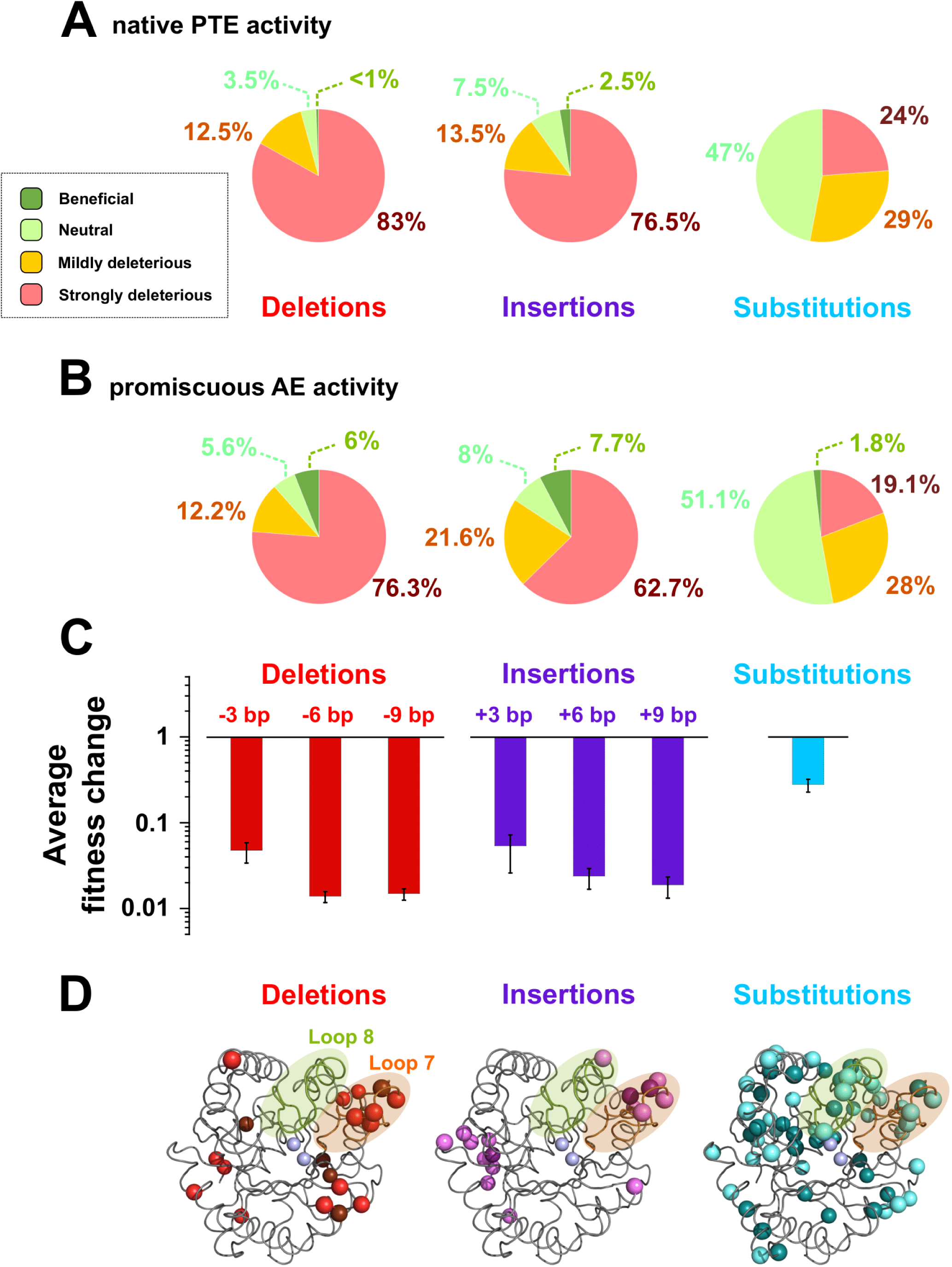
Fitness effects of InDels *versus* substitutions on *wt*PTE phosphotriesterase. **(A)** Distribution of fitness effects on phosphotriesterase activity (paraoxon). **(B)** Distribution of fitness effects on arylesterase activity (4-NPB). Fitness effects are classified as strongly deleterious (>10-fold activity decrease relative to *wt*PTE), mildly deleterious (10-fold to 1.5-fold decrease), neutral (<1.5-fold change), and beneficial (>1.5-fold increase). **(C)** Average fitness change in phosphotriesterase activity by deletions, insertions and substitutions. The average fitness change refers to the change in initial rates as a consequence of mutation and is calculated as the geometric mean of the relative activities of the variants measured as biological replicates (see Supplementary Table S8). Error bars indicate the corresponding confidence intervals (5% risk of error). **(D)** Structural mapping of protein changes observed in variants retaining ≥ 50% of *wt*PTE activity level (PDB ID 4PCP).

Mapping the observed mutations to the 3D structure of *wt*PTE provided insight into the location of adaptive InDels in comparison with point substitutions. While substitutions selected for ≥ 50% of *wt*PTE activity are found throughout the protein, the positions of InDels triggering similar functional effect appear more clustered in loops and on the surface (Figure 5D). Analysis of surface-accessible solvent area (SASA) suggests that mutations affecting the buried residues are more detrimental than surface-exposed ones (Supplementary Figure S10 and Supplementary Table S10). This observation holds for both InDels *and* substitutions. For substitutions, the correlation between SASA and fitness effects on activity is weak, while only ∼20% of neutral or beneficial InDels affect buried residues (cf. ∼40% of substitutions), readily explained by the larger impact of InDels on presumably optimised packing in the protein core.

### Screening and identification of adaptive InDels in *wt*PTE

To demonstrate that TRIAD libraries allow access to functional innovation *via* adaptive InDels, all the libraries generated from the full-length *wt*PTE gene (six libraries in total: -3, -6, -9, +3, +6 and +9 bp) were subjected to two parallel screening campaigns to identify variants with enhanced arylesterase activity against either 4-nitrophenyl butyrate (4-NPB) or 2-naphthyl hexanoate (2-NH) (Figure 4). Both screening campaigns consisted of a general two-step assay workflow. Upon transformation of the TRIAD libraries into *E. coli*, the resulting colonies (around 1 to 3×10^4^ per library) were first screened for either 1-naphthyl butyrate (prior to subsequent screening against 4-NPB in crude cell lysates) or 2-NH hydrolysis (using the FAST Red indicator that reacts with the released naphthol product). Colonies expressing an active variant (300 to 600 per library) were subsequently grown, lysed and tested for enzymatic activity (for either 4-NPB or 2-NH) in 96-well plates. Note that screening assays on colonies and in cell lysates were both performed after expression of *wt*PTE variants in the presence of overexpressed GroEL/ES chaperonin as described previously ^22^.

Overall, 81 hits (55 insertions and 26 deletions) were identified based on improved arylesterase activity against 2-NH or 4-NPB in cell lysates, with increases ranging from 2- to 140-fold in lysate activity compared to *wt*PTE (Table 2; Supplementary Table S11). In contrast with the adaptive substitutions previously identified ^18^, these adaptive InDels appeared to have a more drastic effect on the native phosphotriesterase activity, indicating a more severe trade-off *on average* between maintaining original and enhancing promiscuous activity (average specificity ratio ∼ 260; Supplementary Figure S11). However, numerous *individual* mutants that do not show such strong negative trade-off were also identified (*e.g.*, 64 variants out of 81 showed a specificity ratio < 100; Figures 6A-B; Supplementary Figure S12).

**Table 2.** Analysis of InDel *wt*PTE variants with at least 2-fold improved arylesterase activity. Values refer to the activity change of all or AE positive variants relative to wtPTE obtained by comparing the initial rates v_0_ for the hydrolysis of paraoxon, 4-NPB or 2-NH to that of *wt*PTE at 200 μM substrate concentration, resulting in a dimensionless ratio (recorded in Table 2). The average effect value was determined as the geometric mean of the relative activities of all the variants listed in Supplementary Table S11. The maximum, median and minimum changes correspond to the maximum, median and minimum relative activities for each substrate among the variants (See also Supplementary Figure S12).

|  |  | Activity fold change relative to wtPTE |  |  |  | Location of mutations <sup>[a]</sup> |  |  |  |
| --- | --- | --- | --- | --- | --- | --- | --- | --- | --- |
|  |  | Average effect | Median | Maximum | Minimum | Total | in Loop 7 | in Loop 8 | other locations |
| InDels | Paraoxon | 0.16 | 0.21 | 1.5 | <0.01 | 81 | 58 | 15 | 8 |
|  | 4-NPB | 3.0 | 2.8 | 14.4 | 2.0 | 56 | 50 | 1 | 5 |
|  | 2-NH | 7.4 | 6.4 | 138.6 | 2.6 | 25 | 8 | 14 | 3 |
| Deletions | Paraoxon | 0.08 | 0.15 | 0.86 | <0.01 | 26 | 17 | 4 | 5 |
|  | 4-NPB | 2.7 | 2.4 | 5.2 | 2.0 | 16 | 13 | 0 | 3 |
|  | 2-NH | 5.3 | 5.2 | 10.5 | 2.6 | 10 | 4 | 4 | 2 |
| Insertions | Paraoxon | 0.22 | 0.28 | 1.5 | <0.01 | 55 | 41 | 4 | 10 |
|  | 4-NPB | 3.2 | 2.9 | 14.4 | 2.1 | 40 | 37 | 1 | 2 |
|  | 2-NH | 9.5 | 8.1 | 138.6 | 3 | 15 | 4 | 10 | 1 |

[a] The values refer to the number of insertions and/or deletions observed in the entire sequence of wtPTE or in specific regions (e.g., loop 7 (residues 252-278) and loop 8 (residues 299-313)).

**Figure 6.**
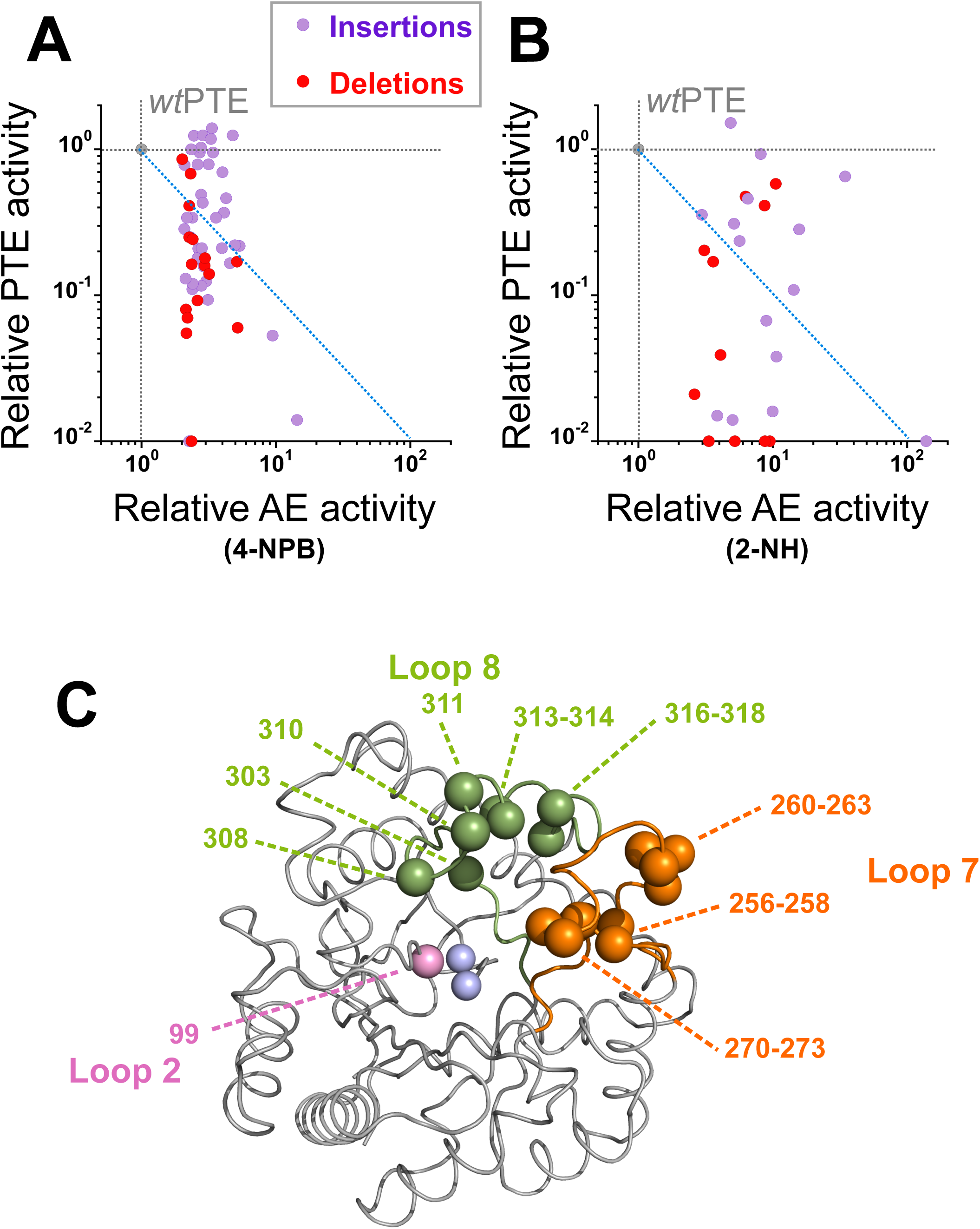
Identification of InDels improving the promiscuous arylesterase activity of *wt*PTE. **(A)(B)** Changes in phosphotriesterase (native; PTE) and arylesterase (promiscuous; AE) activities among *wt*PTE. InDel variants identified upon screening against butyrate (4-NPB; panel A) and hexanoate (2-NH; panel B) esters, respectively. The enzymatic activities for each variant (shown as grey dots) were measured in cell lysates and are plotted relative to those of *wt*PTE. Data are averages of triplicate values and error bars represent +/- 1 SEM. The dashed diagonal lines (in blue) demarcate the different trade-off regimes. Variants below the diagonal show a strong negative trade-off, with a large detriment to the original PTE activity (specificity ratio > 100), as AE activity is improved. Above the diagonal variants with weak trade-off emerge as generalists (specificity ratio < 100). **(C)** Position of the adaptive InDels in the PTE structure highlighting the frequency of mutations in loops 8, 7 and 2.

Sequence analysis of the nature and the location of the InDels responsible for the improvement in arylesterase activity (Table 2) showed that all the adaptive InDels (apart from one double triplet insertion, *e.g.*, V99G/Q99aI99b) were clustered in two flexible regions of *wt*PTE, namely loop 7 (residues L252 to Q278) and loop 8 (residues S299 to P322)(Figure 6C). Activity against 2-NH was improved by single InDels present in either loop while activity against 4-NPB was enhanced by InDels clustered in loop 7 (Table 2; Supplementary Figure S13). Unexpectedly, the best variant (10.5-fold improvement in AE) found in the -9 bp deletion library exhibited a 12 bp deletion (presumably as a result of a rearrangement during the transposition step in the TRIAD process) resulting in a four-amino acid residue deletion (*i.e.*, ΔA270-G273).

To further demonstrate that the identified InDels genuinely improve the arylesterase activity of *wt*PTE, the four variants exhibiting the strongest improvement against the 2-NH substrate (*i.e.*, ΔA270-G273, P256R/G256aA256b, S256aG256b and G311a) were purified and characterized to give a 10- to 20-fold increased *k*_*cat*_*/K*_*M*_ for 2-NH, while decreasing paraoxon hydrolysis by around 100-fold (Figure 6B; Table 3). The quantitation of the improvements varies between cell lysates and purified protein (*e.g.*, 140-fold in cell lysates versus 14-fold with the purified protein for P256R/G256aA256b), which may be ascribed to variation in expression levels.

**Table 3.** Kinetic properties of *wt*PTE variants.

| PTE variant <sup>[a]</sup> | Arylesterase (substrate: 2-NH) |  |  | Phosphotriesterase (substrate: paraoxon) |  |  | Reference |
| --- | --- | --- | --- | --- | --- | --- | --- |
| | $K_M$ ($\mu\text{M}$) | $k_{\text{cat}}$ ($\text{s}^{-1}$) | $k_{\text{cat}}/K_M$ ($\text{M}^{-1} \text{s}^{-1}$) | $K_M$ ($\mu\text{M}$) | $k_{\text{cat}}$ ($\text{s}^{-1}$) | $k_{\text{cat}}/K_M$ ($\text{M}^{-1} \text{s}^{-1}$) | |
| **wtPTE** | 179±21 | 0.075±0.004 | $4.2 \times 10^2$ | 57±5 | 1270±27 | $2.2 \times 10^7$ | Tokuriki *et al.*<sup>18</sup> |
| **H254R** | 250±32 | 0.27±0.01 | $1.1 \times 10^3$ | 7±1 | 62±3 | $8.9 \times 10^6$ | Tokuriki *et al.*<sup>18</sup> |
| **ΔA270L271L272G273** | 257±60 | 1.04±0.08 | $4 \times 10^3$ | 229±121 | 62±21 | $2.7 \times 10^5$ | This work<sup>[b]</sup> |
| **P256R/G256aA256b** | 777±137 | 4.7±0.4 | $6.04 \times 10^3$ | 103±51 | 15±3 | $1.45 \times 10^5$ | This work<sup>[b]</sup> |
| **S256aG256b** | 353±70 | 2.4±0.4 | $6.8 \times 10^3$ | 640±90 | 172±11 | $2.7 \times 10^5$ | This work<sup>[b]</sup> |
| **G311a** | 279±94 | 2.05±0.23 | $7.4 \times 10^3$ | 2566±136 | 346±121 | $1.35 \times 10^5$ | This work<sup>[b]</sup> |

[a] The symbol Δ before a residue (or a group of residues) signifies that this (or these) residue(s) have been deleted. Inserted residues are labelled using the number of the position after which they are inserted and alphabetical order (e.g., glutamine and tyrosine residues inserted in this order after the residues at position 230 would be labelled Q230aY230b). [b] *Conditions*: [S]=0–2 mM; [Tris-HCl]=100 mM (pH 7.5); [ZnCl<sub>2</sub>]=200 μM; [PTE]=10 nM (for paraoxon) and 500 nM (for 2-NH), respectively; T = 25 °C.

## DISCUSSION

Point substitutions, small insertions and deletions account for most evolutionary changes among natural proteins ^1^. The ratio of InDels to point substitutions covers a wide variety of ratios across different species, ranging from 1:5 in humans and primates ^23^ to 1:20 in bacteria ^24^, which indicates that InDels are typically subject to stronger purifying selection. Additionally, protein sequence alignments have established that the majority of InDels fixed in protein-coding genes are short (*i.e.*, encompassing 1 to 5 residues) and occur almost exclusively in loops linking secondary structure elements at the solvent-exposed surfaces of proteins ^25, 26, 27, 28, 29, 30^. While a large body of experimental evidence reports on the effects of substitutions, the impact of InDels on structural stability and functional divergence in protein evolution is still imperfectly understood, no doubt in part because convenient methods to introduce them in library experiments were missing. Substitutions, being merely side chain alterations, tend to have local effects with typically minor consequences for the overall structure of a protein. By contrast, InDels alter the length of the backbone, opening the way to dramatically larger changes in the packing and orientation of domains that may result in more global effect on the protein structure ^31, 32, 33^. Examples of InDels that cause significant repositioning of the backbone and nearby side chains to accommodate the extra or lost residues are on record ^34, 35, 36^. If such rearrangements occur near the active site of a protein, the resulting structural changes can change specificity and activity ^37, 38, 39^. Additionally, short InDels occurring at oligomerisation interfaces have also been shown to have important effects on the stability and/or specificity of protein complexes ^40, 41^. A corollary of the comparatively drastic effect of InDels on protein structure is the perception that they are more deleterious. Indeed, this view is now experimentally corroborated by our work on *wt*PTE (Figure 5) as well as a recent deep mutational scanning study investigating the fitness effects of single amino acid InDels on TEM-1 β-lactamase ^42^. However, InDels have also been shown to be contribute to functional divergence in several enzyme families, such as lactate and malate dehydrogenases ^43^, tRNA nucleotidyltransferases ^44^, nitroreductases ^45^, o-succinylbenzoate synthases ^41^, and phosphotriesterase-like lactonases ^2, 37^.

An experimental platform that gives straightforward access to InDel libraries makes it possible to analyse the respective contributions of InDels and point substitutions as sources of functional innovation in experiments against the molecular fossil record. The reliability of gene randomization methods is essential for success in directed evolution experiments. Popular and practically useful methods must meet several key requirements: a high-yielding library generation protocol should create a large number of variants, avoid bias in gene composition or type of variant introduced, and be technically straightforward. When it comes to amino acid substitutions, several approaches (e.g. error-prone PCR, site-saturation mutagenesis starting with synthetic oligonucleotides) have been developed that partially or fully meet these criteria and are widely used. By contrast, the use of InDels in directed evolution experiments has been curtailed by practical limitations in existing methodologies to randomly incorporate insertions and/or deletions (see Supplementary Table S13). Consequently, their application in protein engineering has been sparse, with very few directed evolution campaigns on record that originate from such libraries. For example, the RID protocol ^46^, the first attempt towards creating InDel libraries, relies on a complex protocol involving random cleavage of single stranded DNA, so that random substitutions are introduced unintentionally alongside the target mutations. Two other early methods, segmental mutagenesis ^8^ and RAISE ^6^, do not control for the length of the InDel and consequently produce libraries that primarily contain frameshifted variants. In contrast, a codon-based protocol dubbed COBARDE ^47^ gives a pool of multiple codon-based deletions with <5% frameshifts but requires custom reprogramming of an oligonucleotide synthesizer to create mutagenic oligonucleotides. Alternatively, the viability of transposon-based protocols has been established for generating deletions of various sizes, up to gene truncation variants ^10, 11, 12^. However, the only reported such protocol to create insertions, namely pentapeptide scanning mutagenesis ^13, 48^, merely gains access to insertions of defined size and sequence.

Improving on existing methodology, the TRIAD protocol meets all major requirements outlined above and gives easy access to large, diverse InDel libraries. The random insertion of a transposon gives excellent sampling of the entire target sequence (Figure 3; Supplementary Figure S8). Extensive sequencing shows that Mu transposon is even less biased than previously thought, so that functional effects upon insertion/deletion in *any* region of the protein can be taken advantage of. Library sizes upwards of 10^5^ variants were accessible by covering most of the theoretical diversity of up to two randomised amino acid insertions (Supplementary Figure S5). Introduction of randomised larger insertions exceeds the typically screenable library size, but can be constructed ^49^. Finally, the procedure is technically straightforward, consisting of transposition and cloning steps, and does not require access to specialized DNA synthesis equipment (as in ^47^). The TRIAD workflow is a versatile process that can be adapted to create libraries focused on a specific region of a protein, applicable in cases where screening throughput is limited. This approach would be analogous to other procedures (although only a few ^47, 50^ have directly exemplified this case). In the case of TRIAD, this was typically achieved by adding an in-frame seamless cloning step using a type IIS restriction enzyme such as SapI (see Supplementary Methods and Results; Supplementary Figure S12; Supplementary Table S12). InDel libraries constructed in this way showed good coverage of the target region, albeit with slightly more pronounced bias than whole-gene TRIAD, presumably due to increased sensitivity to preferential transposon insertions on a short target sequence. Alternatively, TRIAD can be further expanded with a recombination protocol (*e.g.*, DNA shuffling or Staggered Extension Process) to generate variants combining multiple InDels, which can be screened in a high-throughput assay ^51^.

The potential of InDel mutagenesis strategies in directed protein evolution is underlined by our comparative analysis of the fitness effect of InDels and point substitutions that showed InDels to be more likely to yield *wt*PTE variants with improved arylesterase activity than substitutions (Figure 5B). A second point of comparison are the evolutionary trajectories followed starting with InDel vs point substitution libraries. The promiscuous esterase activity of *wt*PTE has previously been used as the starting point of a directed evolution effort that generated an arylesterase which hydrolysed 2-NH with high efficiency ^18^. Here the mutation H254R, selected after the first round of mutagenesis, appeared to be a mutation on which the rest of the trajectory was highly contingent. InDel mutagenesis and selection puts us in a position to address the question whether alternative initial mutations would enable access to different evolutionary trajectories leading towards the same functional outcome. Based on the hypothesis that the use of a wider genetic and functional diversification (*i.e.*, by both substitutions and InDels) might lead to a wider diversity of possible evolutionary trajectories, the first objective was to identify new adaptive mutations improving the promiscuous arylesterase activity of *wt*PTE by screening InDel libraries of *wt*PTE generated *via* TRIAD. This resulted in the identification of multiple beneficial deletions and insertions, confirming that introduction of InDels can give rise to functional and improved catalysts.

We further observed that four of these adaptive InDels increase arylesterase activity 10- to 20-fold (in *k*_*cat*_*/K*_*M*_) against 2-NH, which is more than the 2.6-fold difference brought about by the initial H254R mutation from the previous directed evolution ^18^. For all four InDel variants, the improvement in 2-NH catalytic efficiency appears to be due to increased *k*_*cat*_ (from 13- to 67-fold), which outweighs an increased *K*_*M*_ in all four (from 2- to 6-fold). Similarly, all four variants increased in *K*_*M*_ for paraoxon (from 2 to 45-fold). On the other hand, the substitution H254R showed a different profile: it decreased *K*_*M*_ for paraoxon 8-fold, while hardly increasing it for 2-NH (1.4-fold) ^18^. Therefore, the top InDel hits in the cell lysate screening are more disruptive for both the binding of paraoxon and arylester (2-NH) substrates than substitutions, as may be expected for mutations that alter the backbone structure, while remaining beneficial overall, by improving turnover (*k*_*c*at_ being related, at least in first approximation, to the chemical reaction step, given the small difference in expression^52^).

Despite the scarcity of facile random InDel mutagenesis methods until recently, several examples of an adaptive role of InDels in protein directed evolution have been observed. Work on TEM-1 β-lactamase using the original Mu transposon-based triplet deletion libraries identified variants with increased resistance towards the antibiotic ceftazidime, up to 64-fold in minimum inhibitory concentration ^53^. A similar campaign that selected for eGFP variants with increased brightness in a colony screen identified the surprising eGFP-ΔGly4 deletion, which has significantly more cellular fluorescence likely due to increased refolding efficiency ^54^. Finally, a recent focused library approach in a PTE-like lactonase with insertions into loop 7 (that is shorter in lactonases) led to variants enhanced in phosphotriesterase activity, with increased *k*_*cat*_ and decreased *K*_*M*_ for paraoxon (*k*_*cat*_*/K*_*M*_ increased up to 600-fold) ^37^. Native lactonase activity was strongly affected in those variants with up to 10^4^-fold decreases in catalytic efficiency. These results in an enzyme closely related to PTE are very similar to our observations of the mixed effect of InDels on *wt*PTE, as explored based on the larger diversity of adaptive variants rendered available by TRIAD.

We conclude that unprecedented evolutionary trajectories become accessible by screening InDel libraries obtained *via* TRIAD, establishing a new paradigm that complements current strategies following the ‘one amino acid at the time’ adage ^55^ which are believed to lead to successful outcomes slowly, yet steadily. The effect of InDels is *on average* more deleterious than substitutions (Figure 5A), while the *fraction* of hits is increased in InDel libraries (Figure 5B), suggesting that InDel library strategies tend to ‘polarize’ properties of library members towards extremes. For thermodynamically more difficult reactions than those studied here, this trend to more extreme outcomes may practically imply low hit rates, in which case high throughput screening would become crucial. For example, ultrahigh-throughput screening based on droplet microfluidics ^56, 57^ could be combined with InDel mutagenesis to powerfully explore sequence space for evolutionary trajectories and individual variants that would not arise from epPCR mutagenesis libraries. It remains to be seen whether this new way of ‘jumping’ (rather than ‘tiptoeing’) across sequence space yields functionally better catalysts - or just different ones.

## METHODS

### Reagents

Paraoxon, 4-nitrophenyl butyrate (pNPB), 1-naphthyl butyrate (1-NB), 2-naphthyl hexanoate (2-NH), and Fast Red were purchased from Sigma. FastDigest restriction endonucleases, MuA transposase and T4 DNA ligase were purchased from Thermo Fisher Scientific. DNA Polymerase I, Large (Klenow) Fragment was purchased from New England Biolabs. All DNA modifying enzymes were used according to the manufacturer’s conditions. Oligonucleotides for PCR and adapter cloning experiments (Supplementary Table S14) were from Life Technologies and Sigma Aldrich.

### Plasmid and transposon construction

Detailed procedures and sequences can be found in the online Supplementary Information for the design and construction of transposons (TransDel and TransIns), cloning cassettes (Del2, Del3, Ins1, Ins2 and Ins3) and vectors (pID-T7, pID-Tet). The *wt*PTE gene lacking MlyI and AcuI sites was synthesised by GenScript (NJ, USA). InDel libraries of *wt*PTE prepared in the pID-Tet vector were subcloned with NcoI and HindIII into pET-strep vector ^22^ to express the strep-tag–PTE fusion protein for screening experiments and purification for the enzyme kinetics and stability assay. The trinucleotide substitution library of *wt*PTE used to compare the functional impact of InDels *vs.* point substitutions was generated following the TriNEx method ^58^ as described previously ^59^.

### Construction of InDel variant libraries of wtPTE

#### (1) Generation of transposon insertion libraries

The transposons TransDel and TransIns were extracted from pUC57 by BglII digestion and recovered by gel electrophoresis and purification. Insertion of TransDel or TransIns in the pID-Tet plasmid containing *wt*PTE was performed using *in vitro* transposition using 300 ng of plasmid, 50 ng of transposon and 0.22 μg MuA transposase in a 20 μL reaction volume. After incubation for 2 h at 30°C, the MuA transposase was heat-inactivated for 10 min at 75 °C. DNA products were purified and concentrated in 7 μL deionized water using a DNA clean concentrator kit (Zymo Research). 2 μL of the purified DNA was used to transform *E. coli* E. cloni^®^ 10G cells (> 10^10^ CFU/µg pUC19; Lucigen) by electroporation. The transformants (typically 10,000 to 50,000 CFU) were selected on LB agar containing ampicillin (amp; 100 μg/mL) and chloramphenicol (cam; 34 μg/mL). The resulting colonies were pooled, and their plasmid DNA extracted. The fragments corresponding to *wt*PTE containing the inserted transposon were obtained by double restriction digestion (NcoI/HindIII) followed by gel extraction and ligated in pID-Tet (50-100 ng). The ligation products were then transformed into electrocompetent *E. coli* E. cloni^®^ 10G cells. Upon selection on LB-agar-amp-cam, transformants (generally 1 to 2 × 10^6^ CFU) were pooled and their plasmid DNA extracted, yielding transposon (either TransDel or TransIns) insertion libraries.

#### (2) Generation of libraries of deletion variants

The construction of libraries with triplet deletion variants (−3 bp) was performed as previously described ^11^. TransDel insertion library plasmids were first digested with MlyI to remove TransDel. The fragments corresponding to linear pID-Tet-*wt*PTE plasmids (with a -3 bp deletion in *wt*PTE) were isolated by gel electrophoresis and purified. Self-circularization was then performed using T4 DNA ligase (Thermo Scientific) and 10-50 ng linearized plasmid (final concentration: ≥ 1 ng/μL). Upon purification and concentration, the ligation products were transformed into electrocompetent *E. coli* Ecloni^®^ 10G cells subsequently selected on LB-agar-amp, yielding a library of gene of interest variants with -3bp random deletions. For the construction of libraries of -6 and -9 bp deletion variants, cassettes Del2 and Del3 were ligated into the MlyI linearized plasmid (50-100 ng) in a 1:3 molar ratio. After purification and concentration, these ligation products were transformed into electrocompetent *E. coli* Ecloni^®^ 10G. The transformants (generally 1-3 × 10^6^ colony forming units, CFU) were selected on LB agar containing ampicillin (100 µg/L) and kanamycin (Kan; 50 μg/mL). The plasmids (corresponding to Del2 and Del3 insertion libraries) were extracted from the colonies and subsequently digested using MlyI to remove the Del2 and Del3 cassettes. The resulting linear pID-Tet-*wt*PTE products (containing the gene of interest with -6 or -9 bp deletions) were recovered by gel electrophoresis, purified and subsequently self-circularized. The resulting products were transformed into electrocompetent *E. coli* Ecloni^®^ 10G cells subsequently plated on LB-agar-amp, yielding libraries of *wt*PTE variants with -6 bp or -9 bp random deletions. All libraries were purified and stored in the form of plasmid solutions.

#### (3) Generation of libraries of insertion variants

TransIns insertion library plasmids were digested with NotI and MlyI to remove TransIns. The linearized pID-Tet-*wt*PTE plasmids were recovered by gel electrophoresis and purification. Cassettes Ins1, Ins2 and Ins3 were then inserted in the pID-Tet-*wt*PTE plasmid (50-100 ng) in a 1:3 molar ratio. After purification and concentration, these ligation products were transformed into electrocompetent *E. coli* Ecloni^®^ 10G and the transformants (generally 1.10^6^ to 3.10^6^ CFU) were selected on LB-agar-Amp-Kan. After extraction from the resulting colonies, the plasmids corresponding to Ins1, Ins2 and Ins3 insertion libraries were digested with AcuI. The linearized pID-Tet-*wt*PTE plasmids (with an insertion of 3, 6 and 9 bp in *wt*PTE) were recovered by gel electrophoresis, purified and subsequently treated with the Klenow fragment of DNA Polymerase I to remove 3’ overhangs created by AcuI digestion. After that blunting step, the plasmids were self-circularized. The resulting products were transformed into electrocompetent *E. coli* E. cloni^®^ 10G cells subsequently plated on LB-agar-amp, yielding libraries of *wt*PTE variants with +3, +6 or +9 bp random insertions. All the libraries were purified and stored in the form of plasmid solutions.

### Sequencing and quality analysis

The mutagenesis efficiency of TRIAD was analysed both by Sanger sequencing (Supplementary Tables S1-3) and deep sequencing. For the sequencing of individual *wt*PTE InDel variants obtained upon the transformation of libraries into *E. coli* (see above), individual colonies (∼20 per library; Supplementary Tables S1 and S2) were randomly picked for plasmid extraction and subsequent Sanger sequencing. For deep sequencing, libraries were digested from pID-Tet with FastDigest restriction enzymes Bpu1102I and Van91I to give a pool of 1.3 kb linear fragments, which were processed using Nextera DNA Library Preparation Kit according to manufacturer’s instructions and sequenced on Illumina MiSeq using 2×75 bp paired-end sequencing. The reads were de-multiplexed, adaptors trimmed and assembled using PEAR ^60^. Assembled and unassembled reads were mapped to the reference using Bowtie2 ^61^ and re-aligned to reference using the accurate Needleman-Wunsch algorithm with gap open penalty 15 and gap extend penalty 0.5 ^62^. Placing InDels in particular sequence contexts may be inherently ambiguous because of potential InDel redundancy: when two or more InDels inserted at *different* positions in the target gene result in *identical* final sequence, no algorithm will be able to distinguish between them and the resulting InDel is always assigned to a *single* arbitrarily chosen original insertion or deletion site (see the discussion of examples in the Supplementary Methods 2.7). No attempt was made to correct for such ambiguity at this point. Resulting alignments were used to count the number of reads in which the mutations occur, their type and position using in-house developed Python scripts (see Supplementary Methods 2.4 and 2.5). To analyse the sequence preference for TransDel transposition, the counts were corrected for codon ambiguity by dividing the observed count equally between all positions where the deletion could have originated.

### Screening procedures for *wt*PTE libraries

Prior to screening, InDel variant libraries of *wt*PTE were excised by NcoI/HindIII double digestion and subcloned into pET-Strep vector. The resulting libraries were transformed into *E. coli* BL21 (DE3) containing pGro7 plasmid for overexpression of the GroEL/ES chaperone system. Transformed cells (typically 2000-10000 CFU) were plated on agar-amp-cam. The colonies were replicated using a filter paper (BioTrace NT Pure Nitrocellulose Transfer Membrane 0.2 μm, PALL Life Sciences), which was placed onto a second plate containing IPTG (1 mM), ZnCl_2_ (200 μM) and arabinose (20% (w/v)) for chaperone overexpression. After overnight expression at room temperature, the filter paper was placed into an empty Petri dish and cells were lysed prior to the activity assay by alternating three times between storage at -20 °C and 37 °C. Subsequently, top agar (0.5% agar in 100 mM Tris–HCl pH 7.5) containing either 1-NB or 2-NH (200 μM) and FAST Red (200 μM) was layered and a red precipitate (resulting from the complex formation between Fast Red and the naphthol product) developed within ∼30 minutes. Colonies expressing an active PTE variant were picked, transferred in 96-well plates containing 200 μL LB-Amp-Cam per well and re-grown overnight at 30°C. Subsequently, 25 μL of the resulting cultures were used to inoculate 425 μL LB-Amp-Cam arabinose (20% (w/v)) in deep 96-well plates. After growth for 2 to 3 h at 30°C, expression of PTE variants was induced by adding IPTG (1 mM final concentration) and cultures were incubated for an additional 2 hours at 30 °C. Cells were then pelleted by centrifugation at 4 °C at maximum speed (3320×g) for 5-10 minutes and the supernatant removed. Pellets were frozen overnight at -80°C and, after thawing, lysed in 200 μL 50 mM Tris-HCl pH 7.5 supplemented with 0.1% (w/v) Triton-X100, 200 μM ZnCl_2_, 100 μg/mL lysozyme and 0.8 U/mL benzonase (Novagen). After 30 minutes of lysis, cell debris were spun down at 4 °C at 3320×g for 20 minutes. Enzyme assays were performed in 96-well plates containing a volume of 200 μL per well (20 μL pre-diluted lysate + 180 μL of 200 μM substrate in 50 mM Tris-HCl, pH 7.5 supplemented with Triton-X100 (0.02% in the case of paraoxon and 0.1% in the case of 2-NH/FR). The hydrolysis of paraoxon and 4-NPB were monitored by absorbance readings at 405 nm. The complex formation between 2-Naphthol and Fast Red was monitored at 500 nm.

### Purification of Strep-tagged PTE variants

pET-Strep-PTE plasmids were transformed into *E. coli* BL21 (DE3) grown for 8 h at 30 °C in Overnight Express Instant TB medium (Novagen) containing 100 µg/mL ampicillin and 200 μM ZnCl2 before lowering the temperature to 16 °C and continuing incubation overnight. Cells were harvested by centrifugation, resuspended and lysed using a 1:1 mixture of B-PER Protein Extraction Reagent (Thermo Scientific: 50 mM Tris-HCl buffer, pH 7.5 containing 200 μM ZnCl_2_, 100 μg/mL lysozyme and approx. 1 μL of benzonase per 100 mL. Cell debris was removed by centrifugation and the clarified lysate passed through a 45 μm filter before loading onto a Strep-Tactin Superflow High capacity column (1 mL). Strep-PTE variants were eluted with Elution buffer (100 mM Tris-HCl, pH 7.5, 200 μM ZnCl_2_ and 2.5 mM desthiobiotin) according to the manufacturer’s instructions (IBA Lifesciences).

### Kinetic characterization of PTE variants

Initial velocities (v_0_) were determined at 12 different substrate concentrations measured in triplicate, in the range of 2–2,000 µM in Tris-HCl (100 mM, pH 7.5) and ZnCl_2_ (200 µM). Rates of reaction were monitored by following the complex formation between the product and FastRed at 500 nm for 2-NH hydrolysis (in the presence of 2 mM FAST Red) and product formation at 405 nm for paraoxon hydrolysis. Purified protein was diluted to a concentration of 10 nM for paraoxon and 500 nM for 2-NH. *K*_*M*_ and *k*_*cat*_ were determined by fitting the initial rates at each concentration to the Michaelis-Menten model using KaleidaGraph (Synergy Software).

## Supporting information

EmondPetek Supplementary Information

## ACKNOWLEDGEMENTS

We would like to thank Christopher Kitching for assistance with analysis scripts and Charlotte Miton for helpful comments on the manuscript. SE, NT and SRAD held EU Marie Curie FP7 fellowships, FH is an H2020 ERC Advanced Investigator [695669]. This work was supported by the Biotechnology and Biological Sciences Research Council through a research grant [BB/L002469/1] and a studentship to MP [BB/M011194/1]. We acknowledge support from the Cambridge Service for Data Driven Discovery (CSD3) operated by the University of Cambridge Research Computing Service (http://www.csd3.cam.ac.uk/), provided by Dell EMC and Intel using Tier-2 funding from the Engineering and Physical Sciences Research Council (capital grant EP/P020259/1), and DiRAC funding from the Science and Technology Facilities Council (www.dirac.ac.uk).

## AUTHOR CONTRIBUTIONS

SE, NT and FH designed research. SE, MP, EK and BH performed experiments. MP and SE analysed deep sequencing data. SD was involved in early vector designs. SE, MP and FH wrote paper with input from all authors.

## DATA AVAILABILITY

Illumina raw sequencing reads were deposited with European Nucleotide Archive (https://www.ebi.ac.uk/ena) at accession number PRJEB28011. The source code along with instructions for all scripts involved in data processing are freely available at https://github.com/fhlab/TRIAD.

## SUPPLEMENTARY DATA

Supplementary results describing focused InDel libraries generated by TRIAD, supplementary methods, Supplementary Figures S1-S13 and supplementary tables S1-14 are available online.

## REFERENCES

1. Chothia C, Gough J, Vogel C, Teichmann SA. Evolution of the protein repertoire. Science 300, 1701–1703 (2003).

2. Afriat-Jurnou L, Jackson CJ, Tawfik DS. Reconstructing a missing link in the evolution of a recently diverged phosphotriesterase by active-site loop remodeling. Biochemistry 51, 6047–6055 (2012).

3. Lamminmaki U, et al. Expanding the conformational diversity by random insertions to CDRH2 results in improved anti-estradiol antibodies. J Mol Biol 291, 589–602 (1999).

4. Emond S, et al. A novel random mutagenesis approach using human mutagenic DNA polymerases to generate enzyme variant libraries. Protein Eng Des Sel 21, 267–274 (2008).

5. Kashiwagi K, Isogai Y, Nishiguchi K, Shiba K. Frame shuffling: a novel method for in vitro protein evolution. Protein Eng Des Sel 19, 135–140 (2006).

6. Fujii R, Kitaoka M, Hayashi K. Random insertional-deletional strand exchange mutagenesis (RAISE): a simple method for generating random insertion and deletion mutations. Methods Mol Biol 1179, 151–158 (2014).

7. Hida K, Won SY, Di Pasquale G, Hanes J, Chiorini JA, Ostermeier M. Sites in the AAV5 capsid tolerant to deletions and tandem duplications. Arch Biochem Biophys 496, 1–8 (2010).

8. Pikkemaat MG, Janssen DB. Generating segmental mutations in haloalkane dehalogenase: a novel part in the directed evolution toolbox. Nucleic Acids Res 30, E35–35 (2002).

9. Kipnis Y, Dellus-Gur E, Tawfik DS. TRINS: a method for gene modification by randomized tandem repeat insertions. Protein Eng Des Sel 25, 437–444 (2012).

10. Morelli A, Cabezas Y, Mills LJ, Seelig B. Extensive libraries of gene truncation variants generated by in vitro transposition. Nucleic Acids Res 45, e78 (2017).

11. Jones DD. Triplet nucleotide removal at random positions in a target gene: the tolerance of TEM-1 β-lactamase to an amino acid deletion. Nucleic Acids Res 33, e80 (2005).

12. Liu SS, Wei X, Ji Q, Xin X, Jiang B, Liu J. A facile and efficient transposon mutagenesis method for generation of multi-codon deletions in protein sequences. J Biotechnol 227, 27–34 (2016).

13. Hallet B, Sherratt DJ, Hayes F. Pentapeptide scanning mutagenesis: random insertion of a variable five amino acid cassette in a target protein. Nucleic Acids Res 25, 1866–1867 (1997).

14. Caldwell SR, Newcomb JR, Schlecht KA, Raushel FM. Limits of diffusion in the hydrolysis of substrates by the phosphotriesterase from Pseudomonas diminuta. Biochemistry 30, 7438–7444 (1991).

15. Roodveldt C, Tawfik DS. Shared promiscuous activities and evolutionary features in various members of the amidohydrolase superfamily. Biochemistry 44, 12728–12736 (2005).

16. Haapa S, Taira S, Heikkinen E, Savilahti H. An efficient and accurate integration of mini-Mu transposons in vitro: a general methodology for functional genetic analysis and molecular biology applications. Nucleic Acids Res 27, 2777–2784 (1999).

17. Kaltenbach M, Jackson CJ, Campbell EC, Hollfelder F, Tokuriki N. Reverse evolution leads to genotypic incompatibility despite functional and active site convergence. Elife 4, (2015).

18. Tokuriki N, Jackson CJ, Afriat-Jurnou L, Wyganowski KT, Tang R, Tawfik DS. Diminishing returns and tradeoffs constrain the laboratory optimization of an enzyme. Nat Commun 3, 1257 (2012).

19. Baldwin AJ, Arpino JA, Edwards WR, Tippmann EM, Jones DD. Expanded chemical diversity sampling through whole protein evolution. Mol Biosyst 5, 764–766 (2009).

20. Haapa-Paananen S, Rita H, Savilahti H. DNA transposition of bacteriophage Mu. A quantitative analysis of target site selection in vitro. J Biol Chem 277, 2843–2851 (2002).

21. Mizuuchi M, Mizuuchi K. Target site selection in transposition of phage Mu. Cold Spring Harb Symp Quant Biol 58, 515–523 (1993).

22. Tokuriki N, Tawfik DS. Chaperonin overexpression promotes genetic variation and enzyme evolution. Nature 459, 668–673 (2009).

23. Ng PC, et al. Genetic variation in an individual human exome. PLoS Genet 4, e1000160 (2008).

24. Chen JQ, Wu Y, Yang H, Bergelson J, Kreitman M, Tian D. Variation in the ratio of nucleotide substitution and indel rates across genomes in mammals and bacteria. Mol Biol Evol 26, 1523–1531 (2009).

25. Ajawatanawong P, Baldauf SL. Evolution of protein indels in plants, animals and fungi. BMC Evol Biol 13, 140 (2013).

26. Benner SA, Cohen MA, Gonnet GH. Empirical and structural models for insertions and deletions in the divergent evolution of proteins. J Mol Biol 229, 1065–1082 (1993).

27. Lin M, Whitmire S, Chen J, Farrel A, Shi X, Guo JT. Effects of short indels on protein structure and function in human genomes. Sci Rep 7, 9313 (2017).

28. Pascarella S, Argos P. Analysis of insertions/deletions in protein structures. J Mol Biol 224, 461–471 (1992).

29. Taylor MS, Ponting CP, Copley RR. Occurrence and consequences of coding sequence insertions and deletions in Mammalian genomes. Genome Res 14, 555–566 (2004).

30. Toth-Petroczy A, Tawfik DS. Protein insertions and deletions enabled by neutral roaming in sequence space. Mol Biol Evol 30, 761–771 (2013).

31. Grishin NV. Fold change in evolution of protein structures. J Struct Biol 134, 167–185 (2001).

32. Studer RA, Dessailly BH, Orengo CA. Residue mutations and their impact on protein structure and function: detecting beneficial and pathogenic changes. Biochem J 449, 581–594 (2013).

33. Toth-Petroczy A, Tawfik DS. Hopeful (protein InDel) monsters? Structure 22, 803–804 (2014).

34. Heinz DW, Baase WA, Dahlquist FW, Matthews BW. How amino-acid insertions are allowed in an alpha-helix of T4 lysozyme. Nature 361, 561–564 (1993).

35. O’Neil KT, Bach AC, 2nd, DeGrado WF. Structural consequences of an amino acid deletion in the B1 domain of protein G. Proteins 41, 323–333 (2000).

36. Stott KM, Yusof AM, Perham RN, Jones DD. A surface loop directs conformational switching of a lipoyl domain between a folded and a novel misfolded structure. Structure 17, 1117–1127 (2009).

37. Hoque MA, et al. Stepwise Loop Insertion Strategy for Active Site Remodeling to Generate Novel Enzyme Functions. ACS Chem Biol 12, 1188–1193 (2017).

38. Shortle D, Sondek J. The emerging role of insertions and deletions in protein engineering. Curr Opin Biotechnol 6, 387–393 (1995).

39. Park HS, Nam SH, Lee JK, Yoon CN, Mannervik B, Benkovic, Kim HS. Design and evolution of new catalytic activity with an existing protein scaffold. Science 311, 535–538 (2006).

40. Hashimoto K, Panchenko AR. Mechanisms of protein oligomerization, the critical role of insertions and deletions in maintaining different oligomeric states. Proc Natl Acad Sci U S A 107, 20352–20357 (2010).

41. Odokonyero D, et al. Loss of quaternary structure is associated with rapid sequence divergence in the OSBS family. Proc Natl Acad Sci U S A 111, 8535–8540 (2014).

42. Gonzalez CE, Roberts P, Ostermeier M. Fitness Effects of Single Amino Acid Insertions and Deletions in TEM-1 beta-Lactamase. J Mol Biol 431, 2320–2330 (2019).

43. Boucher JI, Jacobowitz JR, Beckett BC, Classen S, Theobald DL. An atomic-resolution view of neofunctionalization in the evolution of apicomplexan lactate dehydrogenases. Elife 3, (2014).

44. Neuenfeldt A, Just A, Betat H, Morl M. Evolution of tRNA nucleotidyltransferases: a small deletion generated CC-adding enzymes. Proc Natl Acad Sci U S A 105, 7953–7958 (2008).

45. Akiva E, Copp JN, Tokuriki N, Babbitt PC. Evolutionary and molecular foundations of multiple contemporary functions of the nitroreductase superfamily. Proc Natl Acad Sci U S A 114, E9549–E9558 (2017).

46. Murakami H, Hohsaka T, Sisido M. Random insertion and deletion of arbitrary number of bases for codon-based random mutation of DNAs. Nat Biotechnol 20, 76–81 (2002).

47. Osuna J, Yanez J, Soberon X, Gaytan P. Protein evolution by codon-based random deletions. Nucleic Acids Res 32, e136 (2004).

48. Hayes F, Hallet B. Pentapeptide scanning mutagenesis: encouraging old proteins to execute unusual tricks. Trends Microbiol 8, 571–577 (2000).

49. Jones DD, Arpino JA, Baldwin AJ, Edmundson MC. Transposon-based approaches for generating novel molecular diversity during directed evolution. Methods Mol Biol 1179, 159–172 (2014).

50. Tizei PAG, Harris E, Renders M, Pinheiro VB. InDel assembly: A novel framework for engineering protein loops through length and compositional variation. Bioarxiv, DOI: 10.1101/127829 (2017).

51. Skamaki K, et al. In vitro Evolution of Antibody Affinity via Insertional Mutagenesis Scanning of an Entire Antibody Variable Region. Submitted. (2019).

52. Raushel FM, Holden HM. Phosphotriesterase: an enzyme in search of its natural substrate. Adv Enzymol Relat Areas Mol Biol 74, 51–93 (2000).

53. Simm AM, Baldwin AJ, Busse K, Jones DD. Investigating protein structural plasticity by surveying the consequence of an amino acid deletion from TEM-1 beta-lactamase. FEBS Lett 581, 3904–3908 (2007).

54. Arpino JA, Reddington SC, Halliwell LM, Rizkallah PJ, Jones DD. Random single amino acid deletion sampling unveils structural tolerance and the benefits of helical registry shift on GFP folding and structure. Structure 22, 889–898 (2014).

55. Tracewell CA, Arnold FH. Directed enzyme evolution: climbing fitness peaks one amino acid at a time. Curr Opin Chem Biol 13, 3–9 (2009).

56. Colin PY, Zinchenko A, Hollfelder F. Enzyme engineering in biomimetic compartments. Curr Opin Struct Biol 33, 42–51 (2015).

57. Mair P, Gielen F, Hollfelder F. Exploring sequence space in search of functional enzymes using microfluidic droplets. Curr Opin Chem Biol 37, 137–144 (2017).

58. Baldwin AJ, Busse K, Simm AM, Jones DD. Expanded molecular diversity generation during directed evolution by trinucleotide exchange (TriNEx). Nucleic Acids Res 36, e77 (2008).

59. Kaltenbach M, Emond S, Hollfelder F, Tokuriki N. Functional Trade-Offs in Promiscuous Enzymes Cannot Be Explained by Intrinsic Mutational Robustness of the Native Activity. PLoS Genet 12, e1006305 (2016).

60. Zhang J, Kobert K, Flouri T, Stamatakis A. PEAR: a fast and accurate Illumina Paired-End reAd mergeR. Bioinformatics 30, 614–620 (2014).

61. Langmead B, Salzberg SL. Fast gapped-read alignment with Bowtie 2. Nat Methods 9, 357–359 (2012).

62. Rice P, Longden I, Bleasby A. EMBOSS: the European Molecular Biology Open Software Suite. Trends Genet 16, 276–277 (2000).

