## Supplementary material for "Access to unexplored regions of sequence space in directed enzyme evolution *via* insertion/deletion mutagenesis": EmondPetek Supplementary Information

Supplementary Figure S8: Number of distinct insertions observed per position in *wt*PTE. 31

### 1. SUPPLEMENTARY RESULTS

#### 1.1. Focused InDel libraries generated by TRIAD

TRIAD was additionally applied to focus the InDel mutagenesis on a specific targeted region within a protein by adding an in-frame seamless cloning step using a type IIS restriction enzyme such as SapI (strategy outlined in Supplementary Figure S12) [1]. This approach requires the target region to be extracted from its original gene and cloned in its own target plasmid with flanking SapI recognition sequences. This allows transposon integration into the target region in isolation from the rest of the gene. In parallel, an adapter plasmid is constructed, comprising the original gene in which the target region is replaced by an adapter sequence. This adapter sequence is designed with flanking SapI recognition sequences in order to allow the subcloning of the target region containing the randomly inserted transposon back inside its original gene. This last step results in the generation of the transposon insertion library focused on the region of interest, upon which the cloning steps of TRIAD leading to the generation of InDel libraries (Figure 1) can be performed.

To demonstrate this targeted approach of TRIAD, two deletion (-3 and -6 bp) and one insertion (+3 bp) libraries were generated in the sequence encoding *wt*PTE's active site loop 7 (L7), which has shown to be crucial for the specificity of *wt*PTE and homologous lactonases in previous rational InDel mutagenesis studies [2, 3]. In vitro transposition reactions were performed on a vector containing the L7 - encoding DNA sequence (from Leu252 to Gln278; 81 bp) flanked by SapI restriction sites (Supplementary Figure S13). After isolation by SapI digestion, the resulting L7 TransDel and TransIns insertion libraries were then subcloned into a plasmid containing a modified wtPTE gene with a SapI-adapter instead of L7. This additional cloning step enabled to recreate a full wtPTE gene with TransDel or TransIns randomly inserted - in theory - at all 81 positions within L7. After application of the further steps of TRIAD, libraries with insertions and deletions limited to L7 only were generated. Intermediate and final library transformation steps yielded diversities of $>10^6$ variants, practically oversampling by $>10^5$-fold the theoretical diversity of the libraries (81 possible transposon insertion sites in L7). Sequence analysis of randomly chosen variants revealed the distribution of codons deleted in L7. Whilst there is good coverage of the target sequence (~70% of residues are deleted at least once), there is a bias of deletions toward certain residues, especially Leu262, Leu272 and the residues neighbouring them (Supplementary Table S12). These biases are likely caused by preferential transposon insertions at specific points along the DNA sequence encoding Loop 7.

##### 1.2. Detailed consideration of theoretical diversity and InDel redundancy

The theoretical diversity (*i.e.*, the total number of possible variants) accessible *via* such modifications will depend both on the type of InDel that is introduced and on the target sequence (Supplementary Figure 5). For instance, as deletions can occur once at each position of the target DNA sequence, the maximal possible theoretical diversity of deletion libraries is identical to the number of nucleotides (Supplementary Figure S5A), *e.g.*, ~1,000 possible deletion variants for a ~1 kbp target sequence (*e.g.*, *wt*PTE). By contrast, since TRIAD inserts degenerate nucleotide triplets (*e.g.*, $[NNN]_{1,\ 2\ or\ 3}$ corresponding to 64 (=$4^3$), 4,096 (=$4^6$) and 262,144 (=$4^9$) possible sequences, respectively), the number of possible insertion variants will depend both on the length of the target sequence and the size of the insertions. The maximal possible theoretical diversity for insertion libraries generated from *wt*PTE is $6.4\times10^4$ (=$64\times10^3$), ~$4.1\times10^6$ (=$64^2\times10^3$), and ~$2.6\times10^8$ (=$64^3\times10^3$), corresponding to +3, +6 and +9 bp insertions, respectively (Supplementary Figure S5A). Because of potential InDel redundancy depending on the target sequence (*i.e.*, two or more neighbouring InDels can result in the same DNA variant; Supplementary Figure S5B), the theoretical diversities accessible from a given DNA sequence are usually lower (see Supplementary Figure S5C in the specific case of *wt*PTE). Theoretical diversities at the protein level (*i.e.*, the number of protein variants that have the intended InDel length) are further reduced due to codon degeneracy and occurrence of stop codons as a result of certain InDels (Supplementary Figure S5C). Practically, the size of our libraries was limited by transformation efficiency, achieving > $10^6$ variants upon transformation into *E. coli.* Therefore, deletions as well as +3 bp insertions were oversampled such that the library diversity was maintained between transformations, while the diversity of sampled transposition sites was maintained in larger +6 bp and +9 bp insertion libraries, with only a fraction of theoretical library diversity generated from the outset.

##### 1.3. Library quality assessment by Sanger sequencing

In addition to the deep next-generation sequencing described in the main text, the accuracy of the TRIAD approach (specifically the number of intended in-frame InDels, unwanted frameshifts and incidental mutations) was also assessed using Sanger sequencing to give the reader a picture how such an 'everyday analysis' of a handful of individual *wt*PTE InDel variants would fare. To this end around 20 colonies from each naïve InDel library of individual *wt*PTE variants were randomly picked (after the final transformation step into *E. coli*) and 121 variants in total sequenced (Supplementary Tables S1-3). All the sequenced variants displayed only a single modification resulting from the initial transposon insertion and 90 among them (74%; corresponding to 89 unique InDels) showed anticipated in-frame InDel mutations (86% of the deletion variants; 61% of the insertion variants; Supplementary Tables S1-2). Most in-frame InDels were observed only once and were distributed throughout the *wt*PTE sequence (Supplementary Table S3). No frameshift was observed among sequenced variants from the -3 bp library, which is generated without shuttle cloning steps in contrast to the other libraries. Frameshifts were more frequent among variants from the +3, +6 and +9 bp insertion libraries (~40% of the sequenced variants). Higher frameshift frequency in insertion libraries may be due to exonuclease over-digestion by the Klenow fragment of DNA polymerase I, which removes 3' overhangs left by AcuI digestion (Figure 2B). Note that no incidental additional base pair point mutations located elsewhere in the variants' sequence (*i.e.*, at positions different to that of the initial transposon insertion sites) and resulting from the TRIAD cloning process were detected. On the protein level, TRIAD may generate a secondary point substitution contiguous to the introduced InDel [4], depending on the point of insertion of TransDel or TransIns in the reading frame of the target sequence. This occurs when the InDel is not inserted at previous codon boundaries (statistically in two of three cases, although not all such events lead to amino acid substitution). As a result, 22% of the InDels observed in individual *wt*PTE variants exhibited such an adjacent substitution.

We conclude that the accuracy of the TRIAD procedure can be assessed based on a small number of sequences (n = 121, giving 89 unique in-frame InDels), to provide a quality control step informing in TRIAD library synthesis that is a representative measure of the distribution of InDels over the target sequence and assess the coverage afforded by the initial transposition step prior to diversification *via* InDel mutagenesis (Table 1), in lieu of a deep next-generation sequencing approach.

#### 2. SUPPLEMENTARY METHODS

##### 2.1. Design, construction and preparation of transposons and cloning cassettes

DNA sequences corresponding to the TransDel transposon (Supplementary Figure S2A) and the Del2 cassette (previously dubbed Insertion Replacement Cassette in [5]) were synthesized and cloned into pUC57 (Supplementary Figure S2D) at the EcoRV site (GenScript, NJ, USA). Cloning strategies involving double stranded oligonucleotide adapters (Supplementary Table S14) were used to generate pUC57-TransIns (Supplementary Figure S2A) from pUC57-TransDel, and pUC57-Del3 (Supplementary Figure S2B), -Ins1, -Ins2 and -Ins3 (Supplementary Figure S2C) from pUC57-Del2. For all adapter cloning experiments, each pair of custom phosphorylated oligonucleotides (100 µM in 50mM Tris-HCl pH 8.0, 100 mM NaCl, 1mM EDTA) were mixed to a final concentration of 50 µM and annealed in a PCR thermocycler ((1) 2 min at 95°C, (2) 10 min at 52°C and (3) hold at 4°C). The resulting adapters were then ligated to a final concentration of 125 nM into their target plasmid (50-100 ng). The ligation products were then transformed into electrocompetent *E. coli* E. cloni® 10G cells. Plasmid pUC57-TransIns was generated by inserting the TransIns adapter in pUC57-TransDel at EcoRI/SpeI sites. Plasmid pUC57-Del3 was obtained by inserting the Del3 adapter in pUC57-Del2 at EcoRI/SpeI sites. Plasmids pUC57-Ins1, -Ins2 and -Ins3 correspond to libraries of inserts of one, two and three nucleotide triplets, respectively. First, an intermediate plasmid, dubbed pUC57-Ins, was obtained by inserting the Ins adapter in pUC57-Del2 at EcoRI/SpeI sites. Adapters Ins1, Ins2 and Ins3 were then inserted in pUC57-Ins at NcoI/HindIII sites to generate separate plasmid libraries corresponding to pUC57-Ins1, -Ins2 and -Ins3, respectively. In this last step, each DNA library was extracted from around $10^7$ *E. coli* Ecloni® 10G transforming colonies.

##### 2.2. Design and assembly of pID vectors

Two expression vectors, dubbed pID-T7 (expression under the control of T7 promoter) and pID-Tet (expression under the control of Tet promoter), were specifically designed for the generation of InDel libraries following the TRIAD approach. These vectors do not contain any MlyI, AcuI and NotI restriction sites in their sequence and were assembled from three different modules (for origin of replication, ampicillin resistance (AmpR) selection and expression/cloning) separated by three restriction sites, AflII, AatII and SpeI (Supplementary Figure S4).

***Origin of replication module.*** Two successive site-directed saturation mutagenesis experiments (using primer pairs Ori-MlyI and Ori-AcuI; Supplementary Table S14) were performed to remove recognition sites for MlyI and AcuI in the origin of replication (*ori*) of pUC19 used as starting template. Successful removal of the recognition sequences was confirmed by the absence of restriction digest product with the corresponding enzyme. The final *ori* variant (*i.e.*, with no MlyI and AcuI) was then amplified with primers Ori-AflII and Ori-SpeI (Supplementary Table S14), yielding the origin of replication module (framed by AflII and SpeI) for the pID vectors.

***Ampicillin resistance selection and T7 expression modules.*** The sequences corresponding to the T7 expression (Supplementary Figure S4B) and AmpR cassettes (Supplementary Figure S4D) were synthesized by GenScript (NJ, USA). Position T8 of the T7 promoter was mutated to C to remove the MlyI site present in the natural promoter [6]. Silent mutations were introduced in the AmpR sequence to remove recognition sites for AcuI and FokI.

***Assembly of pID-T7.*** The DNA cassettes corresponding to the modified *ori* (AflII /SpeI), AmpR (AflII /AatII) and the T7 expression module (AatII/SpeI) were isolated by double digestion with their corresponding restriction enzymes and agarose gel purification. A ligation reaction with 50 ng of each DNA fragment was then performed using T4 DNA ligase (Fermentas) overnight at 18˚C. After purification, the ligation products were transformed into electrocompetent *E. coli* Ecloni® 10G cells subsequently plated on LB-agar supplemented with 100 µg/mL ampicillin. The pID-T7 constructs extracted from the resulting transforming colonies were confirmed by restriction digestion profile and sequencing.

***Generation and assembly of pID-Tet.*** TetR (encoding the Tet repressor) was amplified from pASK-IBA5plus (IBA Lifesciences) with primers TetR-F and TetR-B (Supplementary Table S14). AmpR was amplified from pID-T7 with primers mTEM1-F and mTEM1-B. The SpeI/AatII module for pID-Tet containing the AmpR-TetR operon was obtained by overlap PCR of these two products with primers mTEM1-F and TetR-B and subsequently inserted into pID-T7 at SpeI/AatII sites (replacing the AmpR cassette) to yield pID-T7-TetR. The Tet promoter sequence was amplified from pASK-IBA5plus with primers TetProm-F and TetProm-B and inserted into pID-T7-TetR at the AflII/NdeI sites (replacing the T7 promoter), yielding plasmid pID-Tet.

##### 2.3. NGS: Reference sequence used in the alignment for NGS analysis

>*wt*PTE
ATGGCCAGATGATTAATTCCTAATTTTTGTTGACACTCTATCATTGATAGAGTTATTTTACC
ACTCCCTATCAGTGATAGAGAAAAGTGAAATGAATAGTTCGACAAAAATCTAGAAATAATT
TTGTTTAACTTTAAGAAGGAGATATACATATGGCTAGCTGGAGCCACCCGCAGTTCGAAA
AAGGCGCCGGATCCTCC*ATGGGCGATCGGATCAATACCGTGCGCGGTCCTATCACAAT*
*CTCCGAGGCGGGTTTCACACTAACCCACGAGCACATCTGCGGCAGCTCGGCAGGATTC*
*TTGCGTGCTTGGCCGGAGTTCTTCGGTAGCCGCAAAGCTCTAGCGGAAAAGGCTGTGA*
*GAGGATTGCGCCGCGCCAGAGCGGCTGGCGTGCGAACGATTGTCGATGTGTCGACTTT*
*CGATCTCGGTCGCGACGTTAGTTTATTGGCCGAGGTTTCGCGGGCTGCCGACGTTCATA*
*TCGTGGCGGCGACCGGCTTGTGGCTCGACCCGCCACTTTCGATGCGATTGAGGAGTGT*
*AGAGGAACTCACACAGTTCTTCCTGCGTGAGATTCAATATGGCATCGAAGACACCGGAA*
*TTAGGGCGGGCATTATCAAGGTCGCGACCACAGGCAAGGTGACCCCCTTTCAGGAGTTA*
*GTGTTAAGGGCAGCTGCCCGGGCCAGCTTGGCCACCGGTGTTCCGGTAACCACTCACA*
*CGGCAGCAAGTCAGCGCGGTGGTGAGCAACAAGCCGCCATTTTTGAATCCGAGGGCTT*
*GAGCCCCTCACGGGTTTGTATTGGCCACAGCGATGATACTGACGATTTGAGCTATCTCA*
*CCGCCCTCGCTGCGCGCGGATACCTCATCGGTCTAGACCATATTCCGCACAGTGCGATT*
*GGTCTAGAAGATAATGCGAGTGCATCAGCCCTCCTGGGTATTCGTTCGTGGCAAACACG*
*GGCTCTCTTGATCAAGGCGCTCATCGACCAAGGCTACATGAAACAAATCCTCGTTTCGAA*
*TGACTGGCTGTTCGGGTTTTCGAGCTATGTCACCAACATCATGGACGTGATGGATAGCG*
*TGAACCCCGACGGAATGGCCTTCATTCCACTGAGAGTGATCCCATTCCTACGAGAGAAG*
*GGTATTCCACAGGAAACGCTGGCAGGCATCACTGTGACTAACCCGGCGCGGTTCTTGTC*
*ACCGACCTTGCGGGCGTCATGA*AGCTTGCTGCGGCACTCGAGCACCACCACCACCACC

ACTGAGATCCGGCTGCTAACAAAGCCCGAAAGGAAGCTGAGTTGGCTGCTGCCACCGC
TGA

The reference sequence contains the *wt*PTE gene (in italics) flanked by plasmid sequence (underlined). This longer sequence was used to obtain sufficient coverage at the ends of the gene.

##### 2.4. NGS Step 1: Raw data processing

The processing of Illumina sequencing data shown here was performed using computational resources provided by University of Cambridge High Performance Computing (CSD3), but it can also be done on a personal computer. All scripts are available at https://github.com/fhlab/TRIAD.

The first part of analysis is done by the script *count.sh*. Briefly, the process consists of:

1. Assembly of paired-end reads into a single, longer read where possible, using PEAR v. 0.9.10 [7]. Through inspection of sequencing quality in FASTQ files and monitoring of assembly statistics, the options chosen were:

--keep-original --min-overlap 5 --min-assembly-length 0 --quality-threshold 15 --max-uncalled-base 0.01

2. Create an index for the reference with Bowtie2 v.2.3.4 [8] and map both assembled and unassembled FASTQ reads, then sort resulting SAM files with samtools v.1.9 [9] to obtain the sequencing depth.

At this point, 95% of the reads aligned to reference sequence.

3. Based on tags in the SAM file, extract well-mapped reads and of those only keep the reads that contain mutations. Since this step detects any difference from reference, it will contain all reads with InDels as well as reads containing single point substitutions from sequencing errors. Hence, the number of substitutions in the final statistics is over-represented.
4. Since accurate identification of InDel position is essential for analysing transposon sequence preference, we use the deterministic Needleman-Wunsch algorithm to obtain the most accurate possible global alignment of the read to reference. Although using the alignment in the SAM file directly is faster, accepting a longer processing time at this stage is an acceptable trade-off to obtain accurate statistics of the library composition. The alignment was done with the Emboss 6.6.0 [10] implementation *needleall*, which compares many sequences to one.

The standard options for alignment were modified to gap open penalty 15 and gap extend penalty 0.5, in order to accurately identify long (9 bp or more) InDels. The default gap open penalty (10) tends to split long InDels into several short InDels separated by one or two nucleotides.

Alternatively, the data can also be processed on a personal computer with the following modification: once reads are extracted from the SAM file, they should be filtered first for those that contain mutations. This reduces the size of resulting fasta and alignment files, which can otherwise exceed >10 GB. Development and testing of the scripts were done using this method on Linux Mint 18 in a virtual machine with two processor cores and 3 GB RAM. The corresponding code is available in *count_PC.sh.*

##### 2.5. NGS Step 2: Convert alignment to variant counts and generate statistics

The following analysis is done by script *PTE_composition.py* implemented in Python 3 with the following options:

--reference full_fragment.fa (sequence given in above in Supplementary Methods 2.3, plus flanking sequence from the plasmid)
--start_offset 200 (to ignore the preceding plasmid sequence)
--end_tail 97

1. Read in all FASTA multisequence alignment generated by previous step.
2. Create a dictionary containing all associated information: reference name, library name (intended as functional activity fraction or in this case, a multiplexed library), sequencing depth, change in DNA/protein terms, relevant counts. The information is nested with DNA variants nested under relevant protein variants, since multiple DNA variants can result in the same protein mutation. The variant information is stored both in internal format with a functional description (substitution/insertion/deletion/frameshift, used to generate statistics) and according to Human Genome Variation Standard.
3. Scan each pair of sequences (reference + aligned read) from the alignment, detect the mutation, translate to protein and add to dictionary.

Once the count dictionary is complete, it can be used to infer the following:

- Number of mutations per position (DNA or protein)
- Transposon consensus sequence for preferred insertion site
- Composition of insertions
- How many expected deletions/insertions (depending on the library) per DNA position are present

The data analysis, code to infer statistics and resulting figures are available in *TRIAD_composition_figures.ipynb*.

#### 2.6. Point mutations

For reasons of computational efficiency, this pipeline focuses on reads that deviate from the reference. This difference may be a genuine mutation, a PCR error or a sequencing error. The resulting counts are therefore artificially enriched for many variants with a single nucleotide substitution, which each appears only once or perhaps twice. In Sanger sequencing (raw data not shown) we do not observe this kind of 'background noise', suggesting it should be disregarded in the NGS dataset. To corroborate this conclusion, we estimate the true number of point substitutions in the library by calculating the background substitution frequency from reads that align outside *wt*PTE, in the plasmid backbone, where no mutations were deliberately introduced during library construction. Such sequencing artefacts (with a single base pair substitution) occur in 3-4% of all reads, which corresponds to the error rate in the Illumina MiSeq NGS technology. An exact estimate is difficult due to a relatively low number of reads that align outside the fragment, as well as sequence dependence of polymerase errors – such that the error rate may be different in and out of the gene. Therefore, in the calculation of the proportion of frameshifts in the library, point mutations were simply counted as wild type (thus removing this 'noise').

#### 2.7. InDel redundancy

While pure substitution mutations can be placed very accurately, correct placement of InDels can be inherently ambiguous depending on the sequence context (as discussed in 1.2). InDels show some inherent redundancy, where distinct transposition and insertion / deletion events result in identical final sequence (Supplementary Figure S5B). For example, in the original sequence …nnGCTACTnn…, -3 bp deletions starting at position 2 (G---CT) and at position 3 (GC---T) result in the same final sequence: nnGCTnn (the remaining sequence context is abbreviated with *n*). The Needleman-Wunsch algorithm consistently (though arbitrarily) assigns this sequence to a deletion at position 2, such that any deletions originating from position 3 *cannot* be directly observed.

Implications:

- The raw counts that describe how many times a mutation was observed at which position, must be adjusted for this ambiguity, if we wish to infer the transposition sequence preference of the Mu transposon. The ambiguity can be partially corrected for deletions by generating a set of baseline reads that contain a -3 bp deletion at every *wt*PTE position and processing them in the same way as sequencing reads. Knowledge of these baseline counts allows us to split the observed counts in the -3 bp library across all originating positions. Data processed in this way was the basis of the frequency plot in Figure 3A.
- The diversity of mutations that can be *observed* is reduced compared to the maximal theoretical library diversity (Supplementary Figure S5). For example, in the -3 bp deletion library the theoretical diversity is one deletion per bp of gene length or 1000 variants for *wt*PTE, but the observable diversity due to ambiguity is 748 variants

(based on the particular sequence of the *wt*PTE gene). Hence, the -3 bp deletions actually observed by deep sequencing at 639 positions reflect 85% (=639/748) coverage of all possible variants, not 64% (=639/1000). Similarly, the maximum diversity of insertion libraries is less than maximal theoretical diversity at DNA level is 64 variants / triplet inserted / bp gene length. For all deletion and the +3 bp libraries, we calculated the theoretical diversity by computationally generating a perfect library (with script *baseline.py*), which contains every variant once (ie. 1 deletion of each length at each position, 1 insertion of each of the 64 codons at each positions), then processed this library in the same way as the NGS dataset. This shows the theoretical diversity in deletion libraries is ~0.75 deletion / bp gene length in *wt*PTE, while in the +3 bp library it is on average 46.22 variants / bp gene length.

In this manuscript, we focus on discussing the *observed* variants, rather than the number of variants inferred, so the InDel redundancy is generally not corrected. The exception is the discussion of Mu transposon sequence preference (Figure 3A).

#### 3. SUPPLEMENTARY FIGURES

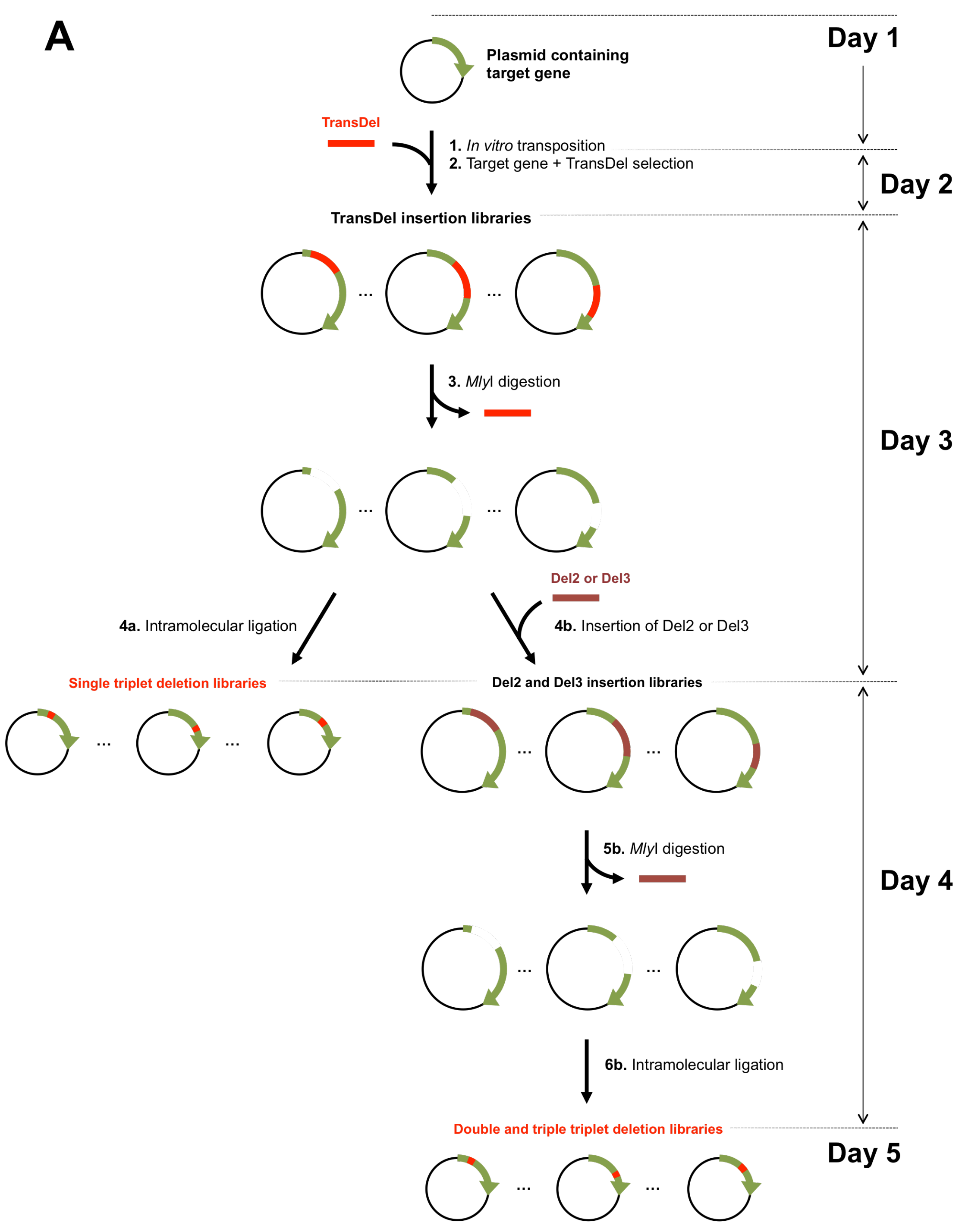

**Supplementary Figure S1** (Continued on next page, legend follows).

(Figure S1 continued)

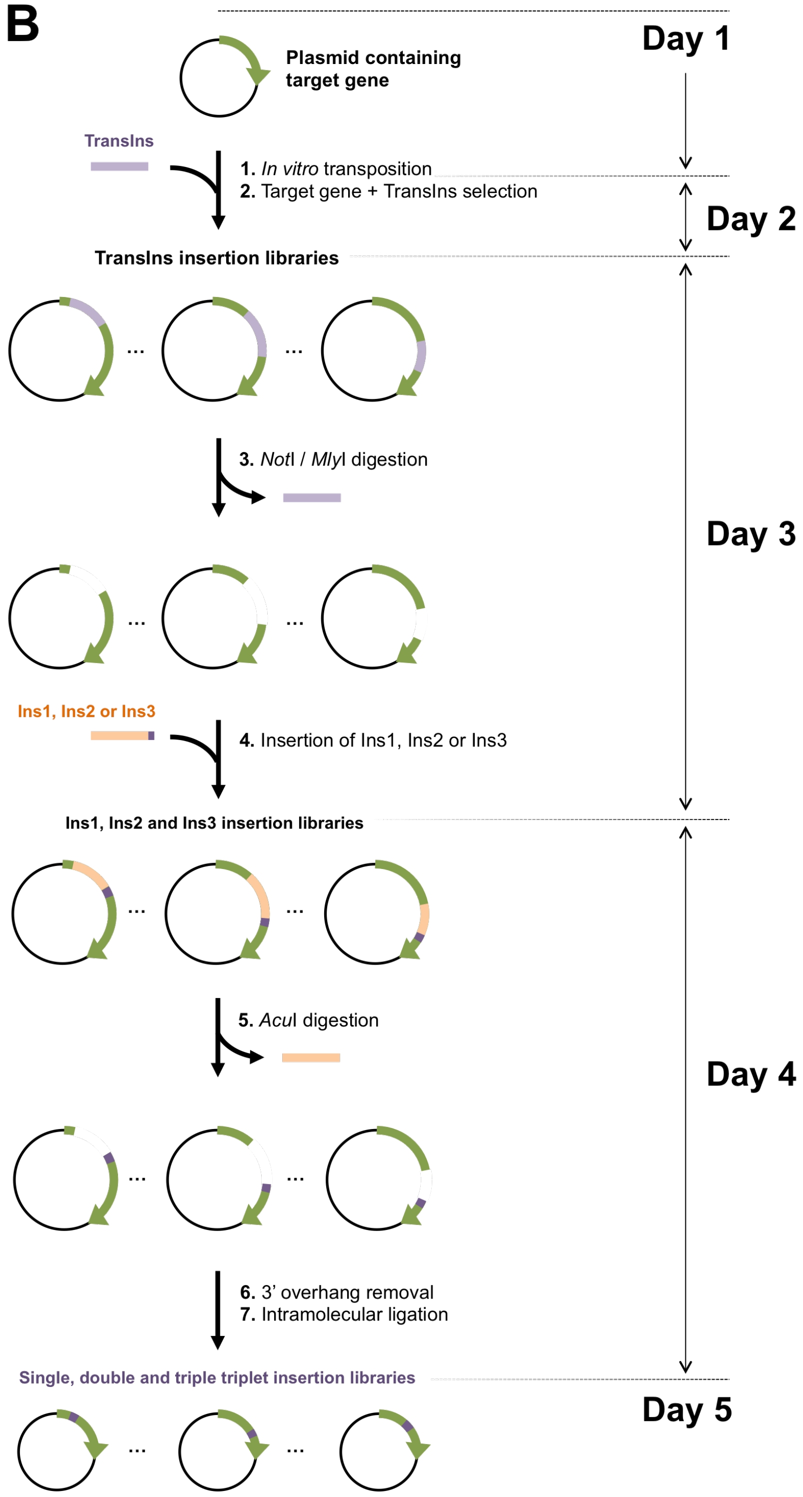

**Supplementary Figure S1** (Continued on next page, legend follows).

(Figure S1 continued)

**Supplementary Figure S1. Schematic outline and timeline of the procedure for the generation of random InDel libraries**

**A.** Generation of deletion libraries.

Step 1: The TransDel insertion library is generated by *in vitro* transposition of the engineered transposon TransDel into the plasmid containing the target gene followed by the subcloning of the fragment comprising the target gene and the transposon into a fresh plasmid.

Step 2: MlyI digestion removes TransDel together with 3 bp of the original target gene and generates a single break per target gene variant.

Step 3a: Intramolecular ligation results in the reformation of the target gene minus 3 bp, yielding a library of single variants with a deletion of 1 triplet [4].

Step 3b: DNA cassettes dubbed Del2 and Del3 are then inserted between the break in the target gene to generate Del2 and Del3 insertion libraries.

Step 4b: MlyI digestion removes Del2 and Del3 together with 3 and 6 additional bp of the original GOI, respectively.

Step 5b: Intramolecular ligation results in the reformation of the target gene minus 6 and 9 bp, yielding libraries of single variants with a deletion of 2 and 3 triplets, respectively. Red vertical lines indicate deletions.

**B.** Generation of insertion libraries.

Step 1: The TransIns insertion library is generated by *in vitro* transposition of the engineered transposon into the target gene.

Step 2: digestion by NotI and MlyI removes TransIns.

Step 3: DNA cassettes dubbed Ins1, Ins 3 and Ins3 (with respectively 1, 2 and 3 randomized bp triplets at one of their extremities; indicated in blue) are then inserted between the break in the target gene to generate the corresponding Ins1, Ins2 and Ins3 insertion libraries.

Step 4: AcuI digestion and 5'end digestion by the Klenow fragment remove the cassettes, leaving the randomized triplet(s) in the original target gene.

Step 5: Intramolecular ligation results in the reformation of the target gene plus 3, 6 and 9 random bp, yielding libraries of single variants with an insertion of 1, 2 and 3 triplets, respectively. Purple vertical lines indicate insertions.

#### (A) Mu transposons TransDel and TransIns

BglII **MlyI** **MuA binding site (R1/R2)**

AGATCT**GACTC**GGCGCACGAAAAACGCGAAAGCGTTTCACGATAAATGCGAAAACTTTTCCCATGCATGGGAATAAATA

CCTGTGACGGAAGATCACTTCGCAGAATAAATAAATCCTGGTGTCCCTGTTGATACCGGGAAGCCCTGGGCCAACTTTT

**Cat promoter**

GGCGAAAATGAGACGTTGATCGGCACGTAAGAGGTTCCAACTTTCACCATAATGAAATAAGATCACTACCGGGCGTATT

**Chloramphenicol acetyltransferase**

TTTTGAGTTGTCGAGATTTTCAGGAGCTAAGGAAGCTAAAATGGAGAAAAAAATCACTGGATATACCACCGTTGATATA

**(CamR)**

TCCCAATGGCATCGTAAAGAACATTTTGAGGCATTTCAGTCAGTTGCTCAATGTACCTATAACCAGACCGTTCAGCTGG

ATATTACGGCCTTTTTAAAGACCGTAAAGAAAAATAAGCACAAGTTTTATCCGGCCTTTATTCACATTCTTGCCCGCCT

GATGAATGCTCATCCGGAATTACGTATGGCAATGAAAGACGGTGAGCTGGTGATATGGGATAGTGTTCACCCTTGTTAC

ACCGTTTTCCATGAGCAAACTGAAACGTTTTCATCGCTCTGGAGTGAATACCACGACGATTTCCGGCAGTTTCTACACA

TATATTCGCAAGATGTGGCGTGTTACGGTGAAAACCTGGCCTATTTCCCTAAAGGGTTTATTGAGAATATGTTTTTCGT

CTCAGCCAATCCCTGGGTGAGTTTCACCAGTTTTGATTTAAACGTGGCCAATATGGACAACTTCTTCGCCCCCGTTTTC

ACTATGGGCAAATATTATACGCAAGGCGACAAGGTGCTGATGCCGCTGGCGATTCAGGTTCATCATGCCGTTTGTGATG

GCTTCCATGTCGGCAGAATGCTTAATGAATTACAACAGTACTGCGATGAGTGGCAGGGCGGGGCGTAATGATATCGAGC

λ **$t_0$ transcription terminator**

TCGCTTTCTGTTGATAGATCCAGTAATGACCTCAGAACTCCATCTGGATTTGTTCAGAACGCTCGGTTGCCGCCGGGCG

SpeI **MuA binding site (R1/R2)**

TTTTTTATTGGTGAGAATCCAAGCACTAGTCGAGATCCGTTTTCGCATTTATCGTGAAACGCTTTCGCGTTTTTCGTGC

**MlyI** BglII

**(TransDel)** GCC**GAGTC**AGATCT

**NotI** BglII

**(TransIns)** **GCGGCCGC**AGATCT

#### (B) Deletion cassettes Del2 and Del3

SmaI **MlyI** EcoRI

**(Del2)** CCCGGGAT**GACTC**CATGG

SmaI **MlyI** EcoRI

**(Del3)** CCCGG**GACTC**CATGG

ACTTCGCAGAATAAATAAATCCTGGTGTCCCTGTTGATACCGGGAAGCCCTGGGCCAACTTTTGGCGAAAATGAGACGT

**Cat promoter**

TGATCGGCACGTAAGAGGTTCCAACTTTCACCATAATGAAATAAGATCACTACCGGGCGTATTTTTTGAGTTGTCGAGA

**Kanamycin nucleotidyltransferase (KanR)**

TTTTCAGGAGCTAAGGAAGCTAAAATGATTGAACAAGATGGATTGCACGCAGGTTCTCCGGCAGCTTGGGTGGAGAGGC

TATTCGGCTATGACTGGGCACAACAGACAATCGGCTGCTCTGATGCCGCCGTGTTCCGGCTGTCAGCGCAGGGGCGCCC

GGTTCTTTTTGTCAAGACCGACCTGTCCGGTGCCCTGAATGAACTGCAAGACGAGGCAGCGCGGCTATCGTGGCTGGCC

ACGACGGGCGTTCCTTGCGCAGCTGTGCTCGACGTTGTCACTGAAGCGGGAAGGGACTGGCTGCTATTGGGCGAAGTGC

CGGGGCAGGATCTCCTGTCATCTCACCTTGCTCCTGCCGAGAAAGTATCCATCATGGCTGATGCAATGCGGCGGCTGCA

TACGCTTGATCCGGCTACCTGCCCATTCGACCACCAAGCGAAACATCGCATCGAGCGAGCACGTACTCGGATGGAAGCC

GGTCTTGTCGATCAGGATGATCTGGACGAAGAGCATCAGGGGCTCGCGCCAGCCGAACTGTTCGCCAGGCTCAAGGCGA

GCATGCCCGACGGCGAGGATCTCGTCGTGACCCACGGCGATGCCTGCTTGCCGAATATCATGGTGGAAAATGGCCGCTT

TTCTGGATTCATCGACTGTGGCCGGCTGGGTGTGGCGGACCGCTATCAGGACATAGCGTTGGCTACCCGTGATATTGCT

GAAGAGCTTGGCGGCGAATGGGCTGACCGCTTCCTCGTGCTTTACGGTATCGCCGCTCCCGATTCGCAGCGCATCGCCT

TCTATCGCCTTCTTGACGAGTTCTTCTGATATCGAGCTCGCTTTCTGTTGATAGATCCAGTAATGACCTCAGAACTCCA

λ **$t_0$ transcription terminator** SpeI **MlyI** SmaI

TCTGGATTTGTTCAGAACGCTCGGTTGCCGCCGGGCGTTTTTTATTGGTGAGAATCCAAGCACTAGTC**GAGTC**CCGGG

**Supplementary Figure S2** (Continued on next page, legend follows).

(Figure S2 continued)

#### (C) Insertion cassettes Ins1, Ins2 and Ins3

MlyI Insert AcuI EcoRI

GAGTCAGCGC **(NNN)**$_n$ ATCCATCTCGAGTGGC**CTTCAG**CCATGGACTTCGCAGAATAAATAAATCCTGGTGTCCCTG

**Cat promoter**

TTGATACCGGGAAGCCCTGGGCCAACTTTTGGCGAAAATGAGACGTTGATCGGCACGTAAGAGGTTCCAACTTTCACCA

**Kanamycin**

TAATGAAATAAGATCACTACCGGGCGTATTTTTTGAGTTGTCGAGATTTTCAGGAGCTAAGGAAGCTAAAATGATTGAA

**nucleotidyltransferase (KanR)**

CAAGATGGATTGCACGCAGGTTCTCCGGCAGCTTGGGTGGAGAGGCTATTCGGCTATGACTGGGCACAACAGACAATCG

GCTGCTCTGATGCCGCCGTGTTCCGGCTGTCAGCGCAGGGGCGCCCGGTTCTTTTTGTCAAGACCGACCTGTCCGGTGC

CCTGAATGAACTGCAAGACGAGGCAGCGCGGCTATCGTGGCTGGCCACGACGGGCGTTCCTTGCGCAGCTGTGCTCGAC

GTTGTCACTGAAGCGGGAAGGGACTGGCTGCTATTGGGCGAAGTGCCGGGGCAGGATCTCCTGTCATCTCACCTTGCTC

CTGCCGAGAAAGTATCCATCATGGCTGATGCAATGCGGCGGCTGCATACGCTTGATCCGGCTACCTGCCCATTCGACCA

CCAAGCGAAACATCGCATCGAGCGAGCACGTACTCGGATGGAAGCCGGTCTTGTCGATCAGGATGATCTGGACGAAGAG

CATCAGGGGCTCGCGCCAGCCGAACTGTTCGCCAGGCTCAAGGCGAGCATGCCCGACGGCGAGGATCTCGTCGTGACCC

ACGGCGATGCCTGCTTGCCGAATATCATGGTGGAAAATGGCCGCTTTTCTGGATTCATCGACTGTGGCCGGCTGGGTGT

GGCGGACCGCTATCAGGACATAGCGTTGGCTACCCGTGATATTGCTGAAGAGCTTGGCGGCGAATGGGCTGACCGCTTC

CTCGTGCTTTACGGTATCGCCGCTCCCGATTCGCAGCGCATCGCCTTCTATCGCCTTCTTGACGAGTTCTTCTGATATC

**$\lambda$ $t_0$ transcription**

GAGCTCGCTTTCTGTTGATAGATCCAGTAATGACCTCAGAACTCCATCTGGATTTGTTCAGAACGCTCGGTTGCCGCCG

**terminator** SpeI **AcuI** NotI

GGCGTTTTTTATTGGTGAGAATCCAAGCACTAGTCAGCAGCGA**CTGAAG**ACGGATGCGGCCGC

## (D)

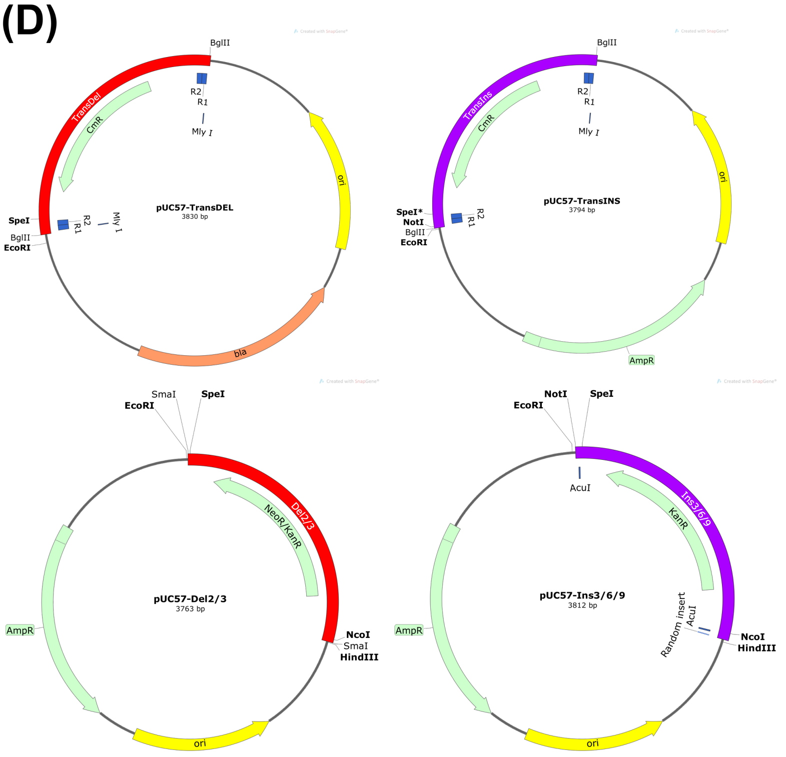

**Supplementary Figure S2** (Continued on next page, legend follows).

(Figure S2 continued)

**Supplementary Figure S2. Engineered transposons and cloning cassettes used in TRIAD.**

Sequences of **(A)** Mu transposons TransDel and TransIns, **(B)** deletion cassettes Del2 and Del3, **(C)** insertion cassettes Ins1, Ins2 and Ins3 (n = 1, 2 or 3 nucleotide triplets), **(D)** Maps of vectors pUC-57-TransDel, -TransIns, -Del2/3, -Ins1/2/3.

**DNA sequence**

NcoI

CC**ATG**GGCGATCGGATCAATACCGTGCGCGGTCCTATCACAATCTCCGAGGCGGGTTTCACACTAACCCACGAGCACATCTGCGGCA
GCTCGGCAGGATTCTTGCGTGCTTGGCCGGAGTTCTTCGGTAGCCGCAAAGCTCTAGCGGAAAAGGCTGTGAGAGGATTGCGCCGCG
CCAGAGCGGCTGGCGTGCGAACGATTGTCGATGTGTCGACTTTCGATCTCGGTCGCGACGTTAGTTTATTGGCCGAGGTTTCGCGGG
CTGCCGACGTTCATATCGTGGCGGCGACCGGCTTGTGGCTCGACCCGCCACTTTCGATGCGATTGAGGAGTGTAGAGGAACTCACAC
AGTTCTTCCTGCGTGAGATTCAATATGGCATCGAAGACACCGGAATTAGGGCGGGCATTATCAAGGTCGCGACCACAGGCAAGGTGA
CCCCCTTTCAGGAGTTAGTGTTAAGGGCAGCTGCCCGGGCCAGCTTGGCCACCGGTGTTCCGGTAACCACTCACACGGCAGCAAGTC
AGCGCGGTGGTGAGCAACAAGCCGCCATTTTTGAATCCGAGGGCTTGAGCCCCTCACGGGTTTGTATTGGCCACAGCGATGATACTG
ACGATTTGAGCTATCTCACCGCCCTCGCTGCGCGCGGATACCTCATCGGTCTAGACCATATTCCGCACAGTGCGATTGGTCTAGAAG
ATAATGCGAGTGCATCAGCCCTCCTGGGTATTCGTTCGTGGCAAACACGGGCTCTCTTGATCAAGGCGCTCATCGACCAAGGCTACA
TGAAACAAATCCTCGTTTCGAATGACTGGCTGTTCGGGTTTTCGAGCTATGTCACCAACATCATGGACGTGATGGATAGCGTGAACC
CCGACGGAATGGCCTTCATTCCACTGAGAGTGATCCCATTCCTACGAGAGAAGGGTATTCCACAGGAAACGCTGGCAGGCATCACTG
TGACTAACCCGGCGCGGTTCTTGTCACCGACCTTGCGGGCGTCA**TGA**AGCTT

HindIII

**Protein sequence**

MASWSHPQFEKGAGSSMG$^{34}$DRINTVRGPITISEAGFTLTHEHICGSSAGFLRAWPEFFGSRKALAEKAVRGLRRARAAGVRTIVDVS
TFDLGRDVSLLAEVSRAADVHIVAATGLWLDPPLSMRLRSVEELTQFFLREIQYGIEDTGIRAGIIKVATTGKVTPFQELVLRAAAR
ASLATGVPVTTHTAASQRGGEQQAAIFESEGLSPSRVCIGHSDDTDDLSYLTALAARGYLIGLDHIPHSAIGLEDNASASALLGIRS
WQTRALLIKALIDQGYMKQILVSNDWLFGFSSYVTNIMDVMDSVNPDGMAFIPLRVIPFLREKGIPQETLAGITVTNPARFLSPTLR
AS$^{365}$

**Supplementary Figure S3: Sequence of the synthetic *wt*PTE gene and its corresponding protein product**

This gene was designed without MlyI, AcuI and NotI restriction sites and cloned into pID-Tet or pET-strep vectors using NcoI and HindIII (underlined). Start and stop codons are shown in bold. The resulting protein (in red) was expressed in fusion with a Strep-tag II peptide (shown in green) at its N-terminus and its sequence corresponds to the one referred to as PTE-R0 in [11, 12] and as *wt*PTE in [5, 13]. Residues were numbered according to PDB 4PCP.

**A**

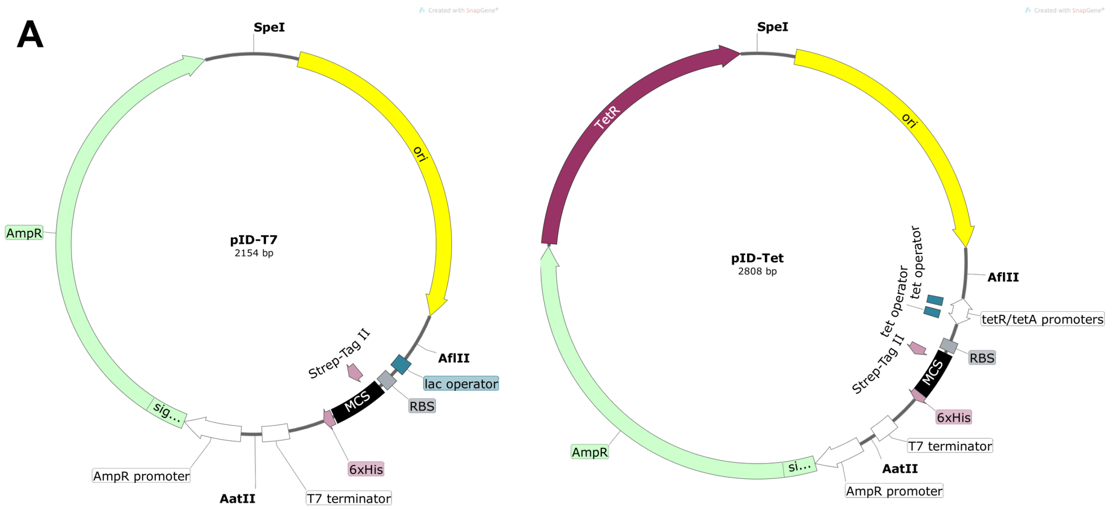

### B Expression cassette of pID-T7

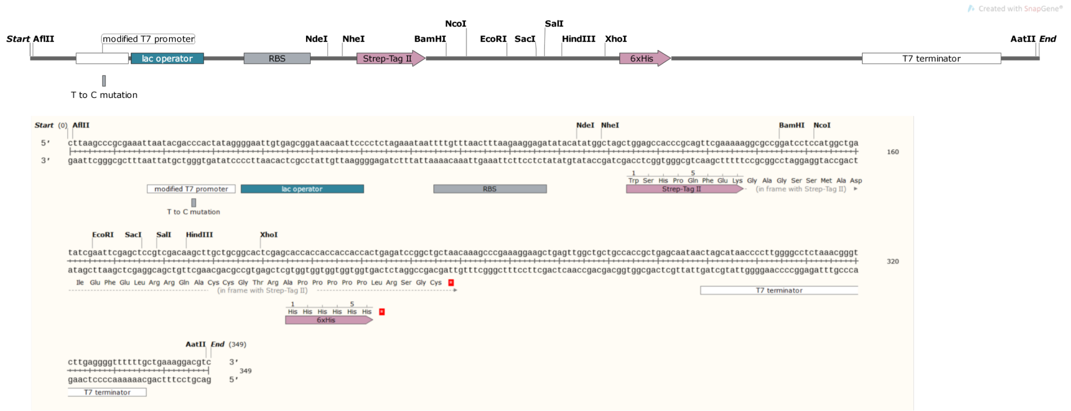

### C Expression cassette of pID-Tet

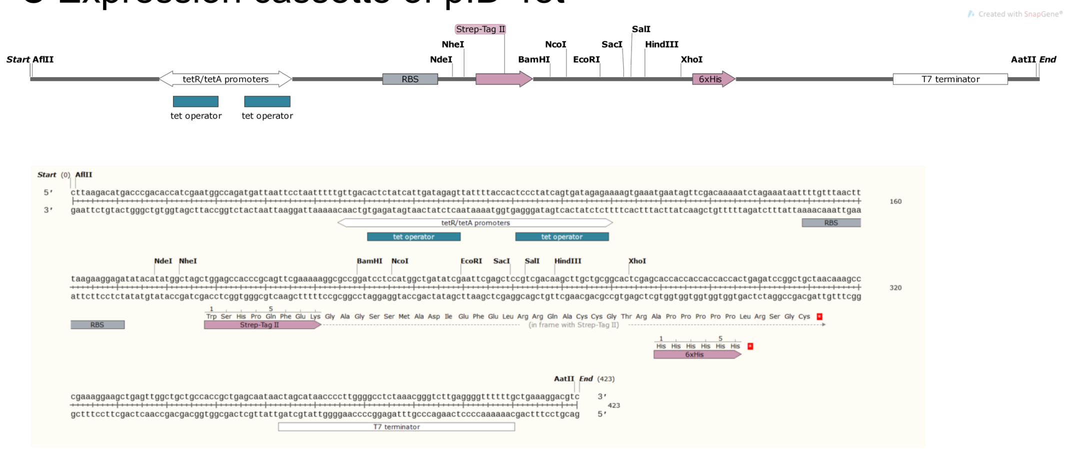

**Supplementary Figure S4** (Continued on next page, legend follows).

(Figure S4 continued)

#### D AmpR cassette for pID vectors

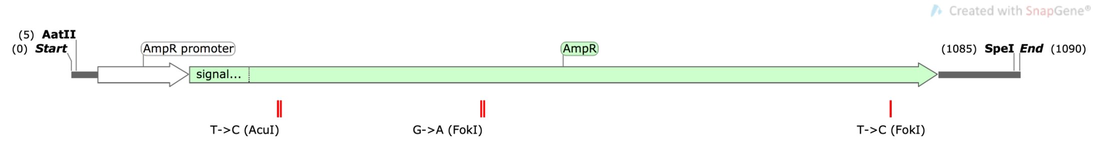

```
AatII                                   AmpR promoter
GACGTCAGGTGGCACTTTTCGGGGAAATGTGCGCGGAACCCCTATTTGTTTATTTTTCTAAATAC

ATTCAAATATGTATCCGCTCATGAGACAATAACCCTGATAAATGCTTCAATAATATTGAAAAAGG
        AmpR
AAGAGTATGAGTATTCAACATTTCCGTGTCGCCCTTATTCCCTTTTTTGCGGCATTTTGCCTTCC
                                  (AcuI) CTGAAG
TGTTTTTGCTCACCCAGAAACGCTGGTGAAAGTAAAAGATGCCGAGGATCAGTTGGGTGCACGAG

TGGGTTACATCGAACTGGATCTCAACAGCGGTAAGATCCTTGAGAGTTTTCGCCCCGAAGAACGT

TTTCCAATGATGAGCACTTTTAAAGTTCTGCTATGTGGCGCGGTATTATCCCGTATTGACGCCGG

GCAAGAGCAACTCGGTCGCCGCATACACTATTCTCAGAATGACTTGGTTGAGTACTCACCAGTCA
         (FokI) GGATG
CAGAAAAGCATCTTACAGACGGCATGACAGTAAGAGAATTATGCAGTGCTGCCATAACCATGAGT

GATAACACTGCGGCCAACTTACTTCTGACAACGATCGGAGGACCGAAGGAGCTAACCGCTTTTTT

GCACAACATGGGGGATCATGTAACTCGCCTTGATCGTTGGGAACCGGAGCTGAATGAAGCCATAC

CAAACGACGAGCGTGACACCACGATGCCTGTAGCAATGGCAACAACGTTGCGCAAACTATTAACT

GGCGAACTACTTACTCTAGCTTCCCGGCAACAATTAATAGACTGGATGGAGGCGGATAAAGTTGC

AGGACCACTTCTGCGCTCGGCCCTTCCGGCTGGCTGGTTTATTGCTGATAAATCTGGAGCCGGTG

AGCGTGGGTCTCGCGGTATCATTGCAGCACTGGGGCCAGATGGTAAGCCCTCCCGTATCGTAGTT
                        (FokI) GGATG
ATCTACACGACGGGCAGTCAGGCAACTATGGACGAACGAAATAGACAGATCGCTGAGATAGGTGC

CTCACTGATTAAGCATTGGTAACTGTCAGACCAAGTTTACTCATATATACTTTAGATTGATTTAA
                                            SpeI
AACTTCATTTTTAATTTAAAAGGATCTAGGTGAAGATCCTTTTTACTAGT
```

**Supplementary Figure S4: Vectors for the generation of InDel variant libraries**

These vectors are designed for the construction of TRIAD libraries and expression of the generated variants (feature not used in the present study). They were designed and assembled (see supplementary methods) from different components separated by three restriction sites: (*i)* ampicillin resistance gene (AmpR; AatII/SpeI) forming an operon with TetR (encoding the tetracycline repressor) in the case of pID-Tet, (*ii*) origin of replication (ori; SpeI/AflII), and (*iii*) expression cassette (AflII/AatII) consisting of a promoter (T7 and Tet for pID-T7 and pID-Tet, respectively), a multiple cloning site (MCS), sequences encoding affinity tags (*i.e.* Strep-tag II and 6xHis-tag) and the T7 terminator sequence. These vectors do not contain any MlyI, AcuI and NotI restriction sites in their sequence. **(A)** Maps of vector pID-T7 and pID-Tet. **(B)** Map and sequence of expression cassette for pID-T7, where position T8 of the T7 promoter was mutated to C to remove MlyI site as in [6]. **(C)** Map and sequence of expression cassette for pID-Tet. **(D)** Map and sequence of AmpR cassette (silent mutations to remove AcuI and FokI sites are indicated).

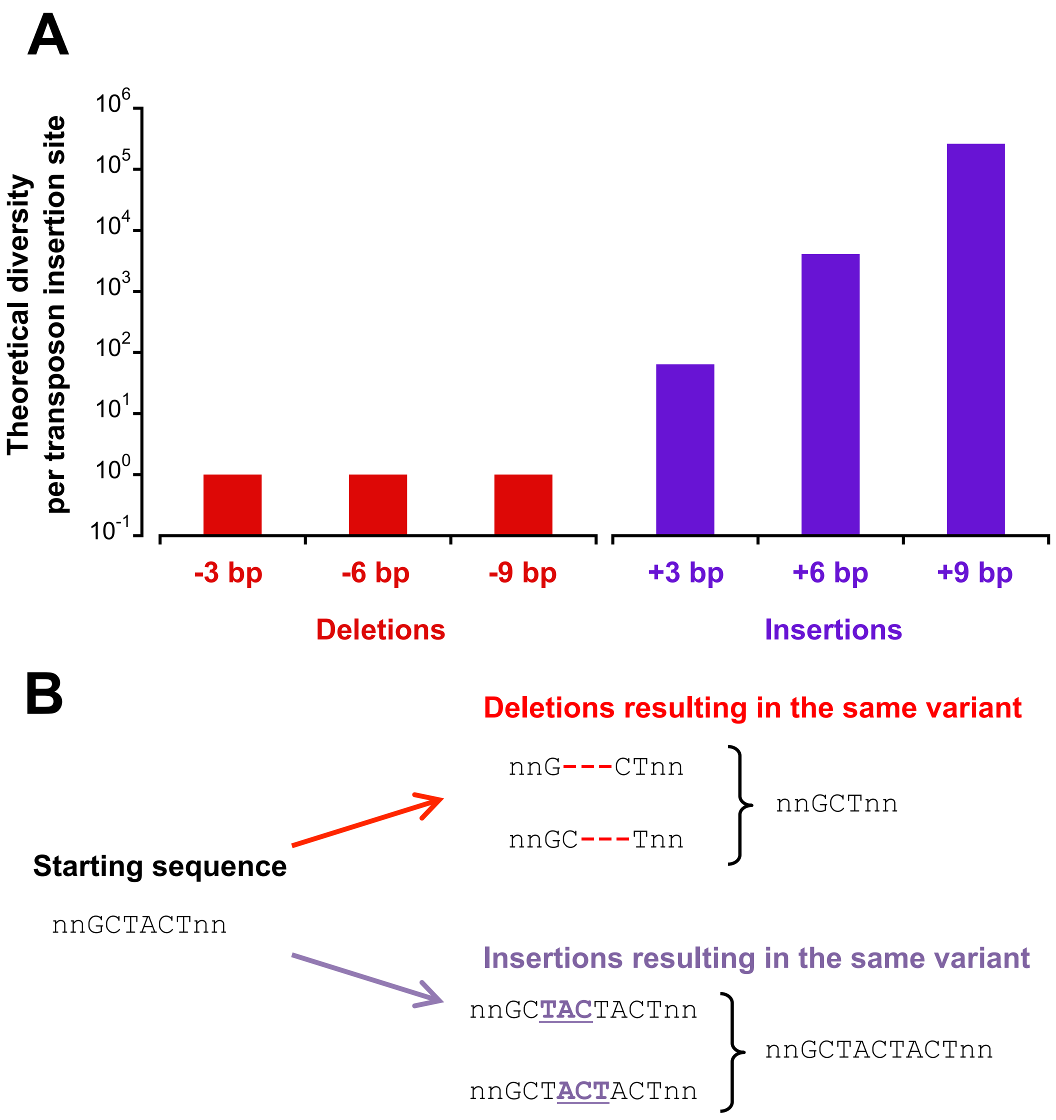

C

| | Deletions | | | Insertions | |
|---|---|---|---|---|---|
| | **-3 bp** | **-6 bp** | **-9 bp** | **+3 bp** | **+6 bp** |
| **Theoretical diversity** | | | | | |
| *at the DNA level* | 748 | 747 | 730 | 46170 | 2844330 |
| *at the protein level* | 590 | 589 | 551 | 13004 | 268159 |

**Supplementary Figure S5: Theoretical diversities of the InDel libraries obtained with TRIAD**

**(A)** This plot shows the theoretical diversity per transposon insertion site within the target gene for each type of InDel generated using TRIAD. Only one deletion, regardless of its length, can occur at a given transposition insertion site. Conversely, insertion diversity is related to the length of the randomized inserted triplet nucleotides ($4^3$, $4^6$ and $4^9$ for one, two and three triplet insertions, respectively). Transposon insertion can occur at each position of the target DNA, providing it does not affect the restriction sites that are necessary in the various subcloning steps of the TRIAD procedure. Therefore, the theoretical diversity for each TRIAD library is obtained by multiplying the values plotted in this figure with the length of the target DNA. In the case of a target gene with the length of *wt*PTE (~1,000 bp), the theoretical diversity for deletion libraries is ~$10^3$ while it is $6.4 \times 10^4$, ~$4.1 \times 10^6$ and ~$2.6 \times 10^8$ for one, two and three triplet insertions, respectively. **(B)** Examples of InDel redundancy in the case of deletions (-3 bp) or insertions (+3 bp) of a triplet resulting in the same DNA variants. **(C)** Theoretical diversities accessible by applying TRIAD to the *wt*PTE gene sequence in TRIAD libraries -3, -6, -9, +3 and +6 bp. The diversities were determined computationally by using the python script *baseline.py* to generate fasta reads with all possible mutations, which were then aligned and counted in the exact same way as the physical libraries.

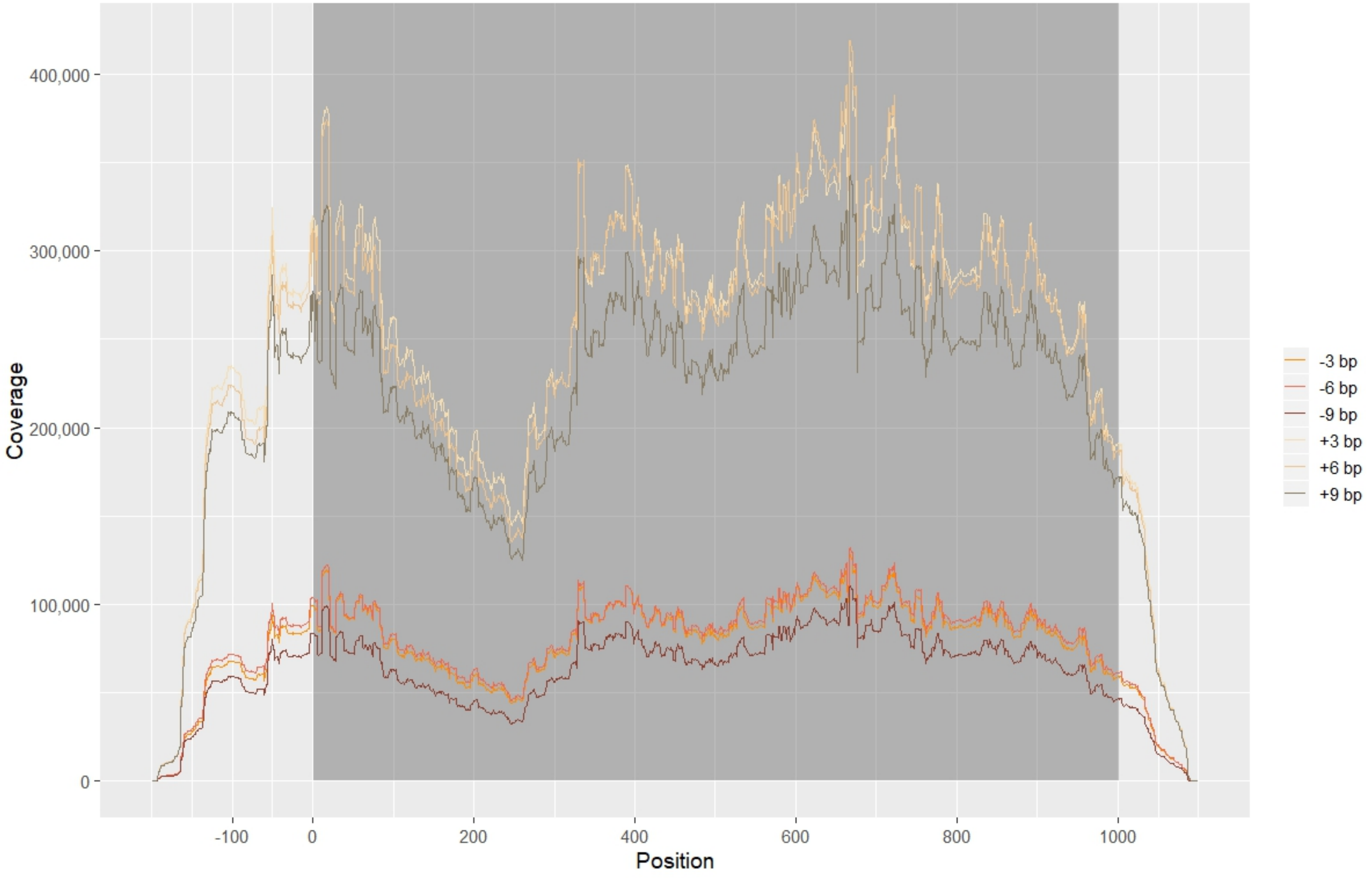

Supplemental Figure S6 (legend next page)

**Supplementary Figure S6: Sequencing coverage in the NGS of the TRIAD libraries of *wt*PTE.**

The final total depth from all assembled and unassembled reads that map to the reference is shown along the entire sequencing fragment. The proportion of reference DNA corresponding to *wt*PTE gene is shown on shaded background. Coverage for insertion libraries is approximately 3× that of deletion libraries, due to higher loading onto the MiSeq flow-cell in order to capture the higher diversity of insertion libraries better. Despite a decrease in coverage around position 200, good coverage is maintained across the entire *wt*PTE gene.

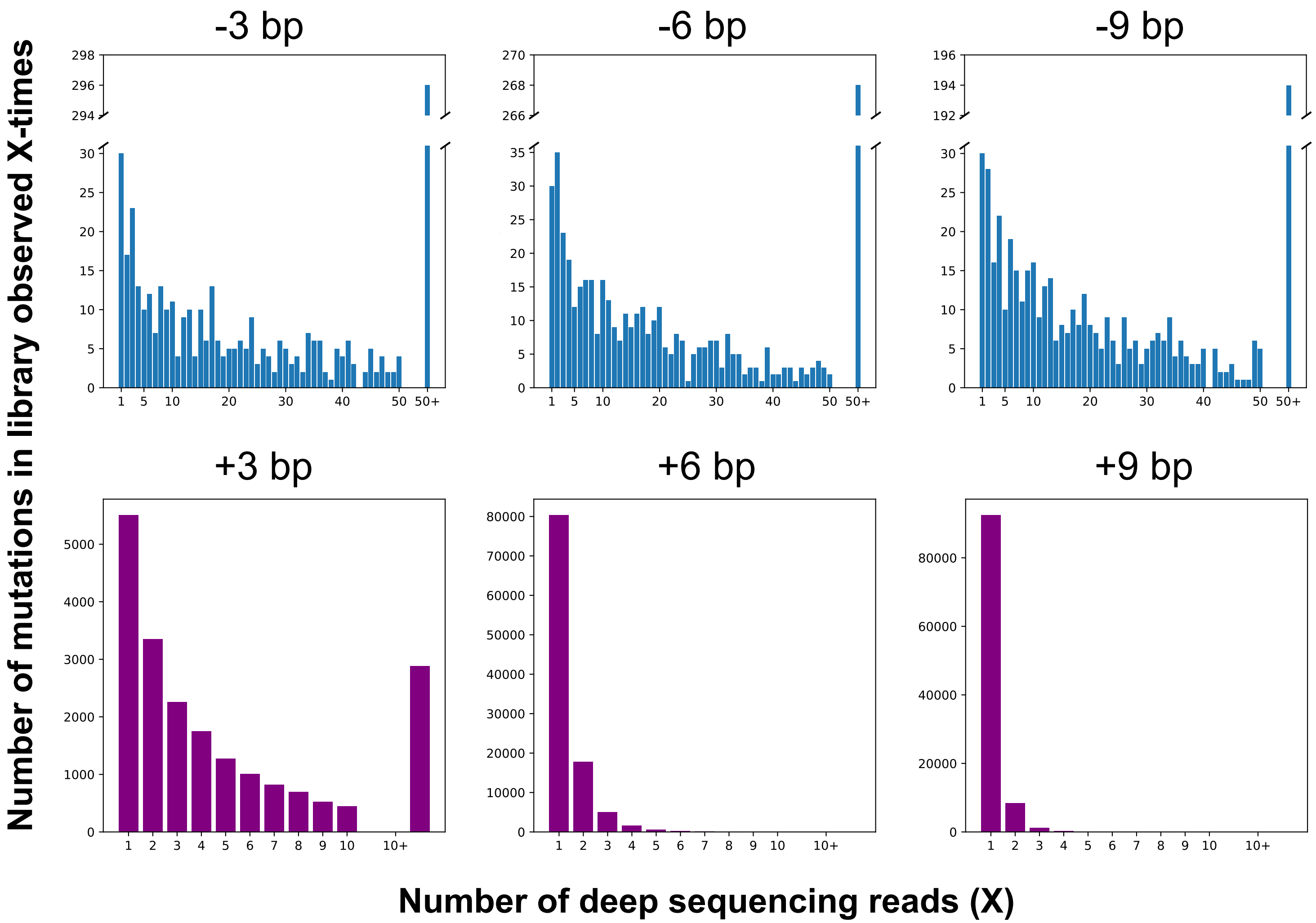

Supplemental Figure S7 (legend next page)

**Supplementary Figure S7: Distribution of observed number of reads per mutation.**

The histograms show how many mutations are observed once, twice, thrice, ten times or more. In deletion libraries we find that most mutations detected are supported by 10-40 observations (each observation is a single read in raw sequencing data).

The bias of transposon site preference results in InDels being observed more often at some positions than other. In the deletion libraries, most mutations are supported by < 50 reads (see Table S6), but there is a long tail generated by positions that are close to the Mu transposon consensus – this is aggregated into one bin in the histograms in this Figure for clarity. Because of the large diversity of insertion libraries, variants are generally observed with lower frequencies (x-axis) compared to the deletion libraries.

# A

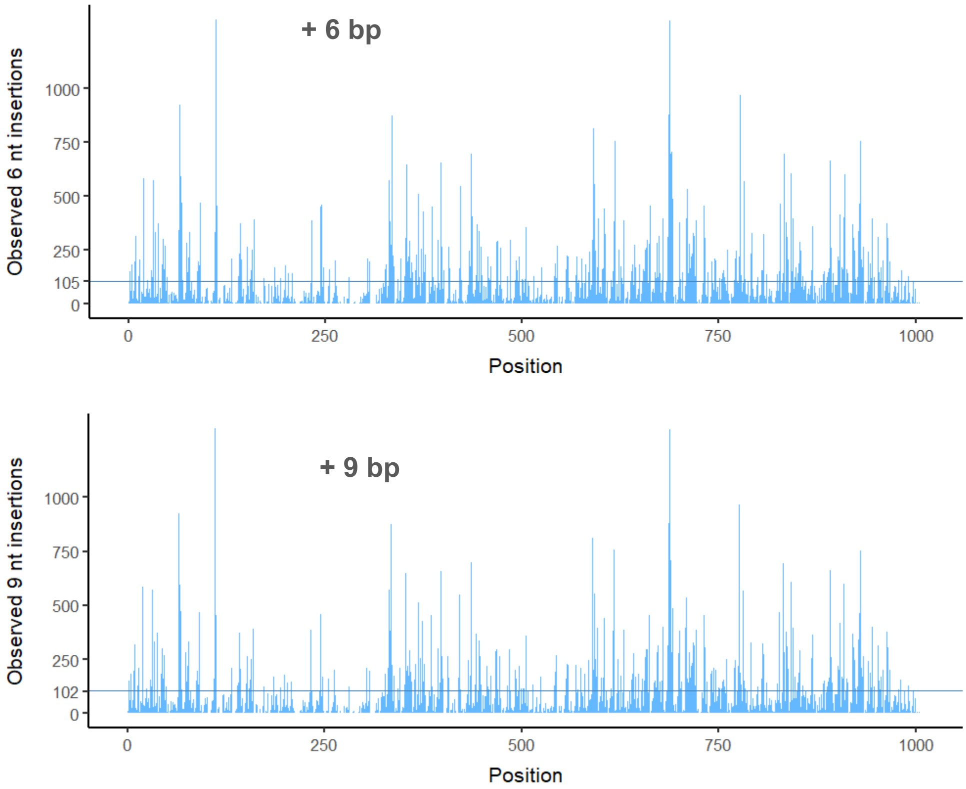

**Supplementary Figure S8** (Continued on next page, legend follows).

(Figure S8 continued)

# B

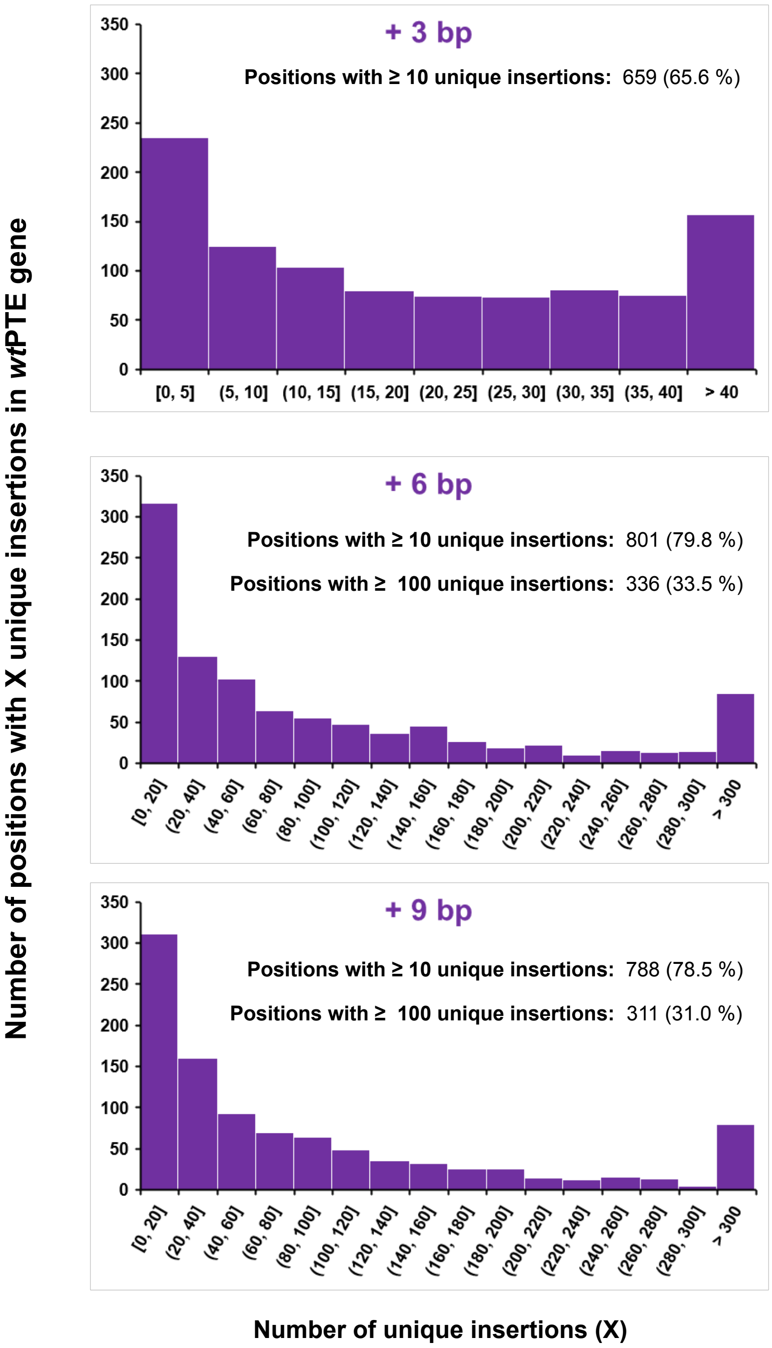

Supplemental Figure S8 (legend next page)

**Supplementary Figure S8: Number of distinct insertions observed per position in *wt*PTE.**

**(A)** Distribution and number of distinct (or unique) insertions per DNA position determined by deep sequencing in +6 bp and + 9 bp bp libraries., compared to the mean per position.In analogy to Figure 3C in the main text, this figure shows the number of observed insertions at each position where mutations are observed. The possible diversity of +6 bp and +9 bp insertions is much higher than for +3 bp library, which results in more pronounced high diversity “spiked” at positions where transposon insertion is favoured. The horizontal line shows the mean number of observed insertions per position (105 and 102 for +6 bp and +9 bp, respectively). These results are not corrected for codon ambiguity, which increases the unevenness of the distribution.

**(B)** Distribution of the number of positions in the gene encoding *wt*PTE *versus* the number of distinct insertions observed by deep sequencing.

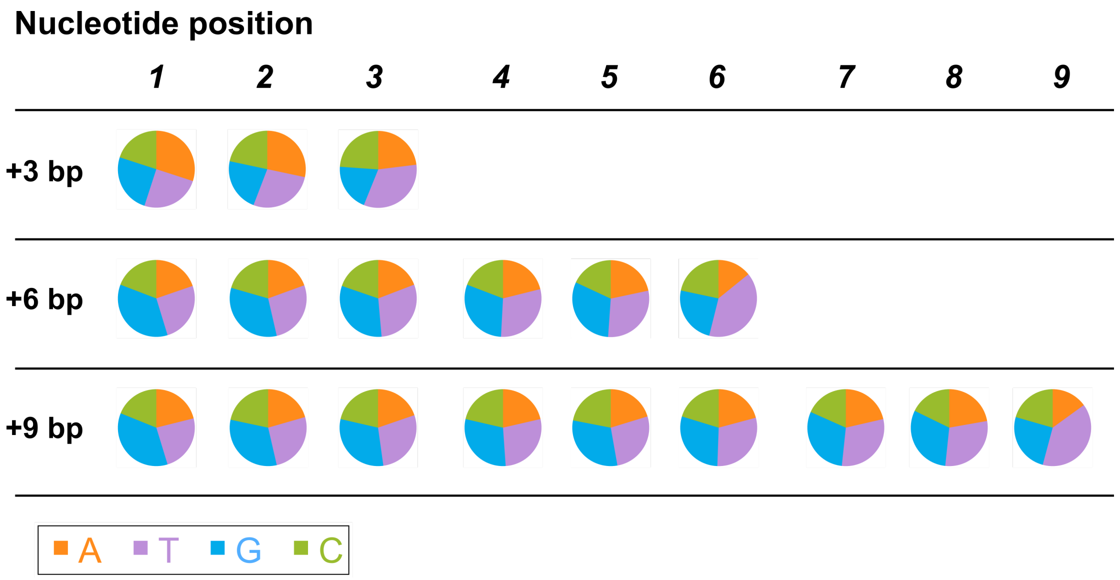

**Supplementary Figure S9: Distribution of nucleotide bases in the randomized inserts of the +3, +6 and +9 bp libraries**

The nucleotide percentage distributions of the in-frame insertions observed among all *wt*PTE variants observed during deep sequencing. Every detected insertion contributes equally to this distribution, regardless of frequency in the library.

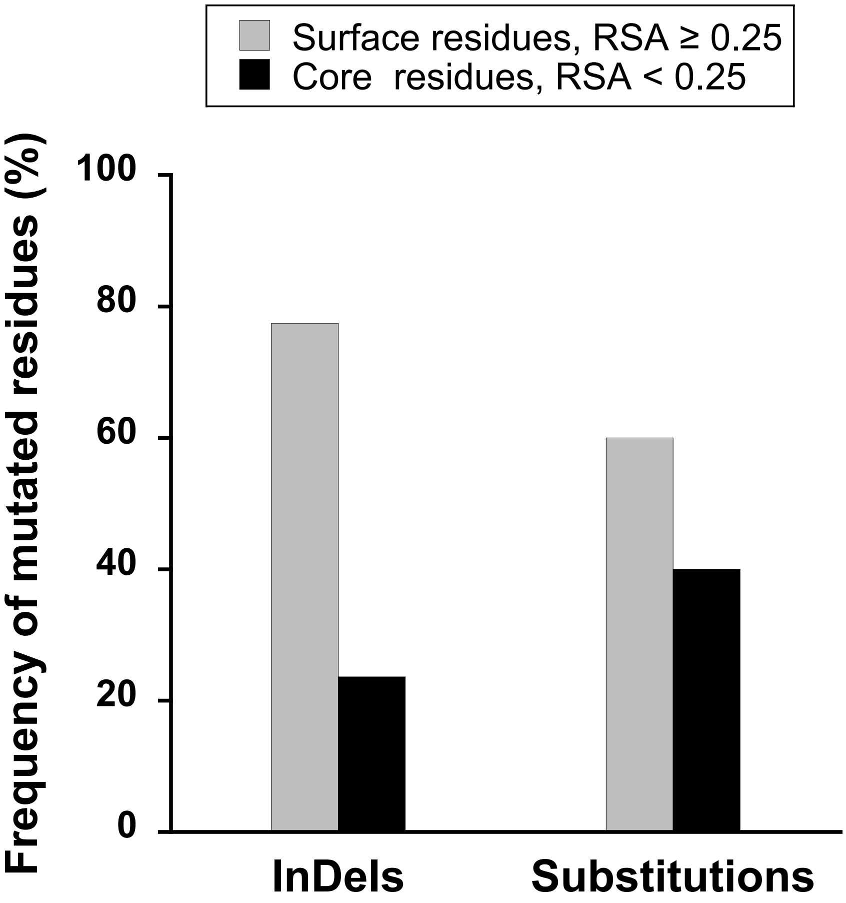

**Supplementary Figure S10: Relationship between mutational tolerance and solvent-accessible surface area (SASA) in *wt*PTE**

The solvent accessible surface area (SASA) of residues mutated (either InDel or substitution) in *wt*PTE variants retaining ≥50% of the parental paraoxonase activity was calculated from the structure of *wt*PTE (PDB code: 4PCP) using the PISA web server at the European Bioinformatics Institute (http://www.ebi.ac.uk/pdbe/prot_int/pistart.html) [14]. Relative accessible surface area (RSA) was defined as the ratio of the SASA for a given residue within the structured protein *vs*. in the free residue [15]. Residues were classified as core for RSA < 0.25, and surface for RSA ≥ 0.25 [16]. Mutated residues are listed in Supplementary Table S10.

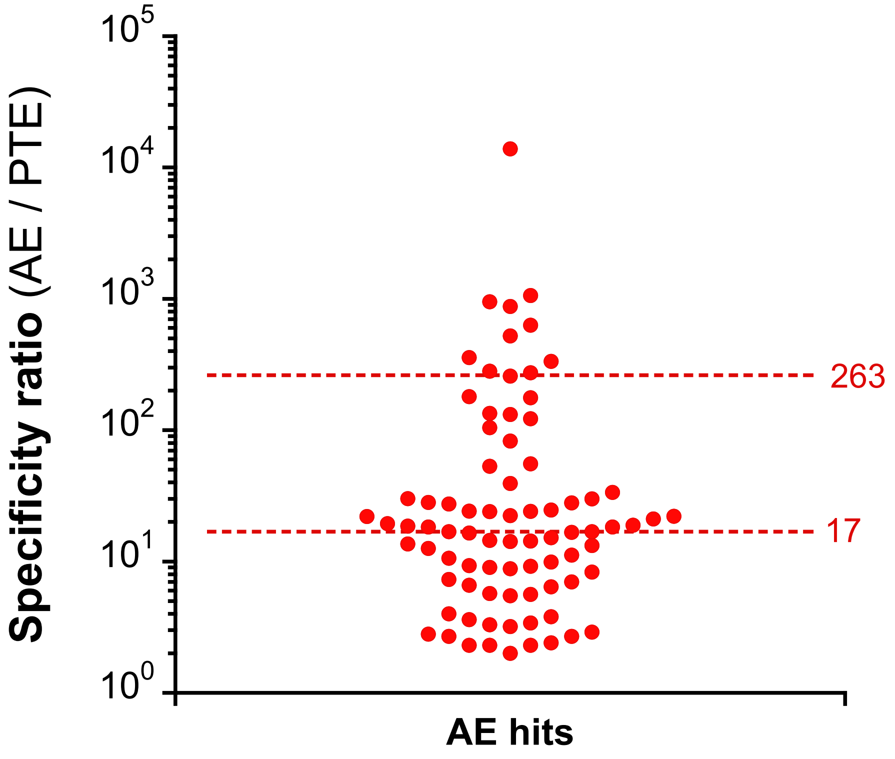

**Supplementary Figure S11: Activity trade-offs in *wt*PTE InDel variants improved in arylesterase activity.**

Activity trade-offs among InDel *wt*PTE variants improved in arylesterase activity (AE) hits (Supplementary Table S11) was evaluated by calculating the specificity ratio, *i.e.* the ratio between the level of AE activity in cell lysate and that of phosphotriesterase activity (PTE). This plot shows the specificity ratio for each InDel variants listed in Supplementary Table S11. The average (~260) and median (~17) specificity ratios are also indicated.

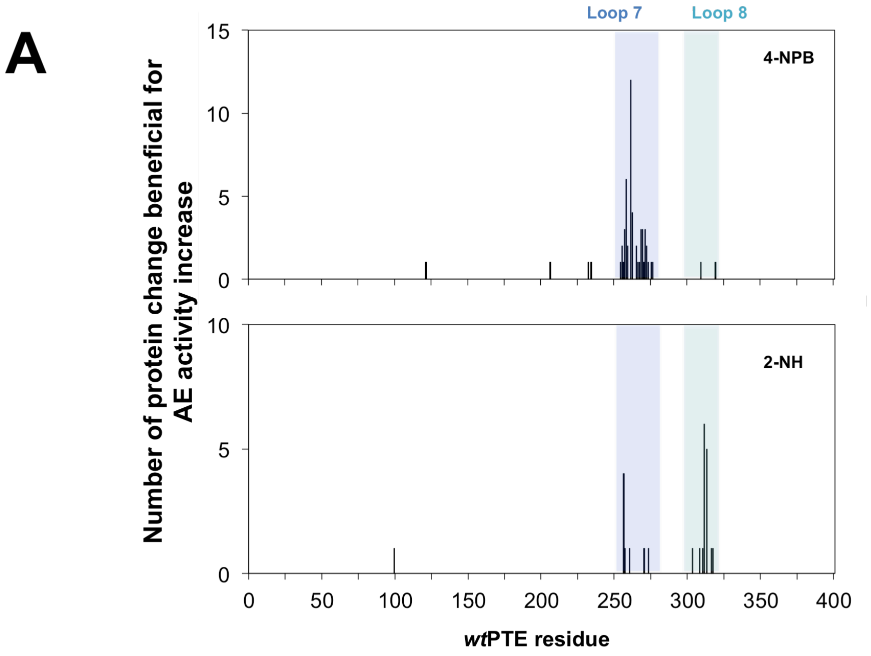

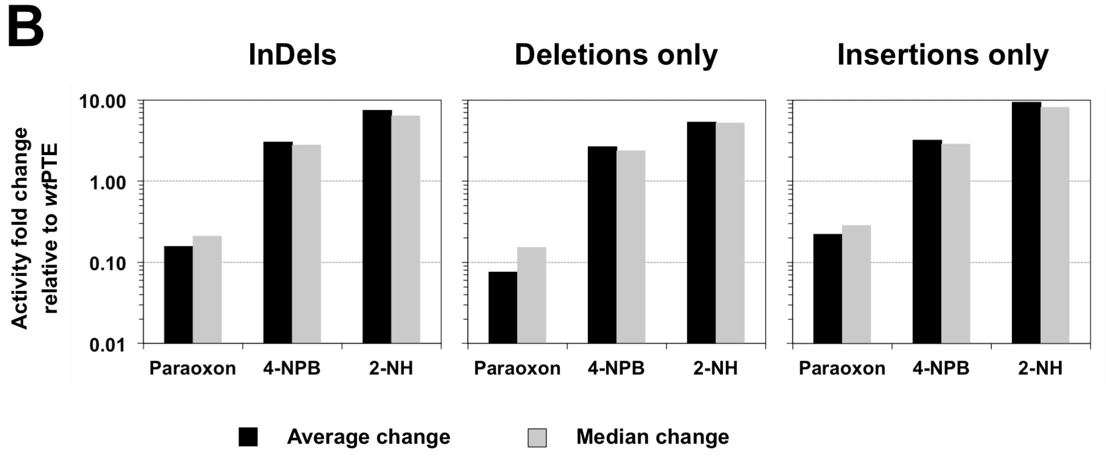

**Supplementary Figure S12: Insertions and deletions improving the arylesterase activity of *wt*PTE by > 2-fold.**

**(A) Location and occurrence of adaptive insertions and deletions in *wt*PTE.** InDels improving arylesterase activity (AE) towards 4-nitrophenyl butyrate (4-NPB; top plot) and 2-naphthyl hexanoate (2-NH; bottom plot) are shown according to their location in the *wt*PTE sequence and the number of their occurrences.

**(B) Average and median activity change in AE-improved *wt*PTE variants.** Values refer to the activity change of all AE-improved variants relative to wtPTE obtained by comparing the initial rates $v_0$ for the hydrolysis of paraoxon (PTE), 4-NPB or 2-NH to that of wtPTE at 200 μM substrate concentration, resulting in a dimensionless ratio. The average change value was determined as the geometric mean of the relative activities of the variants listed in

Supplementary Table S11 and the median change corresponds to the relative activity lying at the midpoint of the recorded relative activities.

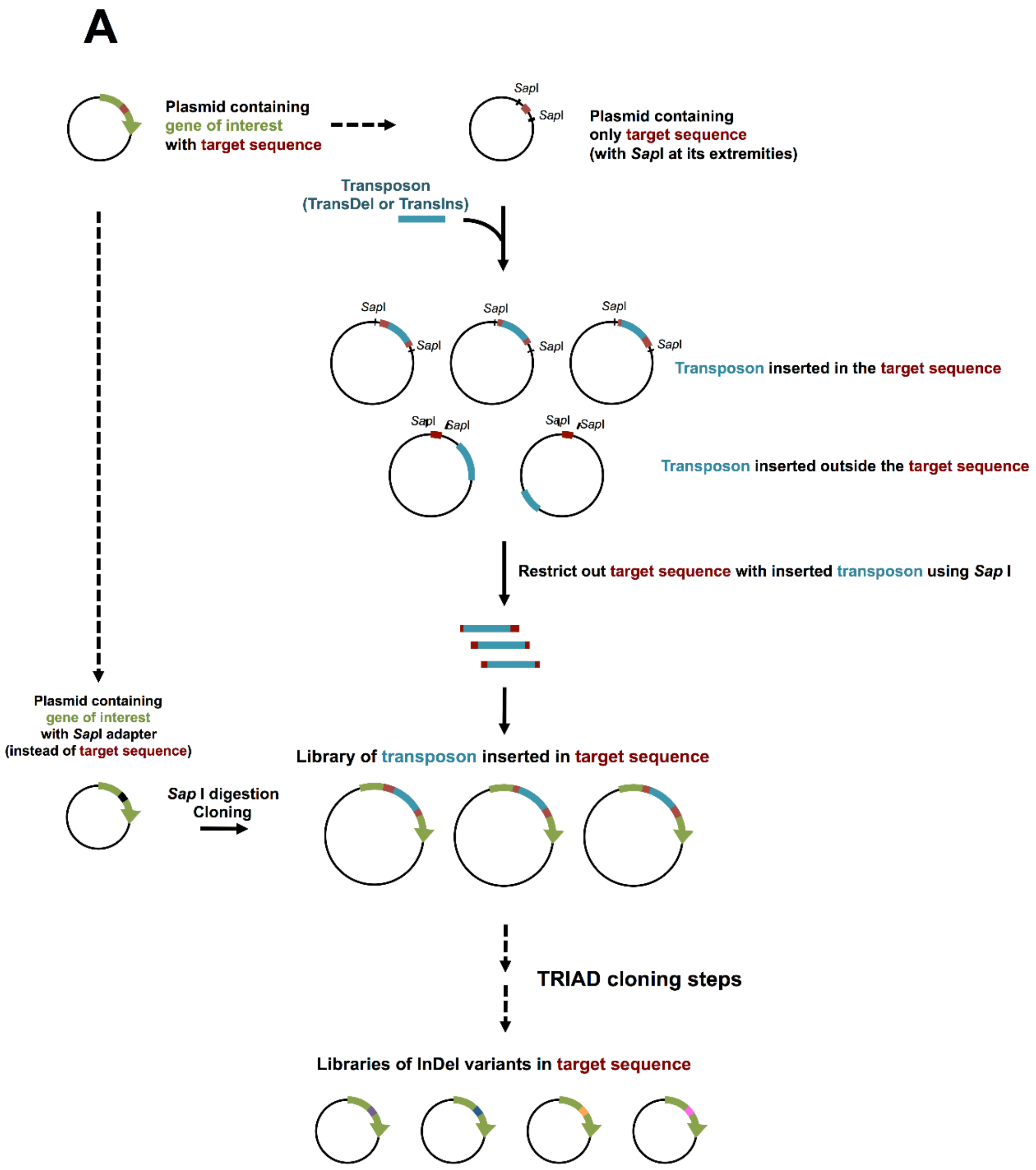

**Supplementary Figure S13** (Continued on next page, legend follows).

(Figure S13 continued)

B

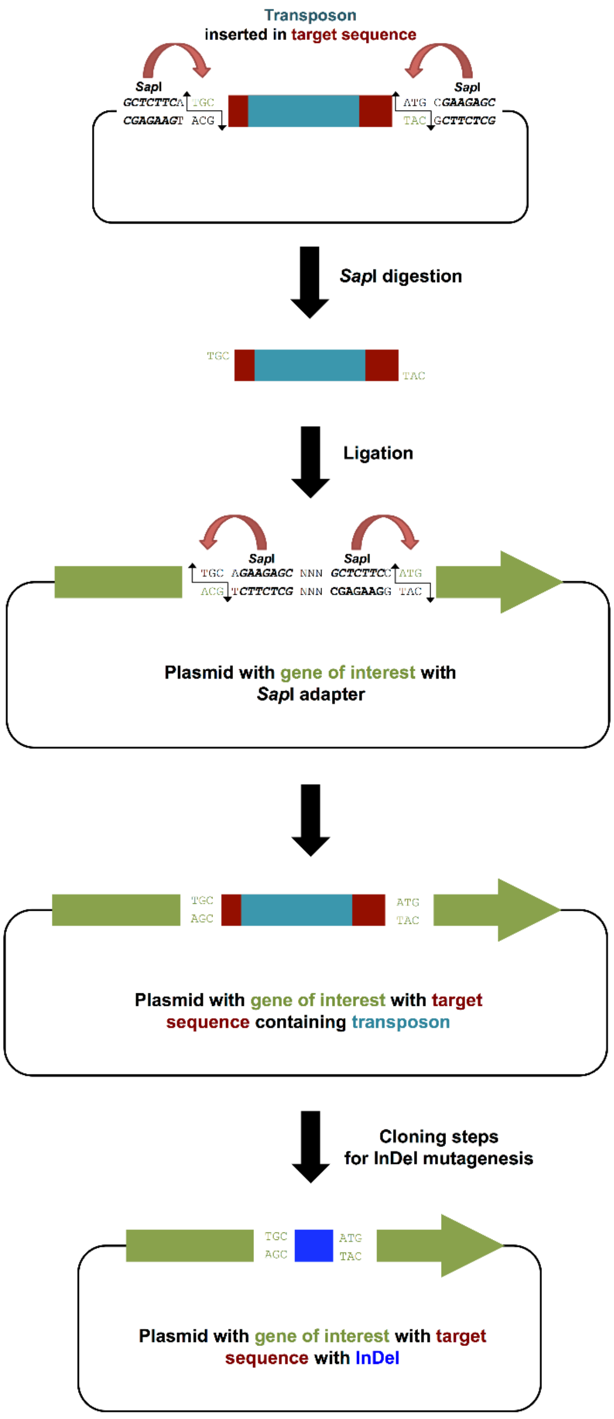

**Supplementary Figure S13: Application of TRIAD for the generation of focused InDel libraries.**

**(A) Schematic outline of the procedure for the generation of focused InDel libraries.** Transposon (*i.e.*, TransDel or TransIns) insertion is carried out on a plasmid containing only the target sequence, 'shielding' the rest of the gene from transposon integration. Target sequences are excised by SapI digestion, and those containing the inserted transposon

(Figure S13 continued)

purified by virtue of their larger size. These fragments can subsequently be cloned into a 'SapI-adapter' plasmid, re-forming the whole gene; now containing transposon only in the target region. Finally, InDel mutagenesis is achieved following the cloning steps illustrated in Figure 1 and Figure S1.

(B) Mechanism for the cloning of focused transposon insertion libraries by seamless cloning using the type IIS restriction enzyme SapI. Seamless cloning with SapI allows cloning the target sequence without altering the original DNA sequence of the gene of interest. In the illustrated example, sequences flanking the target region (TGC and ATG) belong to the gene of interest. Upon digestion with SapI cohesive ends are generated to enable the seamless fusion of the target sequence (containing the inserted transposon) within its original gene sequence. Note that SapI is used here as an example and that any other analogous type IIS restriction enzyme (*e.g.*, BsaI or FokI) could be used to achieve this strategy.

#### 3. SUPPLEMENTARY TABLES

**Supplementary Table S1. Mutagenesis efficiency of TRIAD – individual variants**

Libraries were generated from *wt*PTE. Upon the final transformation step, randomly chosen variants were sequenced by the Sanger method (see also Supplementary Tables S2-3). **In-frame InDels** (*i.e.*, InDels of multiple of three nucleotides) can result in adjacent substitutions (enumerated in the "*InDels with adjacent substitution*" sub-category) depending on the insertion point of the transposon. The number of *unique sequences* was recorded among the observed correct sequences. The *Unique sequences* sub-category refers to in-frame InDels that are observed at least once (see also Supplementary Table S3).

| | **Deletions** | | | | **Insertions** | | | | |
|---|---|---|---|---|---|---|---|---|---|
| **Library** | **-3 bp** | **-6 bp** | **-9 bp** | **All deletions** | **+3 bp** | **+6 bp** | **+9 bp** | **All insertions** | **All Indels** |
| Total number of sequenced variants | 21 | 21 | 22 | 64 (100%) | 23 | 16 | 18 | 57 (100%) | 121 (100%) |
| In-frame InDels | 21 | 17 | 17 | 55 (86%) | 11 | 12 | 12 | 35 (61%) | 90 (74%) |
| InDels with no adjacent substitution | 17 | 12 | 9 | 38 (59%) | 7 | 10 | 8 | 25 (44%) | 63 (52%) |
| InDels with adjacent substitution | 4 | 5 | 8 | 17 (27%) | 4 | 2 | 4 | 10 (18%) | 27 (22%) |
| Unique sequences | 20 | 17 | 17 | 54 (84%) | 11 | 12 | 12 | 35 (61%) | 89 (74%) |
| Frameshifting InDels | 0 | 4 | 5 | 9 (14%) | 12 | 4 | 6 | 22 (39%) | 31 (26%) |

**Supplementary Table S2. Sequence analysis of naïve InDel libraries of *wt*PTE obtained with TRIAD.**

Sequences were determined from randomly chosen variants upon generation of the libraries. Residues are numbered according to the crystal structure of *wt*PTE (PDB: 4PCP). The symbol Δ before a residue (or a group of residues) signifies that this (or these) residue(s) have been deleted. Inserted residues are labelled using the number of the position after which they are inserted and alphabetical order (e.g., glutamine and tyrosine residues inserted in this order after the residues at position 230 would be labelled Q230aY230b).

| Library | Variant number | DNA change | Length change (bp) | Protein mutation |
|---|---|---|---|---|
| -3 bp | 1 | A(TGG)C | -3 | ΔG157 |
| | 2 | A(GGA)A | -3 | ΔE144 |
| | 3 | T(TGC)G | -3 | L66C/ΔR67 |
| | 4 | C(CAC)A | -3 | ΔH230 |
| | 5 | G(GCG)A | -3 | ΔA126 |
| | 6 | C(CGG)G | -3 | ΔR189 |
| | 7 | T(TCG)A | -3 | F104Y/ΔD105 |
| | 8 | G(CTA)T | -3 | ΔY309 |
| | 9 | A(TGA)A | -3 | ΔM293 |
| | 10 | G(TCT)A | -3 | ΔL262 |
| | 11 | A(AGA)T | -3 | ΔE263 |
| | 12 | C(TGG)G | -3 | L272R/ΔG273 |
| | 13 | G(CCG)A | -3 | ΔA114 |
| | 14 | C(TGA)G | -3 | ΔL330 |
| | 15 | C(GGG)C | -3 | R280P/ΔA281 |
| | 16 | C(TAC)G | -3 | ΔL336 |
| | 17 | T(CTA)G | -3 | ΔL252 |
| | 18 | G(GAG)T | -3 | ΔE181 |
| | 19 | C(TAC)A | -3 | ΔY292 |
| | 20 | C(TAC)A | -3 | ΔY292 |
| | 21 | A(TGG)G | -3 | ΔM33 |
| -6 bp | 1 | AAT(GACTGGC)TGT | -7 | frameshift |
| | 2 | TCC(GAGGGCT)TGAG | -7 | frameshift |
| | 3 | ATC(ATGGAC)GTG | -6 | ΔM314D315 |
| | 4 | CAA(TACCGT)GCG | -6 | N38K/ΔT39K40 |
| | 5 | TAA(CCACTC)ACA | -6 | T199N/ΔT200H201 |
| | 6 | AGG(CTACAT)GAA | -6 | ΔY292M293 |
| | 7 | CCA(CTGAGA)GTG | -6 | ΔL330R331 |
| | 8 | TCG(TGGCAA)ACA | -6 | ΔW277Q278 |
| | 9 | TCG(CGGGCT)GCC | -6 | ΔR118A119 |
| | 10 | CGG(TAACCACTCAC)ACG | -11 | frameshift |
| | 11 | GAT(AATGCG)AGT | -6 | ΔN265A266 |
| | 12 | ATT(CCTACG)AGA | -6 | F335L/ΔL336R337 |
| | 13 | GAC(TAACCC)GGC | -6 | ΔN353P354 |

(Table S2 continued)

| Library | Variant number | DNA change | Length change (bp) | Protein mutation |
|---|---|---|---|---|
| | 14 | AAT(CCGAGG)GCT | -6 | S218C/ΔE219G220 |
| | 15 | GAG(GAGTGT)AGA | -6 | ΔS142V143 |
| | 16 | GTG(CGCGGT)CCT | -6 | ΔR41G42 |
| | 17 | TTG(GCCGGA)GTT | -6 | ΔP70E71 |
| | 18 | AAC(CCCGAC)GGA | -6 | ΔP322D323 |
| | 19 | TCC(GAGGGC)TTG | -6 | ΔE219G220 |
| | 20 | GGT(GTTCCGG)TAA | -7 | frameshift |
| | 21 | ACC(CGGCGC)GGT | -6 | P354R/ΔA355R356 |
| -9 bp | 1 | CCC(TCACGGGTTT)GTA | -10 | frameshift |
| | 2 | CGT(TCGTGGCAA)ACA | -9 | ΔS276W277Q278 |
| | 3 | CTT(CCTGCGTGA)GAT | -9 | F150L-ΔL151R152E153 |
| | 4 | AAA(AGGCTGTGA)GAG | -9 | ΔK82A83V84 |
| | 5 | AAC(ATCATGGAC)GTG | -9 | ΔI313M314D315 |
| | 6 | ATT(CCACTGAGA)GTG | -9 | ΔP329L330R331 |
| | 7 | CAG(CTCGGCAGG)ATT | -9 | S61R/ΔS62A63G64 |
| | 8 | GCA(AGTCAGCGC)GGT | -9 | ΔS205Q206R207 |
| | 9 | CGA(CCACAGGCA)AGG | -9 | ΔT172T173G174 |
| | 10 | ACC(GGCTTGTGG)CTC | -9 | ΔG129L130W131 |
| | 11 | AAG(GGCGGCCGC)CCG | -9 | R185S/ΔA186A187A188 |
| | 12 | G**CCC**C -> G**AATT**C | +1 | frameshift |
| | 13 | TCA(TCGACCAAG)GCT | -9 | I288S/ΔD289Q290G291 |
| | 14 | CCT(CCATGGGCG)ATC | -9 | S32Y/ΔM33G34D35 |
| | 15 | CTC(TAGCGGAAA)AGG | -9 | L79Q/ΔA80E81K82 |
| | 16 | GTT(TCGCGGGCTG)CCG | -10 | frameshift |
| | 17 | CGA(TTGGTCTAG)AAG | -9 | I260K/ΔG261L262E263 |
| | 18 | TCA(AGGCGCTCA)TCG | -9 | K285M/ΔA286L287I288 |
| | 19 | GAT(CTCGGTCGC)GAC | -9 | ΔL106G107R108 |
| | 20 | C(TCG)C | -3 | L243P/ΔA244 |
| | 21 | GCT(GCGCGCGGAT)ACC | -10 | frameshift |
| | 22 | GAT(ACTGACGAT)TTG | -9 | ΔT234D235D236 |
| +3 bp | 1 | TT**+C GG+**G CGC | +3 | L87F/G87a |
| | 2 | TT**+G TT+**A GTG | +3 | L182a |
| | 3 | TCCTAT**+TT+**CACAAT | +2 | frameshift |
| | 4 | CTA +CGA+ GAA | +3 | R262a |
| | 5 | T**GGC**A -> T**TT**A | -1 | frameshift |
| | 6 | GA**+C AT+**G GAA | +3 | E144D/M144a |
| | 7 | A**C**G -> A**GGT**G | +2 | frameshift |
| | 8 | TCG **+ACT+** TGG | +3 | T276a |
| | 9 | ACC**+AT+**CGG | +2 | frameshift |
| | 10 | TT**+A CC+**C CTG | +3 | F150L/P150a |
| | 11 | C**CG**A -> C**ACGG**A | +2 | frameshift |
| | 12 | AGA**+CT+**GGA | +2 | frameshift |
| | 13 | TTC**+T+**CGG | +1 | frameshift |
| | 14 | AA**+A TA+**T ACC | +3 | N38K/Y38a |

(Table S2 continued)

| Library | Variant number | DNA change | Length change (bp) | Protein mutation |
|---|---|---|---|---|
| | 15 | GGT**+AA+**CTA | +2 | frameshift |
| | 16 | A**TT**G -> A**ATAT**G | +2 | frameshift |
| | 17 | GGT +GTT+ CTA | +3 | V261a |
| | 18 | ACC **+TTC+** CAC | +3 | F54a |
| | 19 | GG**+A TT+**C ATC | +3 | F157a |
| | 20 | TAG**+GA+**AAG | +2 | frameshift |
| | 21 | GAT **+AAT+** GAT | +3 | N232a |
| | 22 | TCT**+AC+**CCG | +2 | frameshift |
| | 23 | T**G**T -> T**TT**T | +1 | frameshift |
| +6 bp | 1 | GG+G GGG AT+C | +6 | G229aI229b |
| | 2 | TGG +ATG TTT+ CCG | +6 | M69aF69b |
| | 3 | CAG +AAT CTG+ GAA | +6 | N343aL343b |
| | 4 | TCC +CAG AGT+ GAG | +6 | Q47aS47b |
| | 5 | A+AT GAA C+TC | +6 | I44N/E44aL44b |
| | 6 | C+GG ATC C+CC | +6 | R177aI177b |
| | 7 | ACC+TCGTC+GGT | +5 | frameshift |
| | 8 | CG+G AAA TA+C | +6 | K76aY76b |
| | 9 | CC+G ATG CC+C | +6 | M178aP178b |
| | 10 | GGC +GTT AGC+ TAC | +6 | V291aS291b |
| | 11 | GG+G TTA AC+C | +6 | L157aT157b |
| | 12 | TGG +GGT GTA+ CCG | +6 | G69aV69b |
| | 13 | see note [1] | -61 | frameshift |
| | 14 | T+GG TTC G+TG | +6 | L66W/F66aV66b |
| | 15 | CCA+GGTAT+CAG | +5 | frameshift |
| | 16 | TTA+GTGGC+TCA | +5 | frameshift |
| +9 bp | 1 | GCC+GTAAGGTT+CAG | +8 | frameshift |
| | 2 | ACC +GAT TAA TGC+ CAC | +9 | D54a - Stop |
| | 3 | GCT +TGT TGT CCT+ CTC | +9 | C281aC281bP281c |
| | 4 | GA+A AGC TAT GA+G GAA | +9 | S144aY144bE144c |
| | 5 | GTC+CCCACGTCT+CGA | +8 | frameshift |
| | 6 | CTG+GGGTATGG+CGT | +8 | frameshift |
| | 7 | TC+A TAT GGA AT+C | +9 | Y218aG218bI218c |
| | 8 | GGT+CCTCTGGG+TTG | +8 | frameshift |
| | 9 | CG+G GTG TGT CG+CA | +9 | V76aC76bR76c |
| | 10 | GA+A ACT GCA AA+C | +9 | D235E/T235aA235bN235c |
| | 11 | G+GA TCG TGG T+CC | +9 | A90G/S90aW90bS90c |
| | 12 | CC+A GAG ACC GT+G | +9 | E256aT256bV256c |
| | 13 | C+CG AGT TGA T+TG | +9 | L330P/S330a - Stop |
| | 14 | CA+G GGT GGT AT+C | +9 | H57Q/G57aG57bI57c |
| | 15 | TAC +ATC GTT TCG+ ATG | +9 | I292aV292bS292c |
| | 16 | TCA+CCCGTCT+CGG | +7 | frameshift |
| | 17 | GG+A ACA GTG CG+T | +9 | T273aV273bR273c |
| | 18 | ATG+CTTTGGGG+GCA | +8 | frameshift |

(Table S2 continued)

[1] This variant showed a large sequence substitution where C**CATGGGCGAT-CGGATCAATACCGTGCGCGGTCCTATCACAATCTCCGAGGCGGGTTTCACACTAACC**C (modified sequence in bold) was exchanged for C**TCCAGG**C (in bold), resulting in a 61 bp deletion.

**Supplementary Table S3. Frequency of in-frame InDels observed among randomly sequenced *wt*PTE variants (as recorded in Table S2).**

Residues are numbered according to the crystal structure of *wt*PTE (PDB: 4PCP).

| Protein position | Observed InDel | Library | Frequency | Variant name |
|---|---|---|---|---|
| 32 | S32Y/ΔM33G34D35 | -9 bp | 1 | 15 |
| 33 | ΔM33 | -3 bp | 1 | 21 |
| 38 | N38K/ΔT39K40 | -6 bp | 1 | 4 |
| 38 | N38K/Y38a | +3 bp | 1 | 14 |
| 41 | ΔR41G42 | -6 bp | 1 | 18 |
| 44 | I44N/E44aL44b | +6 bp | 1 | 5 |
| 47 | Q47aS47b | +6 bp | 1 | 4 |
| 54 | F54a | +3 bp | 1 | 19 |
| 54 | D54a - Stop | +9 bp | 1 | 2 |
| 57 | H57Q/G57aG57bI57c | +9 bp | 1 | 16 |
| 61 | S61R/ΔS62A63G64 | -9 bp | 1 | 8 |
| 66 | L66C/ΔR67 | -3 bp | 1 | 3 |
| 66 | L66W/F66aV66b | +6 bp | 1 | 17 |
| 69 | M69aF69b | +6 bp | 1 | 2 |
| 69 | G69aV69b | +6 bp | 1 | 14 |
| 70 | ΔP70E71 | -6 bp | 1 | 19 |
| 76 | K76aY76b | +6 bp | 1 | 9 |
| 76 | V76aC76bR76c | +9 bp | 1 | 9 |
| 79 | L79Q/ΔA80E81K82 | -9 bp | 1 | 16 |
| 82 | ΔK82A83V84 | -9 bp | 1 | 4 |
| 87 | L87F/G87a | +3 bp | 1 | 1 |
| 90 | A90G/S90aW90bS90c | +9 bp | 1 | 11 |
| 104 | F104Y/ΔD105 | -3 bp | 1 | 7 |
| 106 | ΔL106G107R108 | -9 bp | 1 | 20 |
| 114 | ΔA114 | -3 bp | 1 | 13 |
| 118 | ΔR118A119 | -6 bp | 1 | 9 |
| 126 | ΔA126 | -3 bp | 1 | 5 |
| 129 | ΔG129L130W131 | -9 bp | 1 | 11 |
| 142 | ΔS142V143 | -6 bp | 1 | 17 |
| 144 | ΔE144 | -3 bp | 1 | 2 |
| 144 | E144D/M144a | +3 bp | 1 | 6 |
| 144 | S144aY144bE144c | +9 bp | 1 | 4 |
| 150 | F150L/ΔL151R152E153 | -9 bp | 1 | 3 |
| 150 | F150L/P150a | +3 bp | 1 | 10 |
| 157 | ΔG157 | -3 bp | 1 | 1 |
| 157 | F157a | +3 bp | 1 | 20 |
| 157 | L157aT157b | +6 bp | 1 | 12 |
| 172 | ΔT172T173G174 | -9 bp | 1 | 10 |
| 177 | R177aI177b | +6 bp | 1 | 7 |
| 178 | M178aP178b | +6 bp | 1 | 10 |

(Table S3 continued)

| Protein position | Observed InDel | Library | Frequency | Variant name |
|---|---|---|---|---|
| 181 | ΔE181 | -3 bp | 1 | 18 |
| 182 | L182a | +3 bp | 1 | 2 |
| 185 | R185S/ΔA186A187A188 | -9 bp | 1 | 12 |
| 189 | ΔR189 | -3 bp | 1 | 6 |
| 199 | T199N/ΔT200H201 | -6 bp | 1 | 5 |
| 205 | ΔS205Q206R207 | -9 bp | 1 | 9 |
| 218 | S218C/ΔE219G220 | -6 bp | 1 | 16 |
| 218 | Y218aG218bI218c | +9 bp | 1 | 7 |
| 219 | ΔE219G220 | -6 bp | 1 | 22 |
| 229 | G229aI229b | +6 bp | 1 | 1 |
| 230 | ΔH230 | -3 bp | 1 | 4 |
| 232 | N232a | +3 bp | 1 | 22 |
| 234 | ΔT234D235D236 | -9 bp | 1 | 23 |
| 235 | D235E/T235aA235bN235c | +9 bp | 1 | 10 |
| 252 | ΔL252 | -3 bp | 1 | 17 |
| 256 | E256aT256bV256c | +9 bp | 1 | 12 |
| 260 | I260K/ΔG261L262E263 | -9 bp | 1 | 18 |
| 261 | V261a | +3 bp | 1 | 17 |
| 262 | ΔL262 | -3 bp | 1 | 10 |
| 262 | R262a | +3 bp | 1 | 4 |
| 263 | ΔE263 | -3 bp | 1 | 11 |
| 265 | ΔN265A266 | -6 bp | 1 | 13 |
| 272 | L272R/ΔG273 | -3 bp | 1 | 12 |
| 273 | T273aV273bR273c | +9 bp | 1 | 19 |
| 276 | ΔS276W277Q278 | -9 bp | 1 | 2 |
| 276 | T276a | +3 bp | 1 | 8 |
| 277 | ΔW277Q278 | -6 bp | 1 | 8 |
| 280 | R280P/ΔA281 | -3 bp | 1 | 15 |
| 281 | C281aC281bP281c | +9 bp | 1 | 3 |
| 285 | K285M/ΔA286L287I288 | -9 bp | 1 | 19 |
| 288 | I288S/ΔD289Q290G291 | -9 bp | 1 | 14 |
| 291 | V291aS291b | +6 bp | 1 | 11 |
| 292 | ΔY292 | -3 bp | 2 | 19, 20 |
| 292 | ΔY292M293 | -6 bp | 1 | 6 |
| 292 | I292aV292bS292c | +9 bp | 1 | 17 |
| 293 | ΔM293 | -3 bp | 1 | 9 |
| 309 | ΔY309 | -3 bp | 1 | 8 |
| 313 | ΔI313M314D315 | -9 bp | 1 | 5 |
| 314 | ΔM314D315 | -6 bp | 1 | 3 |
| 322 | ΔP322D323 | -6 bp | 1 | 21 |
| 329 | ΔP329L330R331 | -9 bp | 1 | 6 |
| 330 | ΔL330 | -3 bp | 1 | 14 |
| 330 | ΔL330R331 | -6 bp | 1 | 7 |
| 330 | L330P/S330a - Stop | +9 bp | 1 | 14 |

(Table S3 continued)

| Protein position | Observed InDel | Library | Frequency | Variant name |
|---|---|---|---|---|
| 335 | F335L/ΔL336R337 | -6 bp | 1 | 14 |
| 336 | ΔL336 | -3 bp | 1 | 16 |
| 343 | N343aL343b | +6 bp | 1 | 3 |
| 353 | ΔN353P354 | -6 bp | 1 | 15 |
| 354 | P354R/ΔA355R356 | -6 bp | 1 | 24 |

**Supplementary Table S4A: deep sequencing coverage statistics.**

All six libraries were sequenced as part of one MiSeq 2×75 bp run. Since insertion libraries have a greater theoretical diversity, they were loaded onto the flow cell at 3× the amount of deletion libraries.

| **Library** | **-3 bp** | **-6 bp** | **-9 bp** |
|---|---|---|---|
| **Total reads** | $1.04\times10^{6}$ | $1.09\times10^{6}$ | $8.99\times10^{5}$ |
| **Assembled reads** | $7.50\times10^{5}$ | $8.22\times10^{5}$ | $6.48\times10^{5}$ |
| **Alignment rate** | 96.4% | 95.1% | 90.3% |
| **Unassembled reads** | $2.86\times10^{5}$ | $2.67\times10^{5}$ | $2.51\times10^{5}$ |
| **Alignment rate** | 93.5% | 95.9% | 88.6% |
| **Total aligned reads** | $9.90\times10^{5}$ | $1.04\times10^{6}$ | $8.07\times10^{5}$ |
| **Mean ± SD coverage per base** | $(8.59\pm1.7)\times10^{4}$ | $(8.84\pm1.7)\times10^{4}$ | $(6.99\pm1.6)\times10^{4}$ |

| **Library** | **+3 bp** | **+6 bp** | **+9 bp** |
|---|---|---|---|
| **Total reads** | $3.36\times10^{6}$ | $3.38\times10^{6}$ | $3.09\times10^{6}$ |
| **Assembled reads** | $2.56\times10^{6}$ | $2.51\times10^{6}$ | $2.47\times10^{6}$ |
| **Alignment rate** | 97.1% | 95.0% | 95.8% |
| **Unassembled reads** | $8.08\times10^{5}$ | $8.67\times10^{5}$ | $6.28\times10^{5}$ |
| **Alignment rate** | 96.6% | 94.0% | 95.6% |
| **Total aligned reads** | $3.27\times10^{6}$ | $3.20\times10^{6}$ | $2.968\times10^{6}$ |
| **Mean ± SD coverage per base** | $(2.76\pm0.54)\times10^{5}$ | $(2.73\pm0.58)\times10^{5}$ | $(2.38\pm0.45)\times10^{5}$ |

**Supplementary Table S4B: Proportion of frameshifts**

The proportion of variants containing frameshifts was estimated as follows:

$$Est.\% \, frameshifted \; variants = \frac{frameshift \; reads/all \; reads}{read \; length/length \; of \; gene}$$

Where read length is equal to 2×75=150 bp and the gene length is 999 bp.

| | Deletions | | |
|---|---|---|---|
| Sequencing reads | -3 bp | -6 bp | -9 bp |
| All reads | $9.90\times10^{5}$ | $1.04\times10^{6}$ | $8.07\times10^{5}$ |
| Reads with target mutations | 74923 | 48911 | 42358 |
| % with target mutations | 7.6% | 4.7% | 5.3% |
| Reads with frameshifts | 6082 | 30813 | 17163 |
| % of reads with frameshifts | 0.6% | 3.0% | 2.1% |
| Est. % frameshifted variants | 4.1% | 19.8% | 14.2% |

| | Insertions | | |
|---|---|---|---|
| Sequencing reads | +3 bp | +6 bp | +9 bp |
| All reads | $3.27\times10^{6}$ | $3.20\times10^{6}$ | $2.968\times10^{6}$ |
| Reads with target mutations | 121089 | 145374 | 115899 |
| % with target mutations | 3.7% | 4.5% | 3.9% |
| Reads with frameshifts | 179412 | 139738 | 117505 |
| % of reads with frameshifts | 5.5% | 4.4% | 4.0% |
| Est. % frameshifted variants | 36.6% | 29.1% | 26.4% |

**Supplementary Table S5: TransDel consensus site preference**

(see WebLogo in Figure 3A)

Mu transposons insert within a five-nucleotide sequence, which is duplicated during the insertion. The insertion preference of TransDel in *wt*PTE gene can be calculated with precision from the location of -3 bp deletions, because these deletions are symmetrical and centred within the insertion site. The consensus site preference is built from detected -3 bp mutations, weighed according to frequency of occurrence and adjusted for orientation of transposon and GC composition.

| **Position** | **A** | **C** | **G** | **T** |
|---|---|---|---|---|
| **1** | 14.4% | 34.4% | 25.7% | 25.5% |
| **2** | 12.4% | 38.3% | 17.7% | 31.7% |
| **3** | 16.5% | 33.5% | 33.5% | 16.5% |
| **4** | 31.7% | 17.7% | 38.3% | 12.4% |
| **5** | 25.5% | 25.7% | 34.4% | 14.4% |

**Supplementary Table S6: Statistics on number of reads supporting each variant**

Each observed variant is associated with a count: this gives the number of reads (either an assembled paired-end 2×75 bp or a single end read from an unassembled pair) that support the detection of that variant. Some variants are observed more frequently than others. This frequency reflects the inherent bias of Mu transposon insertion, amplification bias during transformations (where one variant may randomly grow to greater abundance than another), and stochastic fluctuations resulting from sequencing. This table summarizes the *median* count in the -3 bp and +3 bp libraries, as well as maximum and interquartile range. The distribution is shown in Supplementary Figure S7 as a histogram.

| **Count** | ***wt*PTE -3 bp** | ***wt*PTE + 3bp** |
|---|---|---|
| **Minimum observed** | 0 | 0 |
| **Q1** | 4.9 | 11.0 |
| **Median** | 22.0 | 47.2 |
| **Q3** | 75.5 | 133.3 |
| **Maximum** | 1420 | 3336 |
| **Q3-Q1** | 70.6 | 122.3 |

**Supplementary Table S7. Number of reads per distinct deletion observed by deep sequencing in *wt*PTE deletion libraries generated *via* TRIAD.** The histograms relative to these distributions are plotted in Supplementary Figure S7.

| | Number of distinct deletions | | | | | | | |
|---|---|---|---|---|---|---|---|---|
| | **- 3 bp** | | **-6 bp** | | **-9 bp** | | **All deletions** | |
| **Total number of deletions** | 633 | | 682 | | 608 | | 1923 | |
| **Number of reads per deletion** | | | | | | | | |
| *1-4 reads* | 83 | | 107 | | 96 | | 286 | |
| *5-9 reads* | 52 | | 67 | | 70 | | 189 | |
| *10-39 reads* | 168 | 310 (49%) | 212 | 369 (54.1%) | 217 | 324 (53.3%) | 597 | 1003 (52,2%) |
| *40-99 reads* | 142 | | 157 | | 107 | | 406 | |
| *100-199 reads* | 86 | | 82 | | 64 | | 232 | |
| *200-999 reads* | 93 | 102 (16.1%) | 57 | 57 (8.4%) | 53 | 64 (8.9%) | 203 | 213 (11.1%) |
| *≥1000 reads* | 9 | | 0 | | 1 | | 10 | |

**Supplementary Table S8. Fitness effects in TRIAD (insertion and deletion) and trinucleotide substitution libraries of *wt*PTE.**

| | | Deletions | | | | Insertions | | | | Substitutions |
|---|---|---|---|---|---|---|---|---|---|---|
| | | **-3 bp** | **-6 bp** | **-9 bp** | **All deletions** | **+3 bp** | **+6 bp** | **+9 bp** | **All insertions** | **TriNEx** |
| | *Number of variants[a]* | 175 | 154 | 156 | 485 | 92 | 134 | 125 | 351 | 342 |
| **PARAOXONASE** | *Fitness effect[b]:* | | | | | | | | | |
| | *Strongly deleterious* | 117 (66.9%) | 143 (92.8%) | 143 (91.7%) | 403 (83.1%) | 58 (63.2%) | 106 (79.1%) | 105 (84.0%) | 269 (76.7%) | 81 (23.8%) |
| | *Mildly deleterious* | 41 (23.4%) | 9 (5.9%) | 11 (7.1%) | 61 (12.6%) | 14 (15.2%) | 20 (14.9%) | 13 (10.4%) | 47 (13.4%) | 100 (29.2%) |
| | *Neutral* | 14 (8.0%) | 2 (1.3%) | 2 (1.3%) | 18 (3.7%) | 18 (19.5%) | 7 (5.2%) | 1 (0.8%) | 26 (7.4%) | 161 (47.0%) |
| | *Beneficial* | 3 (1.7%) | 0 (0.0%) | 0 (0.0%) | 3 (0.6%) | 2 (2.2%) | 1 (0.7%) | 6 (4.8%) | 9 (2.6%) | 0 (0.0%) |
| | *Average fitness change][d]* | 0.048 | 0.014 | 0.015 | 0.022 | 0.054 | 0.024 | 0.019 | 0.027 | 0.28 |
| | | [0.038 ; 0.062] | [0.012 ; 0.016] | [0.013 ; 0.017] | [0.02 ; 0.025] | [0.036 ; 0.082] | [0.018 ; 0.031] | [0.015 ; 0.025] | [0.023 ; 0.033] | [0.24 ; 0.33] |
| | *Median fitness change[c]* | 0.03 | <0.01 | <0.01 | <0.01 | 0.01 | <0.01 | <0.01 | <0.01 | 0.63 |
| | *Minimum fitness change[c]* | <0.01 | <0.01 | <0.01 | <0.01 | <0.01 | <0.01 | <0.01 | <0.01 | <0.01 |
| | *Maximum fitness change[c]* | 1.59 | 0.94 | 1.31 | 1.59 | 1.79 | 1.60 | 2.78 | 2.78 | 1.50 |
| **ARIYLESTERASE (4-nitrophenyl butyrate)** | *Fitness effect[b]:* | | | | | | | | | |
| | *Strongly deleterious* | 100 (57.2%) | 137 (88.9%) | 133 (85.2%) | 370 (76.3%) | 53 (57.7%) | 99 (73.9%) | 68 (54.3%) | 220 (62.7%) | 65 (19.1%) |
| | *Mildly deleterious* | 40 (22.8%) | 10 (6.5%) | 9 (5.8%) | 59 (12.2%) | 16 (17.3%) | 18 (13.4%) | 42 (33.6%) | 76 (21.6%) | 96 (28.0%) |
| | *Neutral* | 14 (8.0%) | 6 (3.9%) | 7 (4.5%) | 27 (5.6%) | 15 (16.3%) | 8 (6.0%) | 5 (4.0%) | 28 (8.0%) | 175 (51.1%) |
| | *Beneficial* | 21 (12.0%) | 1 (0.7%) | 7 (4.5%) | 29 (6.0%) | 8 (8.7%) | 9 (6.7%) | 10 (8.0%) | 27 (7.7%) | 6 (1.8%) |
| | *Average fitness change[c][d]* | 0.07 | 0.017 | 0.02 | 0.03 | 0.085 | 0.031 | 0.098 | 0.061 | 0.34 |
| | | [0.051 ; 0.095] | [0.014 ; 0.02] | [0.015 ; 0.026] | [0.025 ; 0.035] | [0.056 ; 0.13] | [0.023 ; 0.043] | [0.075 ; 0.13] | [0.05; 0.074] | [0.3 ; 0.4] |
| | *Median fitness change[c]* | 0.03 | <0.01 | <0.01 | <0.01 | 0.05 | <0.01 | 0.09 | 0.04 | 0.70 |
| | *Minimum fitness change[c]* | <0.01 | <0.01 | <0.01 | <0.01 | <0.01 | <0.01 | <0.01 | <0.01 | <0.01 |
| | *Maximum fitness change[c]* | 2.46 | 1.52 | 5.02 | 5.02 | 4.12 | 3.13 | 5.36 | 5.36 | 2.98 |

[a] The number of variants sampled in each library was corrected to only take in-frame mutations into account. Overall, 178 variants from each TRIAD library (6 × 178 in total) and 435 trinucleotide substitution variants were randomly picked, expressed in *E. coli* and screened for hydrolysis of paraoxon and 4-nitrophenyl butyrate (see **Table S9**). The *estimated* number of frame-shifted variants (based on the frequencies in **Table S1**) was then subtracted from the numbers of highly deleterious variants (<0.01 in both activities).

(Table S8 continued)

[b] Mutations are classified as strongly deleterious (>10-fold activity decrease relative to wtPTE), mildly deleterious (10-fold—1.5-fold decrease), neutral (<1.5-fold change), and beneficial (>1.5-fold increase).

[c] Changes in phosphotriesterase (native substrate: paraoxon) and esterase (promiscuous substrate: pNPB) activities are determined relative to those of *wt*PTE by comparing the initial rates in cell lysates measured under identical conditions with 200 μM of the respective substrates, resulting in a dimensionless ratio (see Methods).

[d] The average fitness change refers to the change in initial rates as a consequence of mutation and is calculated as the geometric mean of the relative activities of the variants (see Supplementary Table S9). The corresponding confidence intervals (5% risk of error) are indicated between brackets.

**Supplementary Table S9. Functional analysis of TRIAD and trinucleotide substitution libraries of *wt*PTE against paraoxon and 4-NPB**

Changes in paraoxonase (native substrate: paraoxon) and arylesterase (promiscuous substrate: 4-nitrophenylbutyrate, 4-NPB) activities are determined relative to those of *wt*PTE by comparing the initial rates in cell lysates measured under identical conditions with 200 μM of the respective substrates (see Methods). Data are averages of triplicate values from three independent experiments and error values represent +/- 1 SEM.

| | **Relative activity** | | |
|---|---|---|---|
| **Variant** | **Paraoxon** | **4-NPB** | **Protein mutation** |
| **-3 bp** | | | |
| 1 | <0.01 | <0.01 | n.d. |
| 2 | <0.01 | <0.01 | n.d. |
| 3 | 0.03 ± 0.01 | 0.01 ± 0.01 | n.d. |
| 4 | <0.01 | 0.05 ± 0.01 | n.d. |
| 5 | 0.53 ± 0.01 | 0.95 ± 0.02 | ΔR337 |
| 6 | 0.74 ± 0.01 | 1.72 ± 0.05 | ΔS269 |
| 7 | 0.64 ± 0.08 | 0.82 ± 0.12 | ΔA49 |
| 8 | <0.01 | <0.01 | n.d. |
| 9 | 0.11 ± 0.01 | 0.13 ± 0.02 | n.d. |
| 10 | 0.86 ± 0.04 | 2.01 ± 0.01 | ΔS269 |
| 11 | 0.16 ± 0.01 | 2.37 ± 0.01 | L272R/ΔG273 |
| 12 | <0.01 | <0.01 | n.d. |
| 13 | 0.51 ± 0.06 | 0.61 ± 0.06 | ΔG174 |
| 14 | 0.03 ± 0.01 | 0.04 ± 0.01 | n.d. |
| 15 | <0.01 | <0.01 | n.d. |
| 16 | <0.01 | <0.01 | n.d. |
| 17 | <0.01 | 0.33 ± 0.03 | n.d. |
| 18 | <0.01 | <0.01 | n.d. |
| 19 | <0.01 | <0.01 | n.d. |
| 20 | <0.01 | <0.01 | n.d. |
| 21 | 0.62 ± 0.04 | 0.66 ± 0.05 | ΔY292 |
| 22 | 0.23 ± 0.05 | 1.53 ± 0.01 | ΔQ211 |
| 23 | <0.01 | <0.01 | n.d. |
| 24 | 0.17 ± 0.01 | 2.29 ± 0.12 | L272R/ΔG273 |
| 25 | 1.52 ± 0.03 | 1.43 ± 0.03 | ΔL362 |
| 26 | 0.01 ± 0.01 | <0.01 | n.d. |
| 27 | 0.21 ± 0.01 | 0.44 ± 0.01 | n.d. |
| 28 | 0.06 ± 0.01 | 1.93 ± 0.1 | ΔS276 |
| 29 | <0.01 | <0.01 | n.d. |
| 30 | 0.02 ± 0.01 | <0.01 | n.d. |
| 31 | <0.01 | <0.01 | n.d. |
| 32 | <0.01 | <0.01 | n.d. |
| 33 | <0.01 | 0.17 ± 0.01 | n.d. |
| 34 | 0.04 ± 0.01 | 0.65 ± 0.02 | n.d. |

(Table S9 continued)

| Variant | Relative activity | | Protein mutation |
|---|---|---|---|
| | Paraoxon | 4-NPB | |
| 35 | <0.01 | <0.01 | n.d. |
| 36 | 0.59 ± 0.02 | 0.74 ± 0.02 | ΔE338 |
| 37 | 0.09 ± 0.01 | 0.07 ± 0.01 | n.d. |
| 38 | 0.70 ± 0.05 | 1.74 ± 0.15 | ΔL262 |
| 39 | 0.03 ± 0.01 | <0.01 | n.d. |
| 40 | 0.08 ± 0.02 | 0.03 ± 0.01 | n.d. |
| 41 | <0.01 | <0.01 | n.d. |
| 42 | <0.01 | <0.01 | n.d. |
| 43 | 0.46 ± 0.1 | 0.47 ± 0.1 | ΔD289 |
| 44 | 1.02 ± 0.02 | 0.96 ± 0.03 | ΔE263 |
| 45 | 0.22 ± 0.02 | 0.29 ± 0.04 | n.d. |
| 46 | <0.01 | <0.01 | n.d. |
| 47 | 0.78 ± 0.12 | 0.63 ± 0.11 | ΔT173 |
| 48 | 0.50 ± 0.1 | 0.47 ± 0.1 | ΔQ155 |
| 49 | <0.01 | <0.01 | n.d. |
| 50 | <0.01 | <0.01 | n.d. |
| 51 | <0.01 | <0.01 | n.d. |
| 52 | 0.18 ± 0.01 | 1.56 ± 0.05 | n.d. |
| 53 | 0.76 ± 0.06 | 1.90 ± 0.13 | ΔL262 |
| 54 | 1.59 ± 0.06 | 1.42 ± 0.08 | wt |
| 55 | 0.44 ± 0.06 | 1.01 ± 0.1 | n.d. |
| 56 | <0.01 | <0.01 | n.d. |
| 57 | 0.01 ± 0.01 | 0.05 ± 0.01 | n.d. |
| 58 | 0.03 ± 0.01 | 0.04 ± 0.01 | n.d. |
| 59 | 0.24 ± 0.03 | 0.42 ± 0.07 | ΔY239 |
| 60 | 0.69 ± 0.04 | 1.68 ± 0.04 | ΔL262 |
| 61 | <0.01 | <0.01 | n.d. |
| 62 | 0.17 ± 0.02 | 0.32 ± 0.04 | n.d. |
| 63 | 0.11 ± 0.01 | 1.14 ± 0.08 | n.d. |
| 64 | <0.01 | <0.01 | n.d. |
| 65 | 0.03 ± 0.01 | 0.18 ± 0.01 | n.d. |
| 66 | <0.01 | <0.01 | K169I/ΔV170 |
| 67 | <0.01 | <0.01 | ΔH55 |
| 68 | <0.01 | <0.01 | D301E/ΔW302 |
| 69 | <0.01 | <0.01 | ΔE144 |
| 70 | <0.01 | <0.01 | ΔA355 |
| 71 | <0.01 | <0.01 | frameshift |
| 72 | <0.01 | <0.01 | n.d. |
| 73 | 0.04 ± 0.01 | 0.05 ± 0.02 | n.d. |
| 74 | <0.01 | <0.01 | n.d. |
| 75 | <0.01 | <0.01 | n.d. |
| 76 | 0.78 ± 0.02 | 1.19 ± 0.03 | ΔA266 |
| 77 | 0.02 ± 0.01 | <0.01 | n.d. |
| 78 | 0.08 ± 0.01 | 0.08 ± 0.01 | n.d. |

(Table S9 continued)

| Variant | Relative activity | | Protein mutation |
|---|---|---|---|
| | Paraoxon | 4-NPB | |
| 79 | <0.01 | <0.01 | n.d. |
| 80 | <0.01 | <0.01 | n.d. |
| 81 | <0.01 | 0.02 ± 0.01 | n.d. |
| 82 | 0.41 ± 0.03 | 2.27 ± 0.13 | ∆T234 |
| 83 | 0.06 ± 0.01 | <0.01 | n.d. |
| 84 | 0.23 ± 0.05 | 0.63 ± 0.14 | n.d. |
| 85 | <0.01 | <0.01 | ∆E210 |
| 86 | <0.01 | <0.01 | n.d. |
| 87 | <0.01 | <0.01 | n.d. |
| 88 | 0.70 ± 0.01 | 0.42 ± 0.01 | ∆A203 |
| 89 | <0.01 | <0.01 | n.d. |
| 90 | 1.58 ± 0.02 | 1.62 ± 0.04 | ∆R36 |
| 91 | 0.04 ± 0.01 | <0.01 | n.d. |
| 92 | <0.01 | 0.10 ± 0.03 | n.d. |
| 93 | 0.02 ± 0.01 | <0.01 | n.d. |
| 94 | 0.02 ± 0.01 | <0.01 | n.d. |
| 95 | 0.04 ± 0.01 | 1.51 ± 0.12 | n.d. |
| 96 | <0.01 | <0.01 | n.d. |
| 97 | 0.05 ± 0.01 | 0.07 ± 0.02 | V84G/∆R85 |
| 98 | <0.01 | <0.01 | ∆H55 |
| 99 | 0.02 ± 0.01 | <0.01 | n.d. |
| 100 | <0.01 | <0.01 | ∆C59 |
| 101 | <0.01 | 0.57 ± 0.01 | n.d. |
| 102 | 0.06 ± 0.01 | 0.13 ± 0.02 | n.d. |
| 103 | 0.15 ± 0.03 | 0.12 ± 0.03 | n.d. |
| 104 | 0.06 ± 0.01 | 0.06 ± 0.02 | n.d. |
| 105 | <0.01 | <0.01 | n.d. |
| 106 | 0.21 ± 0.04 | 1.49 ± 0.01 | ∆Q211 |
| 107 | 0.78 ± 0.08 | 0.73 ± 0.05 | ∆M293 |
| 108 | 0.04 ± 0.01 | 0.37 ± 0.02 | n.d. |
| 109 | <0.01 | <0.01 | n.d. |
| 110 | <0.01 | <0.01 | n.d. |
| 111 | <0.01 | <0.01 | n.d. |
| 112 | <0.01 | <0.01 | n.d. |
| 113 | 0.05 ± 0.01 | 0.03 ± 0.01 | n.d. |
| 114 | 0.05 ± 0.01 | 0.22 ± 0.02 | n.d. |
| 115 | 0.02 ± 0.01 | 0.28 ± 0.02 | n.d. |
| 116 | <0.01 | <0.01 | n.d. |
| 117 | 0.07 ± 0.01 | <0.01 | n.d. |
| 118 | <0.01 | <0.01 | n.d. |
| 119 | <0.01 | <0.01 | n.d. |
| 120 | 0.06 ± 0.01 | 2.16 ± 0.06 | ∆D232 |
| 121 | 0.34 ± 0.01 | 1.56 ± 0.03 | ∆D235 |
| 122 | 0.51 ± 0.11 | 0.47 ± 0.12 | ∆Q155 |

(Table S9 continued)

| Variant | Relative activity | | Protein mutation |
|---|---|---|---|
| | Paraoxon | 4-NPB | |
| 123 | 0.03 ± 0.01 | 0.03 ± 0.01 | n.d. |
| 124 | <0.01 | <0.01 | n.d. |
| 125 | <0.01 | <0.01 | n.d. |
| 126 | 0.33 ± 0.04 | 1.91 ± 0.02 | n.d. |
| 127 | 0.64 ± 0.02 | 1.95 ± 0.07 | A266G/ΔS267 |
| 128 | <0.01 | <0.01 | ΔW69 |
| 129 | 0.44 ± 0.15 | 0.43 ± 0.14 | ΔR76 |
| 130 | <0.01 | <0.01 | n.d. |
| 131 | 0.83 ± 0.01 | 1.16 ± 0.05 | n.d. |
| 132 | <0.01 | <0.01 | n.d. |
| 133 | 1.33 ± 0.13 | 1.09 ± 0.06 | ΔY292 |
| 134 | <0.01 | <0.01 | n.d. |
| 135 | <0.01 | 0.43 ± 0.01 | n.d. |
| 136 | 0.19 ± 0.01 | 0.13 ± 0.01 | n.d. |
| 137 | 0.54 ± 0.01 | 1.24 ± 0.03 | ΔL262 |
| 138 | 0.16 ± 0.05 | 0.11 ± 0.04 | n.d. |
| 139 | <0.01 | <0.01 | n.d. |
| 140 | <0.01 | <0.01 | n.d. |
| 141 | 0.23 ± 0.02 | 1.91 ± 0.02 | L272R/ΔG273 |
| 142 | <0.01 | <0.01 | n.d. |
| 143 | <0.01 | <0.01 | n.d. |
| 144 | 0.04 ± 0.01 | 0.03 ± 0.01 | n.d. |
| 145 | 0.37 ± 0.08 | 0.27 ± 0.06 | P43L/ΔI44 |
| 146 | 0.05 ± 0.01 | 0.07 ± 0.02 | n.d. |
| 147 | 0.29 ± 0.01 | 0.46 ± 0.01 | ΔA119 |
| 148 | 0.02 ± 0.01 | <0.01 | n.d. |
| 149 | 0.03 ± 0.01 | 0.28 ± 0.02 | n.d. |
| 150 | 0.03 ± 0.01 | <0.01 | n.d. |
| 151 | 0.01 ± 0.01 | <0.01 | n.d. |
| 152 | 0.88 ± 0.01 | 2.46 ± 0.06 | A266G/ΔS267 |
| 153 | 0.05 ± 0.01 | 0.02 ± 0.01 | n.d. |
| 154 | <0.01 | <0.01 | n.d. |
| 155 | 0.29 ± 0.01 | 0.22 ± 0.01 | ΔP197 |
| 156 | <0.01 | <0.01 | n.d. |
| 157 | 0.37 ± 0.06 | 0.42 ± 0.07 | ΔP43 |
| 158 | 0.03 ± 0.01 | <0.01 | n.d. |
| 159 | 0.35 ± 0.02 | 0.29 ± 0.02 | ΔA165 |
| 160 | <0.01 | <0.01 | n.d. |
| 161 | <0.01 | <0.01 | n.d. |
| 162 | 0.03 ± 0.01 | <0.01 | n.d. |
| 163 | <0.01 | <0.01 | n.d. |
| 164 | <0.01 | <0.01 | n.d. |
| 165 | 0.48 ± 0.09 | 0.40 ± 0.09 | V176A/ΔT177 |
| 166 | <0.01 | <0.01 | n.d. |

(Table S9 continued)

| Variant | Relative activity | | Protein mutation |
|---|---|---|---|
| | Paraoxon | 4-NPB | |
| 167 | 0.05 ± 0.01 | 0.23 ± 0.01 | n.d. |
| 168 | 0.40 ± 0.03 | 0.35 ± 0.03 | ΔG74 |
| 169 | <0.01 | <0.01 | n.d. |
| 170 | 0.07 ± 0.02 | 0.03 ± 0.01 | n.d. |
| 171 | 0.33 ± 0.06 | 0.53 ± 0.09 | ΔY239 |
| 172 | <0.01 | <0.01 | n.d. |
| 173 | 0.03 ± 0.01 | 0.64 ± 0.04 | n.d. |
| 174 | <0.01 | <0.01 | n.d. |
| 175 | <0.01 | <0.01 | n.d. |
| 176 | 0.33 ± 0.04 | 2.02 ± 0.01 | L272R/ΔG273 |
| 177 | 0.73 ± 0.14 | 0.48 ± 0.10 | M293I/ΔK294 |
| 178 | 0.63 ± 0.03 | 1.52 ± 0.07 | A266G/ΔS267 |
| **-6 bp** | | | |
| 1 | <0.01 | <0.01 | n.d. |
| 2 | <0.01 | <0.01 | n.d. |
| 3 | <0.01 | <0.01 | n.d. |
| 4 | <0.01 | <0.01 | n.d. |
| 5 | <0.01 | <0.01 | n.d. |
| 6 | <0.01 | <0.01 | n.d. |
| 7 | <0.01 | <0.01 | n.d. |
| 8 | <0.01 | <0.01 | n.d. |
| 9 | <0.01 | <0.01 | n.d. |
| 10 | 0.41 ± 0.02 | 1.41 ± 0.07 | ΔL262 |
| 11 | <0.01 | <0.01 | n.d. |
| 12 | <0.01 | <0.01 | n.d. |
| 13 | 0.03 ± 0.01 | 0.01 ± 0.01 | n.d. |
| 14 | <0.01 | 0.07 ± 0.02 | n.d. |
| 15 | <0.01 | <0.01 | n.d. |
| 16 | <0.01 | <0.01 | n.d. |
| 17 | <0.01 | <0.01 | n.d. |
| 18 | <0.01 | <0.01 | n.d. |
| 19 | 0.02 ± 0.01 | <0.01 | n.d. |
| 20 | 0.12 ± 0.02 | 0.53 ± 0.06 | n.d. |
| 21 | <0.01 | <0.01 | n.d. |
| 22 | <0.01 | <0.01 | n.d. |
| 23 | <0.01 | 0.07 ± 0.01 | n.d. |
| 24 | 0.07 ± 0.01 | 0.03 ± 0.01 | n.d. |
| 25 | <0.01 | <0.01 | n.d. |
| 26 | <0.01 | <0.01 | n.d. |
| 27 | <0.01 | <0.01 | n.d. |
| 28 | <0.01 | <0.01 | n.d. |
| 29 | <0.01 | <0.01 | n.d. |
| 30 | <0.01 | <0.01 | n.d. |
| 31 | <0.01 | <0.01 | n.d. |

(Table S9 continued)

| Variant | Relative activity | | Protein mutation |
|---|---|---|---|
| | Paraoxon | 4-NPB | |
| 32 | <0.01 | <0.01 | n.d. |
| 33 | <0.01 | <0.01 | n.d. |
| 34 | <0.01 | <0.01 | n.d. |
| 35 | 0.05 ± 0.01 | 0.08 ± 0.03 | n.d. |
| 36 | <0.01 | <0.01 | n.d. |
| 37 | 0.20 ± 0.01 | 0.21 ± 0.01 | n.d. |
| 38 | 0.93 ± 0.08 | 1.05 ± 0.11 | ∆D35R36 |
| 39 | <0.01 | <0.01 | n.d. |
| 40 | <0.01 | <0.01 | n.d. |
| 41 | <0.01 | <0.01 | n.d. |
| 42 | <0.01 | <0.01 | n.d. |
| 43 | <0.01 | <0.01 | ∆I157E158 |
| 44 | <0.01 | <0.01 | E219V/∆G220L221 |
| 45 | <0.01 | <0.01 | n.d. |
| 46 | <0.01 | <0.01 | n.d. |
| 47 | <0.01 | <0.01 | n.d. |
| 48 | <0.01 | <0.01 | n.d. |
| 49 | <0.01 | <0.01 | n.d. |
| 50 | <0.01 | <0.01 | n.d. |
| 51 | 0.01 ± 0.01 | <0.01 | n.d. |
| 52 | <0.01 | <0.01 | n.d. |
| 53 | <0.01 | <0.01 | n.d. |
| 54 | <0.01 | <0.01 | n.d. |
| 55 | <0.01 | 0.05 ± 0.01 | n.d. |
| 56 | <0.01 | <0.01 | n.d. |
| 57 | <0.01 | <0.01 | n.d. |
| 58 | <0.01 | <0.01 | n.d. |
| 59 | <0.01 | <0.01 | n.d. |
| 60 | <0.01 | 0.48 ± 0.09 | n.d. |
| 61 | <0.01 | <0.01 | n.d. |
| 62 | <0.01 | <0.01 | n.d. |
| 63 | <0.01 | <0.01 | n.d. |
| 64 | 0.94 ± 0.10 | 1.52 ± 0.02 | V310G/T311N/S311a |
| 65 | <0.01 | <0.01 | n.d. |
| 66 | <0.01 | <0.01 | n.d. |
| 67 | <0.01 | <0.01 | n.d. |
| 68 | <0.01 | <0.01 | n.d. |
| 69 | <0.01 | <0.01 | n.d. |
| 70 | <0.01 | 0.15 ± 0.01 | n.d. |
| 71 | <0.01 | <0.01 | n.d. |
| 72 | <0.01 | <0.01 | n.d. |
| 73 | <0.01 | 0.12 ± 0.04 | n.d. |
| 74 | <0.01 | <0.01 | n.d. |
| 75 | <0.01 | <0.01 | n.d. |

(Table S9 continued)

| Variant | Relative activity | | Protein mutation |
|---|---|---|---|
| | Paraoxon | 4-NPB | |
| 76 | <0.01 | <0.01 | n.d. |
| 77 | <0.01 | <0.01 | n.d. |
| 78 | <0.01 | <0.01 | n.d. |
| 79 | <0.01 | <0.01 | n.d. |
| 80 | 0.06 ± 0.01 | 0.77 ± 0.03 | n.d. |
| 81 | <0.01 | <0.01 | n.d. |
| 82 | <0.01 | <0.01 | n.d. |
| 83 | <0.01 | <0.01 | n.d. |
| 84 | <0.01 | <0.01 | n.d. |
| 85 | <0.01 | <0.01 | n.d. |
| 86 | <0.01 | <0.01 | n.d. |
| 87 | <0.01 | <0.01 | n.d. |
| 88 | 0.14 ± 0.01 | 0.45 ± 0.05 | n.d. |
| 89 | <0.01 | <0.01 | n.d. |
| 90 | <0.01 | <0.01 | n.d. |
| 91 | <0.01 | <0.01 | n.d. |
| 92 | <0.01 | <0.01 | n.d. |
| 93 | <0.01 | <0.01 | n.d. |
| 94 | <0.01 | <0.01 | n.d. |
| 95 | 0.05 ± 0.01 | 0.64 ± 0.07 | n.d. |
| 96 | <0.01 | <0.01 | n.d. |
| 97 | 0.03 ± 0.01 | <0.01 | n.d. |
| 98 | <0.01 | <0.01 | ΔV198T199 |
| 99 | 0.03 ± 0.01 | 0.11 ± 0.01 | ΔW277Q278 |
| 100 | <0.01 | <0.01 | n.d. |
| 101 | <0.01 | <0.01 | ΔA83V84 |
| 102 | <0.01 | <0.01 | n.d. |
| 103 | <0.01 | <0.01 | n.d. |
| 104 | <0.01 | <0.01 | n.d. |
| 105 | <0.01 | <0.01 | n.d. |
| 106 | <0.01 | <0.01 | n.d. |
| 107 | <0.01 | <0.01 | n.d. |
| 108 | <0.01 | <0.01 | n.d. |
| 109 | <0.01 | <0.01 | n.d. |
| 110 | <0.01 | <0.01 | n.d. |
| 111 | <0.01 | <0.01 | n.d. |
| 112 | <0.01 | <0.01 | n.d. |
| 113 | <0.01 | <0.01 | n.d. |
| 114 | 0.07 ± 0.01 | 1.23 ± 0.08 | n.d. |
| 115 | <0.01 | <0.01 | n.d. |
| 116 | <0.01 | <0.01 | n.d. |
| 117 | <0.01 | <0.01 | n.d. |
| 118 | <0.01 | <0.01 | n.d. |
| 119 | <0.01 | <0.01 | n.d. |

(Table S9 continued)

| Variant | Relative activity | | Protein mutation |
|---|---|---|---|
| | **Paraoxon** | **4-NPB** | |
| 120 | 0.21 ± 0.04 | 0.16 ± 0.05 | ΔE48A49 |
| 121 | <0.01 | <0.01 | n.d. |
| 122 | <0.01 | <0.01 | n.d. |
| 123 | <0.01 | <0.01 | n.d. |
| 124 | <0.01 | <0.01 | n.d. |
| 125 | <0.01 | <0.01 | n.d. |
| 126 | <0.01 | <0.01 | n.d. |
| 127 | <0.01 | <0.01 | n.d. |
| 128 | <0.01 | <0.01 | n.d. |
| 129 | 0.16 ± 0.01 | 1.24 ± 0.01 | n.d. |
| 130 | <0.01 | <0.01 | n.d. |
| 131 | <0.01 | <0.01 | n.d. |
| 132 | <0.01 | <0.01 | n.d. |
| 133 | <0.01 | <0.01 | n.d. |
| 134 | <0.01 | <0.01 | n.d. |
| 135 | <0.01 | <0.01 | n.d. |
| 136 | 0.02 ± 0.01 | <0.01 | n.d. |
| 137 | <0.01 | <0.01 | n.d. |
| 138 | <0.01 | <0.01 | n.d. |
| 139 | <0.01 | <0.01 | n.d. |
| 140 | <0.01 | <0.01 | n.d. |
| 141 | <0.01 | <0.01 | n.d. |
| 142 | <0.01 | <0.01 | n.d. |
| 143 | <0.01 | <0.01 | n.d. |
| 144 | <0.01 | <0.01 | n.d. |
| 145 | <0.01 | <0.01 | n.d. |
| 146 | <0.01 | <0.01 | n.d. |
| 147 | <0.01 | <0.01 | n.d. |
| 148 | <0.01 | <0.01 | n.d. |
| 149 | <0.01 | <0.01 | n.d. |
| 150 | <0.01 | <0.01 | n.d. |
| 151 | <0.01 | <0.01 | n.d. |
| 152 | <0.01 | <0.01 | n.d. |
| 153 | <0.01 | <0.01 | n.d. |
| 154 | <0.01 | <0.01 | n.d. |
| 155 | <0.01 | <0.01 | n.d. |
| 156 | <0.01 | <0.01 | n.d. |
| 157 | <0.01 | <0.01 | n.d. |
| 158 | <0.01 | <0.01 | n.d. |
| 159 | <0.01 | <0.01 | n.d. |
| 160 | <0.01 | <0.01 | n.d. |
| 161 | <0.01 | <0.01 | n.d. |
| 162 | <0.01 | 0.03 ± 0.01 | n.d. |
| 163 | <0.01 | <0.01 | n.d. |

(Table S9 continued)

| Variant | Relative activity | | Protein mutation |
|---|---|---|---|
| | **Paraoxon** | **4-NPB** | |
| 164 | <0.01 | <0.01 | n.d. |
| 165 | 0.53 ± 0.12 | 0.24 ± 0.07 | Q206H/ΔR207G208 |
| 166 | 0.16 ± 0.01 | 1.42 ± 0.07 | S269F/ΔA270L271 |
| 167 | <0.01 | <0.01 | n.d. |
| 168 | <0.01 | <0.01 | n.d. |
| 169 | <0.01 | <0.01 | n.d. |
| 170 | <0.01 | <0.01 | n.d. |
| 171 | <0.01 | <0.01 | n.d. |
| 172 | 0.11 ± 0.01 | 0.07 ± 0.01 | n.d. |
| 173 | <0.01 | <0.01 | n.d. |
| 174 | <0.01 | <0.01 | n.d. |
| 175 | <0.01 | <0.01 | n.d. |
| 176 | <0.01 | <0.01 | n.d. |
| 177 | <0.01 | <0.01 | n.d. |
| 178 | <0.01 | <0.01 | n.d. |
| **-9 bp** | | | |
| 1 | <0.01 | <0.01 | n.d. |
| 2 | <0.01 | <0.01 | n.d. |
| 3 | <0.01 | <0.01 | n.d. |
| 4 | <0.01 | <0.01 | n.d. |
| 5 | <0.01 | <0.01 | n.d. |
| 6 | <0.01 | <0.01 | n.d. |
| 7 | <0.01 | <0.01 | n.d. |
| 8 | <0.01 | <0.01 | n.d. |
| 9 | 0.05 ± 0.01 | 0.68 ± 0.01 | n.d. |
| 10 | <0.01 | <0.01 | n.d. |
| 11 | <0.01 | <0.01 | n.d. |
| 12 | <0.01 | <0.01 | n.d. |
| 13 | <0.01 | <0.01 | n.d. |
| 14 | 0.08 ± 0.01 | 3.95 ± 0.16 | ΔG261L262E263 |
| 15 | <0.01 | <0.01 | n.d. |
| 16 | <0.01 | <0.01 | n.d. |
| 17 | <0.01 | <0.01 | n.d. |
| 18 | 0.02 ± 0.01 | <0.01 | n.d. |
| 19 | <0.01 | <0.01 | n.d. |
| 20 | <0.01 | <0.01 | n.d. |
| 21 | <0.01 | <0.01 | n.d. |
| 22 | <0.01 | <0.01 | n.d. |
| 23 | <0.01 | <0.01 | n.d. |
| 24 | <0.01 | <0.01 | n.d. |
| 25 | <0.01 | 0.18 ± 0.02 | n.d. |
| 26 | <0.01 | <0.01 | n.d. |
| 27 | <0.01 | <0.01 | n.d. |
| 28 | <0.01 | <0.01 | n.d. |

(Table S9 continued)

| Variant | Relative activity | | Protein mutation |
|---|---|---|---|
| | Paraoxon | 4-NPB | |
| 29 | 0.09 ± 0.01 | 2.61 ± 0.02 | ∆A268S269A270 |
| 30 | <0.01 | <0.01 | n.d. |
| 31 | <0.01 | <0.01 | n.d. |
| 32 | <0.01 | <0.01 | n.d. |
| 33 | <0.01 | 0.05 ± 0.01 | n.d. |
| 34 | <0.01 | <0.01 | n.d. |
| 35 | 0.02 ± 0.01 | 0.02 ± 0.01 | n.d. |
| 36 | 0.33 ± 0.01 | 1.33 ± 0.06 | n.d. |
| 37 | <0.01 | <0.01 | n.d. |
| 38 | <0.01 | <0.01 | n.d. |
| 39 | <0.01 | <0.01 | n.d. |
| 40 | <0.01 | <0.01 | n.d. |
| 41 | <0.01 | 0.03 ± 0.01 | n.d. |
| 42 | <0.01 | <0.01 | ∆H201T202A203 |
| 43 | <0.01 | <0.01 | ∆A188R189A190 |
| 44 | <0.01 | 0.25 ± 0.01 | ∆S276W277Q278 |
| 45 | 0.10 ± 0.01 | 0.38 ± 0.01 | ∆I341P342Q343 |
| 46 | <0.01 | <0.01 | N321K/∆P322D323G324 |
| 47 | <0.01 | <0.01 | n.d. |
| 48 | <0.01 | <0.01 | n.d. |
| 49 | <0.01 | <0.01 | n.d. |
| 50 | <0.01 | <0.01 | n.d. |
| 51 | 0.11 ± 0.01 | 4.17 ± 0.01 | ∆G261L262E263 |
| 52 | <0.01 | <0.01 | n.d. |
| 53 | <0.01 | <0.01 | n.d. |
| 54 | <0.01 | <0.01 | n.d. |
| 55 | <0.01 | <0.01 | n.d. |
| 56 | <0.01 | <0.01 | n.d. |
| 57 | <0.01 | <0.01 | n.d. |
| 58 | <0.01 | <0.01 | n.d. |
| 59 | <0.01 | <0.01 | n.d. |
| 60 | 0.03 ± 0.01 | <0.01 | n.d. |
| 61 | <0.01 | <0.01 | n.d. |
| 62 | 0.24 ± 0.06 | 0.68 ± 0.18 | n.d. |
| 63 | <0.01 | <0.01 | n.d. |
| 64 | <0.01 | <0.01 | n.d. |
| 65 | <0.01 | <0.01 | n.d. |
| 66 | <0.01 | <0.01 | n.d. |
| 67 | <0.01 | <0.01 | n.d. |
| 68 | <0.01 | <0.01 | n.d. |
| 69 | 0.03 ± 0.01 | 0.03 ± 0.01 | n.d. |
| 70 | <0.01 | <0.01 | n.d. |
| 71 | <0.01 | <0.01 | n.d. |
| 72 | 0.68 ± 0.02 | 0.81 ± 0.01 | ∆S75R76K77 |

(Table S9 continued)

| Variant | Relative activity | | Protein mutation |
|---|---|---|---|
| | Paraoxon | 4-NPB | |
| 73 | <0.01 | <0.01 | n.d. |
| 74 | <0.01 | <0.01 | n.d. |
| 75 | <0.01 | <0.01 | n.d. |
| 76 | <0.01 | <0.01 | n.d. |
| 77 | 0.03 ± 0.01 | <0.01 | n.d. |
| 78 | <0.01 | <0.01 | n.d. |
| 79 | <0.01 | <0.01 | n.d. |
| 80 | 0.10 ± 0.01 | 0.84 ± 0.07 | n.d. |
| 81 | <0.01 | <0.01 | n.d. |
| 82 | <0.01 | <0.01 | n.d. |
| 83 | 0.08 ± 0.01 | 0.94 ± 0.04 | n.d. |
| 84 | 0.05 ± 0.01 | <0.01 | n.d. |
| 85 | <0.01 | <0.01 | n.d. |
| 86 | <0.01 | <0.01 | n.d. |
| 87 | <0.01 | <0.01 | n.d. |
| 88 | <0.01 | <0.01 | n.d. |
| 89 | <0.01 | <0.01 | n.d. |
| 90 | <0.01 | <0.01 | n.d. |
| 91 | <0.01 | <0.01 | n.d. |
| 92 | <0.01 | <0.01 | n.d. |
| 93 | <0.01 | <0.01 | n.d. |
| 94 | 1.31 ± 0.02 | 1.27 ± 0.07 | n.d. |
| 95 | <0.01 | <0.01 | n.d. |
| 96 | <0.01 | <0.01 | n.d. |
| 97 | 0.14 ± 0.01 | 5.02 ± 0.09 | ΔG261L262E263 |
| 98 | 0.08 ± 0.02 | 0.08 ± 0.03 | R91S/ΔA92A93G94 |
| 99 | <0.01 | <0.01 | frameshift |
| 100 | <0.01 | <0.01 | frameshift |
| 101 | <0.01 | <0.01 | n.d. |
| 102 | <0.01 | 0.32 ± 0.01 | n.d. |
| 103 | <0.01 | <0.01 | n.d. |
| 104 | <0.01 | <0.01 | n.d. |
| 105 | <0.01 | <0.01 | n.d. |
| 106 | <0.01 | <0.01 | n.d. |
| 107 | <0.01 | <0.01 | n.d. |
| 108 | <0.01 | <0.01 | n.d. |
| 109 | <0.01 | <0.01 | n.d. |
| 110 | <0.01 | <0.01 | n.d. |
| 111 | <0.01 | <0.01 | n.d. |
| 112 | <0.01 | <0.01 | n.d. |
| 113 | <0.01 | <0.01 | n.d. |
| 114 | <0.01 | <0.01 | n.d. |
| 115 | <0.01 | <0.01 | n.d. |
| 116 | <0.01 | <0.01 | n.d. |

(Table S9 continued)

| Variant | Relative activity | | Protein mutation |
|---|---|---|---|
| | Paraoxon | 4-NPB | |
| 117 | <0.01 | <0.01 | n.d. |
| 118 | 0.13 ± 0.01 | 0.16 ± 0.01 | n.d. |
| 119 | <0.01 | <0.01 | n.d. |
| 120 | <0.01 | <0.01 | n.d. |
| 121 | <0.01 | <0.01 | n.d. |
| 122 | <0.01 | <0.01 | n.d. |
| 123 | <0.01 | <0.01 | n.d. |
| 124 | <0.01 | <0.01 | n.d. |
| 125 | <0.01 | <0.01 | n.d. |
| 126 | <0.01 | <0.01 | n.d. |
| 127 | <0.01 | <0.01 | n.d. |
| 128 | 0.24 ± 0.01 | 2.41 ± 0.11 | ΔL272G273I274 |
| 129 | <0.01 | <0.01 | n.d. |
| 130 | 0.26 ± 0.01 | 1.99 ± 0.02 | ΔS269A270L271 |
| 131 | 0.14 ± 0.01 | 0.15 ± 0.01 | n.d. |
| 132 | 0.12 ± 0.01 | 0.12 ± 0.01 | n.d. |
| 133 | <0.01 | <0.01 | n.d. |
| 134 | <0.01 | <0.01 | n.d. |
| 135 | <0.01 | <0.01 | n.d. |
| 136 | <0.01 | 0.15 ± 0.01 | n.d. |
| 137 | <0.01 | <0.01 | n.d. |
| 138 | <0.01 | <0.01 | n.d. |
| 139 | <0.01 | <0.01 | n.d. |
| 140 | <0.01 | <0.01 | n.d. |
| 141 | <0.01 | <0.01 | n.d. |
| 142 | <0.01 | <0.01 | n.d. |
| 143 | <0.01 | <0.01 | n.d. |
| 144 | <0.01 | <0.01 | n.d. |
| 145 | <0.01 | <0.01 | n.d. |
| 146 | 0.06 ± 0.01 | 0.02 ± 0.01 | n.d. |
| 147 | <0.01 | <0.01 | n.d. |
| 148 | <0.01 | <0.01 | n.d. |
| 149 | <0.01 | <0.01 | n.d. |
| 150 | <0.01 | <0.01 | n.d. |
| 151 | 0.21 ± 0.01 | 4.55 ± 0.09 | ΔG261L262E263 |
| 152 | <0.01 | <0.01 | n.d. |
| 153 | <0.01 | <0.01 | n.d. |
| 154 | <0.01 | <0.01 | n.d. |
| 155 | <0.01 | <0.01 | n.d. |
| 156 | <0.01 | <0.01 | n.d. |
| 157 | <0.01 | <0.01 | n.d. |
| 158 | <0.01 | <0.01 | n.d. |
| 159 | <0.01 | <0.01 | n.d. |
| 160 | <0.01 | <0.01 | n.d. |

(Table S9 continued)

| Variant | Relative activity | | Protein mutation |
|---|---|---|---|
| | Paraoxon | 4-NPB | |
| 161 | <0.01 | <0.01 | n.d. |
| 162 | <0.01 | <0.01 | n.d. |
| 163 | <0.01 | <0.01 | n.d. |
| 164 | <0.01 | <0.01 | n.d. |
| 165 | <0.01 | <0.01 | n.d. |
| 166 | <0.01 | <0.01 | n.d. |
| 167 | <0.01 | <0.01 | n.d. |
| 168 | <0.01 | <0.01 | n.d. |
| 169 | <0.01 | <0.01 | n.d. |
| 170 | <0.01 | <0.01 | n.d. |
| 171 | <0.01 | <0.01 | n.d. |
| 172 | <0.01 | <0.01 | n.d. |
| 173 | <0.01 | <0.01 | n.d. |
| 174 | <0.01 | <0.01 | n.d. |
| 175 | <0.01 | <0.01 | n.d. |
| 176 | <0.01 | 0.26 ± 0.01 | n.d. |
| 177 | <0.01 | <0.01 | n.d. |
| 178 | <0.01 | <0.01 | n.d. |
| **+3 bp** | | | |
| 1 | <0.01 | 0.07 ± 0.01 | n.d. |
| 2 | 0.06 ± 0.01 | 0.05 ± 0.01 | n.d. |
| 3 | <0.01 | <0.01 | n.d. |
| 4 | 0.78 ± 0.01 | 2.09 ± 0.1 | S261a |
| 5 | <0.01 | 0.06 ± 0.01 | n.d. |
| 6 | <0.01 | <0.01 | n.d. |
| 7 | <0.01 | 0.07 ± 0.01 | n.d. |
| 8 | <0.01 | <0.01 | n.d. |
| 9 | <0.01 | <0.01 | n.d. |
| 10 | <0.01 | <0.01 | n.d. |
| 11 | 1.04 ± 0.01 | 1.39 ± 0.01 | Q339a |
| 12 | 0.95 ± 0.01 | 2.71 ± 0.01 | P261a |
| 13 | 0.95 ± 0.06 | 3.41 ± 0.18 | P259a |
| 14 | 1.41 ± 0.02 | 1.30 ± 0.04 | n.d. |
| 15 | 0.31 ± 0.04 | 0.27 ± 0.05 | A161a |
| 16 | 0.23 ± 0.05 | 0.23 ± 0.06 | n.d. |
| 17 | <0.01 | <0.01 | n.d. |
| 18 | <0.01 | 0.04 ± 0.01 | n.d. |
| 19 | <0.01 | <0.01 | n.d. |
| 20 | 0.78 ± 0.01 | 0.68 ± 0.01 | P43H/A43a |
| 21 | 0.15 ± 0.02 | 0.14 ± 0.02 | n.d. |
| 22 | <0.01 | <0.01 | n.d. |
| 23 | <0.01 | <0.01 | n.d. |
| 24 | <0.01 | <0.01 | n.d. |
| 25 | 1.24 ± 0.02 | 2.46 ± 0.04 | S319a |

(Table S9 continued)

| Variant | Relative activity | | Protein mutation |
|---|---|---|---|
| | Paraoxon | 4-NPB | |
| 26 | <0.01 | 0.03 ± 0.01 | frameshift |
| 27 | <0.01 | <0.01 | frameshift |
| 28 | <0.01 | <0.01 | frameshift |
| 29 | 0.18 ± 0.01 | 1.72 ± 0.02 | T311R/A311a |
| 30 | <0.01 | <0.01 | frameshift |
| 31 | <0.01 | <0.01 | n.d. |
| 32 | <0.01 | <0.01 | n.d. |
| 33 | 0.02 ± 0.01 | <0.01 | n.d. |
| 34 | <0.01 | 0.08 ± 0.02 | n.d. |
| 35 | <0.01 | <0.01 | n.d. |
| 36 | 0.90 ± 0.03 | 0.60 ± 0.10 | H42a |
| 37 | <0.01 | <0.01 | n.d. |
| 38 | <0.01 | <0.01 | n.d. |
| 39 | <0.01 | <0.01 | n.d. |
| 40 | <0.01 | <0.01 | n.d. |
| 41 | <0.01 | <0.01 | n.d. |
| 42 | <0.01 | <0.01 | n.d. |
| 43 | <0.01 | <0.01 | n.d. |
| 44 | 0.59 ± 0.02 | 0.62 ± 0.01 | K339M/Q339a |
| 45 | <0.01 | <0.01 | n.d. |
| 46 | <0.01 | <0.01 | n.d. |
| 47 | <0.01 | <0.01 | n.d. |
| 48 | <0.01 | <0.01 | n.d. |
| 49 | 0.38 ± 0.02 | 1.89 ± 0.06 | S311a |
| 50 | <0.01 | <0.01 | n.d. |
| 51 | <0.01 | <0.01 | n.d. |
| 52 | <0.01 | <0.01 | n.d. |
| 53 | <0.01 | <0.01 | n.d. |
| 54 | <0.01 | <0.01 | n.d. |
| 55 | 0.02 ± 0.01 | <0.01 | n.d. |
| 56 | <0.01 | 0.07 ± 0.01 | n.d. |
| 57 | <0.01 | <0.01 | n.d. |
| 58 | 0.03 ± 0.01 | 0.16 ± 0.03 | n.d. |
| 59 | 0.98 ± 0.01 | 0.55 ± 0.08 | T161K/P161a |
| 60 | <0.01 | <0.01 | n.d. |
| 61 | 0.02 ± 0.01 | 0.69 ± 0.05 | n.d. |
| 62 | <0.01 | <0.01 | n.d. |
| 63 | 0.02 ± 0.01 | <0.01 | n.d. |
| 64 | <0.01 | <0.01 | n.d. |
| 65 | <0.01 | <0.01 | n.d. |
| 66 | <0.01 | 0.04 ± 0.01 | n.d. |
| 67 | 1.16 ± 0.07 | 1.24 ± 0.05 | E263V/Q263a |
| 68 | <0.01 | <0.01 | n.d. |
| 69 | <0.01 | 0.09 ± 0.01 | n.d. |

(Table S9 continued)

| Variant | Relative activity | | Protein mutation |
|---|---|---|---|
| | Paraoxon | 4-NPB | |
| 70 | <0.01 | <0.01 | n.d. |
| 71 | <0.01 | <0.01 | n.d. |
| 72 | <0.01 | <0.01 | n.d. |
| 73 | <0.01 | 0.03 ± 0.01 | n.d. |
| 74 | <0.01 | <0.01 | n.d. |
| 75 | <0.01 | <0.01 | n.d. |
| 76 | 1.01 ± 0.04 | 0.62 ± 0.09 | A242V |
| 77 | <0.01 | <0.01 | n.d. |
| 78 | 0.26 ± 0.07 | 0.25 ± 0.05 | N90a |
| 79 | 1.58 ± 0.07 | 1.72 ± 0.01 | V339a |
| 80 | 0.37 ± 0.08 | 0.50 ± 0.1 | D142a |
| 81 | <0.01 | <0.01 | n.d. |
| 82 | 1.79 ± 0.05 | 1.43 ± 0.01 | K362a |
| 83 | <0.01 | <0.01 | n.d. |
| 84 | <0.01 | <0.01 | n.d. |
| 85 | <0.01 | <0.01 | n.d. |
| 86 | <0.01 | <0.01 | n.d. |
| 87 | <0.01 | <0.01 | n.d. |
| 88 | <0.01 | <0.01 | n.d. |
| 89 | <0.01 | <0.01 | n.d. |
| 90 | <0.01 | <0.01 | n.d. |
| 91 | 0.03 ± 0.01 | <0.01 | n.d. |
| 92 | <0.01 | <0.01 | n.d. |
| 93 | <0.01 | <0.01 | n.d. |
| 94 | 1.33 ± 0.07 | 1.25 ± 0.06 | P35a |
| 95 | <0.01 | <0.01 | n.d. |
| 96 | 1.00 ± 0.06 | 1.18 ± 0.01 | I262a |
| 97 | <0.01 | <0.01 | n.d. |
| 98 | 0.35 ± 0.05 | 0.31 ± 0.01 | T311R/S311a |
| 99 | <0.01 | <0.01 | n.d. |
| 100 | <0.01 | <0.01 | n.d. |
| 101 | <0.01 | <0.01 | n.d. |
| 102 | <0.01 | <0.01 | frameshift |
| 103 | <0.01 | <0.01 | frameshift |
| 104 | <0.01 | <0.01 | I183a |
| 105 | <0.01 | <0.01 | frameshift |
| 106 | <0.01 | <0.01 | n.d. |
| 107 | <0.01 | <0.01 | n.d. |
| 108 | <0.01 | 0.02 ± 0.01 | n.d. |
| 109 | <0.01 | <0.01 | n.d. |
| 110 | <0.01 | 0.04 ± 0.01 | n.d. |
| 111 | <0.01 | <0.01 | n.d. |
| 112 | <0.01 | <0.01 | n.d. |
| 113 | 0.07 ± 0.01 | 1.23 ± 0.06 | n.d. |

(Table S9 continued)

| Variant | Relative activity | | Protein mutation |
|---|---|---|---|
| | Paraoxon | 4-NPB | |
| 114 | <0.01 | <0.01 | n.d. |
| 115 | 0.37 ± 0.02 | 4.12 ± 0.11 | A259G/S259a |
| 116 | <0.01 | 0.02 ± 0.01 | n.d. |
| 117 | <0.01 | <0.01 | n.d. |
| 118 | <0.01 | <0.01 | n.d. |
| 119 | <0.01 | <0.01 | n.d. |
| 120 | <0.01 | <0.01 | n.d. |
| 121 | <0.01 | <0.01 | n.d. |
| 122 | <0.01 | <0.01 | n.d. |
| 123 | <0.01 | <0.01 | n.d. |
| 124 | <0.01 | 0.08 ± 0.01 | S231R/F231a |
| 125 | <0.01 | <0.01 | n.d. |
| 126 | <0.01 | <0.01 | n.d. |
| 127 | 0.08 ± 0.02 | <0.01 | n.d. |
| 128 | 0.08 ± 0.01 | 0.02 ± 0.01 | n.d. |
| 129 | 0.44 ± 0.02 | 1.14 ± 0.02 | V309a |
| 130 | <0.01 | <0.01 | n.d. |
| 131 | 0.14 ± 0.02 | 0.18 ± 0.06 | n.d. |
| 132 | <0.01 | <0.01 | n.d. |
| 133 | <0.01 | <0.01 | n.d. |
| 134 | <0.01 | <0.01 | n.d. |
| 135 | <0.01 | <0.01 | n.d. |
| 136 | 0.37 ± 0.06 | 0.16 ± 0.01 | I291a |
| 137 | <0.01 | <0.01 | n.d. |
| 138 | <0.01 | <0.01 | n.d. |
| 139 | <0.01 | 0.03 ± 0.01 | n.d. |
| 140 | 0.02 ± 0.01 | <0.01 | n.d. |
| 141 | <0.01 | <0.01 | n.d. |
| 142 | <0.01 | <0.01 | n.d. |
| 143 | <0.01 | <0.01 | n.d. |
| 144 | <0.01 | <0.01 | n.d. |
| 145 | 1.16 ± 0.03 | 1.09 ± 0.11 | V339a |
| 146 | <0.01 | <0.01 | n.d. |
| 147 | 1.25 ± 0.04 | 1.31 ± 0.07 | F35a |
| 148 | <0.01 | <0.01 | n.d. |
| 149 | <0.01 | <0.01 | n.d. |
| 150 | <0.01 | <0.01 | n.d. |
| 151 | <0.01 | <0.01 | n.d. |
| 152 | <0.01 | <0.01 | n.d. |
| 153 | 1.09 ± 0.01 | 0.91 ± 0.14 | P35a |
| 154 | <0.01 | <0.01 | n.d. |
| 155 | <0.01 | 0.05 ± 0.02 | n.d. |
| 156 | <0.01 | <0.01 | n.d. |
| 157 | <0.01 | <0.01 | n.d. |

(Table S9 continued)

| Variant | Relative activity | | Protein mutation |
|---|---|---|---|
| | Paraoxon | 4-NPB | |
| 158 | 1.09 ± 0.04 | 1.14 ± 0.05 | V339a |
| 159 | 0.84 ± 0.09 | 0.49 ± 0.01 | E145A |
| 160 | 0.02 ± 0.01 | 0.31 ± 0.04 | n.d. |
| 161 | <0.01 | <0.01 | n.d. |
| 162 | <0.01 | <0.01 | n.d. |
| 163 | <0.01 | <0.01 | n.d. |
| 164 | <0.01 | <0.01 | n.d. |
| 165 | <0.01 | <0.01 | n.d. |
| 166 | <0.01 | <0.01 | n.d. |
| 167 | <0.01 | <0.01 | n.d. |
| 168 | <0.01 | <0.01 | n.d. |
| 169 | <0.01 | <0.01 | n.d. |
| 170 | <0.01 | 0.13 ± 0.04 | n.d. |
| 171 | <0.01 | <0.01 | n.d. |
| 172 | <0.01 | <0.01 | n.d. |
| 173 | <0.01 | <0.01 | n.d. |
| 174 | <0.01 | <0.01 | n.d. |
| 175 | <0.01 | <0.01 | n.d. |
| 176 | <0.01 | <0.01 | n.d. |
| 177 | 0.47 ± 0.02 | 0.82 ± 0.04 | F310a |
| 178 | <0.01 | <0.01 | n.d. |
| **+6 bp** | | | |
| 1 | <0.01 | <0.01 | n.d. |
| 2 | <0.01 | <0.01 | n.d. |
| 3 | <0.01 | <0.01 | n.d. |
| 4 | 0.34 ± 0.01 | 1.56 ± 0.01 | n.d. |
| 5 | <0.01 | <0.01 | n.d. |
| 6 | <0.01 | <0.01 | n.d. |
| 7 | <0.01 | <0.01 | n.d. |
| 8 | <0.01 | <0.01 | n.d. |
| 9 | <0.01 | <0.01 | n.d. |
| 10 | <0.01 | <0.01 | n.d. |
| 11 | <0.01 | <0.01 | n.d. |
| 12 | <0.01 | <0.01 | n.d. |
| 13 | <0.01 | <0.01 | R311aV311b |
| 14 | <0.01 | <0.01 | A69aC69b |
| 15 | <0.01 | <0.01 | F155aI155b |
| 16 | <0.01 | <0.01 | H230R/Q230aY230b |
| 17 | <0.01 | <0.01 | frameshift |
| 18 | <0.01 | <0.01 | n.d. |
| 19 | <0.01 | <0.01 | n.d. |
| 20 | <0.01 | <0.01 | n.d. |
| 21 | <0.01 | <0.01 | n.d. |
| 22 | <0.01 | <0.01 | n.d. |

(Table S9 continued)

| Variant | Relative activity | | Protein mutation |
|---|---|---|---|
| | Paraoxon | 4-NPB | |
| 23 | 0.13 ± 0.01 | 3.03 ± 0.13 | Q271aR271b |
| 24 | <0.01 | <0.01 | n.d. |
| 25 | <0.01 | <0.01 | n.d. |
| 26 | <0.01 | 0.12 ± 0.04 | n.d. |
| 27 | <0.01 | <0.01 | n.d. |
| 28 | 0.16 ± 0.03 | 0.87 ± 0.14 | n.d. |
| 29 | 0.58 ± 0.01 | 0.47 ± 0.01 | n.d. |
| 30 | <0.01 | <0.01 | n.d. |
| 31 | <0.01 | <0.01 | n.d. |
| 32 | <0.01 | <0.01 | n.d. |
| 33 | <0.01 | <0.01 | n.d. |
| 34 | <0.01 | <0.01 | n.d. |
| 35 | <0.01 | <0.01 | n.d. |
| 36 | 0.09 ± 0.01 | 3.13 ± 0.14 | Y275aG275b |
| 37 | 0.21 ± 0.01 | 2.65 ± 0.01 | P261aC261b |
| 38 | 1.50 ± 0.07 | 1.43 ± 0.08 | K35aC35b |
| 39 | <0.01 | <0.01 | n.d. |
| 40 | 0.03 ± 0.01 | <0.01 | n.d. |
| 41 | <0.01 | <0.01 | n.d. |
| 42 | <0.01 | <0.01 | n.d. |
| 43 | <0.01 | <0.01 | n.d. |
| 44 | <0.01 | <0.01 | n.d. |
| 45 | <0.01 | <0.01 | n.d. |
| 46 | <0.01 | <0.01 | n.d. |
| 47 | <0.01 | <0.01 | n.d. |
| 48 | <0.01 | <0.01 | n.d. |
| 49 | <0.01 | <0.01 | n.d. |
| 50 | <0.01 | <0.01 | n.d. |
| 51 | 0.16 ± 0.01 | 0.13 ± 0.01 | n.d. |
| 52 | 0.03 ± 0.01 | <0.01 | n.d. |
| 53 | <0.01 | <0.01 | n.d. |
| 54 | <0.01 | 0.11 ± 0.02 | n.d. |
| 55 | <0.01 | <0.01 | n.d. |
| 56 | <0.01 | <0.01 | n.d. |
| 57 | <0.01 | <0.01 | n.d. |
| 58 | <0.01 | <0.01 | n.d. |
| 59 | 0.20 ± 0.01 | 0.11 ± 0.01 | n.d. |
| 60 | <0.01 | <0.01 | n.d. |
| 61 | <0.01 | <0.01 | n.d. |
| 62 | <0.01 | <0.01 | n.d. |
| 63 | 0.48 ± 0.11 | 0.36 ± 0.1 | L291aS291b |
| 64 | <0.01 | <0.01 | n.d. |
| 65 | <0.01 | 0.09 ± 0.01 | n.d. |
| 66 | <0.01 | <0.01 | n.d. |

(Table S9 continued)

| Variant | Relative activity | | Protein mutation |
|---|---|---|---|
| | Paraoxon | 4-NPB | |
| 67 | 0.12 ± 0.03 | 0.14 ± 0.04 | n.d. |
| 68 | <0.01 | <0.01 | n.d. |
| 69 | 0.16 ± 0.01 | 2.92 ± 0.14 | Y268aR268b |
| 70 | <0.01 | <0.01 | n.d. |
| 71 | <0.01 | 0.02 ± 0.01 | n.d. |
| 72 | <0.01 | <0.01 | n.d. |
| 73 | <0.01 | <0.01 | n.d. |
| 74 | <0.01 | <0.01 | n.d. |
| 75 | <0.01 | <0.01 | n.d. |
| 76 | 0.08 ± 0.02 | 0.03 ± 0.01 | n.d. |
| 77 | <0.01 | <0.01 | n.d. |
| 78 | 0.03 ± 0.01 | 0.07 ± 0.02 | n.d. |
| 79 | <0.01 | <0.01 | n.d. |
| 80 | <0.01 | <0.01 | n.d. |
| 81 | <0.01 | 0.08 ± 0.01 | n.d. |
| 82 | 1.60 ± 0.01 | 1.59 ± 0.06 | F34aN34b |
| 83 | <0.01 | <0.01 | n.d. |
| 84 | <0.01 | <0.01 | n.d. |
| 85 | 1.26 ± 0.06 | 1.01 ± 0.07 | T37aI37b |
| 86 | <0.01 | <0.01 | n.d. |
| 87 | 0.86 ± 0.05 | 1.90 ± 0.01 | n.d. |
| 88 | 0.30 ± 0.01 | 1.33 ± 0.02 | n.d. |
| 89 | <0.01 | <0.01 | n.d. |
| 90 | <0.01 | <0.01 | frameshift |
| 91 | <0.01 | 0.59 ± 0.03 | S313aF313b |
| 92 | <0.01 | <0.01 | E339a-Stop |
| 93 | <0.01 | <0.01 | D168aL168b |
| 94 | <0.01 | <0.01 | n.d. |
| 95 | <0.01 | <0.01 | n.d. |
| 96 | <0.01 | <0.01 | n.d. |
| 97 | 0.42 ± 0.01 | 0.57 ± 0.02 | R342aV342b |
| 98 | 0.18 ± 0.04 | 0.17 ± 0.03 | n.d. |
| 99 | 0.04 ± 0.01 | <0.01 | n.d. |
| 100 | 0.79 ± 0.10 | 2.62 ± 0.06 | G266aR266b |
| 101 | 1.35 ± 0.03 | 1.48 ± 0.02 | n.d. |
| 102 | <0.01 | <0.01 | n.d. |
| 103 | <0.01 | <0.01 | n.d. |
| 104 | <0.01 | <0.01 | n.d. |
| 105 | 0.30 ± 0.08 | 0.35 ± 0.10 | I44R/Q44aF44b |
| 106 | <0.01 | <0.01 | n.d. |
| 107 | <0.01 | <0.01 | n.d. |
| 108 | <0.01 | <0.01 | n.d. |
| 109 | <0.01 | <0.01 | n.d. |
| 110 | <0.01 | <0.01 | n.d. |

(Table S9 continued)

| Variant | Relative activity | | Protein mutation |
|---|---|---|---|
| | Paraoxon | 4-NPB | |
| 111 | <0.01 | <0.01 | n.d. |
| 112 | <0.01 | <0.01 | n.d. |
| 113 | 0.22 ± 0.05 | 0.68 ± 0.13 | T238aA238b |
| 114 | <0.01 | <0.01 | n.d. |
| 115 | <0.01 | <0.01 | n.d. |
| 116 | 0.10 ± 0.01 | 0.14 ± 0.02 | n.d. |
| 117 | <0.01 | <0.01 | n.d. |
| 118 | <0.01 | <0.01 | n.d. |
| 119 | <0.01 | <0.01 | n.d. |
| 120 | <0.01 | <0.01 | n.d. |
| 121 | <0.01 | 0.08 ± 0.01 | n.d. |
| 122 | <0.01 | <0.01 | n.d. |
| 123 | 0.40 ± 0.10 | 0.45 ± 0.13 | I37M/L37aS37b |
| 124 | <0.01 | <0.01 | n.d. |
| 125 | <0.01 | <0.01 | n.d. |
| 126 | 0.02 ± 0.01 | <0.01 | n.d. |
| 127 | <0.01 | <0.01 | n.d. |
| 128 | <0.01 | <0.01 | n.d. |
| 129 | <0.01 | <0.01 | n.d. |
| 130 | <0.01 | <0.01 | n.d. |
| 131 | <0.01 | <0.01 | n.d. |
| 132 | <0.01 | <0.01 | n.d. |
| 133 | 0.18 ± 0.04 | 0.15 ± 0.05 | S179aG179b |
| 134 | 0.56 ± 0.03 | 1.51 ± 0.09 | G261aR261b |
| 135 | <0.01 | <0.01 | n.d. |
| 136 | <0.01 | <0.01 | n.d. |
| 137 | <0.01 | <0.01 | n.d. |
| 138 | <0.01 | <0.01 | n.d. |
| 139 | <0.01 | <0.01 | n.d. |
| 140 | <0.01 | <0.01 | n.d. |
| 141 | <0.01 | <0.01 | n.d. |
| 142 | <0.01 | <0.01 | n.d. |
| 143 | 0.18 ± 0.01 | 0.19 ± 0.01 | n.d. |
| 144 | <0.01 | <0.01 | n.d. |
| 145 | <0.01 | <0.01 | n.d. |
| 146 | <0.01 | <0.01 | n.d. |
| 147 | <0.01 | <0.01 | n.d. |
| 148 | <0.01 | <0.01 | n.d. |
| 149 | <0.01 | <0.01 | n.d. |
| 150 | 0.02 ± 0.01 | <0.01 | n.d. |
| 151 | <0.01 | <0.01 | n.d. |
| 152 | <0.01 | <0.01 | n.d. |
| 153 | <0.01 | <0.01 | n.d. |
| 154 | <0.01 | <0.01 | n.d. |

(Table S9 continued)

| Variant | Relative activity | | Protein mutation |
| --- | --- | --- | --- |
| | Paraoxon | 4-NPB | |
| 155 | <0.01 | <0.01 | n.d. |
| 156 | <0.01 | <0.01 | n.d. |
| 157 | <0.01 | <0.01 | n.d. |
| 158 | <0.01 | <0.01 | n.d. |
| 159 | <0.01 | <0.01 | n.d. |
| 160 | <0.01 | 0.14 ± 0.05 | n.d. |
| 161 | 0.25 ± 0.04 | 0.20 ± 0.05 | C289aS289b |
| 162 | 0.89 ± 0.01 | 1.06 ± 0.01 | C45aP45b |
| 163 | <0.01 | <0.01 | n.d. |
| 164 | <0.01 | <0.01 | n.d. |
| 165 | <0.01 | <0.01 | n.d. |
| 166 | <0.01 | <0.01 | n.d. |
| 167 | <0.01 | <0.01 | n.d. |
| 168 | 1.13 ± 0.11 | 1.00 ± 0.03 | F35aH35b |
| 169 | <0.01 | <0.01 | n.d. |
| 170 | <0.01 | <0.01 | n.d. |
| 171 | <0.01 | <0.01 | n.d. |
| 172 | <0.01 | <0.01 | n.d. |
| 173 | <0.01 | <0.01 | n.d. |
| 174 | <0.01 | <0.01 | n.d. |
| 175 | <0.01 | <0.01 | n.d. |
| 176 | <0.01 | <0.01 | n.d. |
| 177 | <0.01 | <0.01 | n.d. |
| 178 | <0.01 | 0.27 ± 0.01 | n.d. |
| **+9 bp** | | | |
| 1 | <0.01 | 0.09 ± 0.01 | n.d. |
| 2 | <0.01 | 0.22 ± 0.03 | n.d. |
| 3 | <0.01 | 0.06 ± 0.01 | n.d. |
| 4 | <0.01 | 0.11 ± 0.02 | n.d. |
| 5 | 1.83 ± 0.01 | 2.35 ± 0.03 | K363aR363bR363c |
| 6 | <0.01 | 0.16 ± 0.04 | n.d. |
| 7 | <0.01 | 0.20 ± 0.06 | n.d. |
| 8 | <0.01 | 0.14 ± 0.03 | G96aC96bR96c |
| 9 | <0.01 | 0.13 ± 0.03 | R138aE138bA138c |
| 10 | <0.01 | 0.02 ± 0.01 | D253A/R253a -Stop |
| 11 | <0.01 | 0.04 ± 0.02 | F342aR342bL342c |
| 12 | <0.01 | 0.12 ± 0.02 | n.d. |
| 13 | <0.01 | 0.07 ± 0.01 | n.d. |
| 14 | <0.01 | 0.06 ± 0.01 | n.d. |
| 15 | 0.30 ± 0.02 | 0.51 ± 0.03 | S211aV211bS211c |
| 16 | <0.01 | 0.19 ± 0.03 | n.d. |
| 17 | <0.01 | 0.03 ± 0.01 | n.d. |
| 18 | 0.17 ± 0.01 | 4.55 ± 0.24 | K257aH257bG257c |
| 19 | <0.01 | 0.09 ± 0.01 | n.d. |

(Table S9 continued)

| Variant | Relative activity | | Protein mutation |
|---|---|---|---|
| | Paraoxon | 4-NPB | |
| 20 | <0.01 | 0.09 ± 0.02 | n.d. |
| 21 | <0.01 | <0.01 | n.d. |
| 22 | <0.01 | 0.02 ± 0.01 | n.d. |
| 23 | <0.01 | 0.07 ± 0.01 | n.d. |
| 24 | <0.01 | 0.02 ± 0.01 | n.d. |
| 25 | <0.01 | <0.01 | frameshift |
| 26 | <0.01 | <0.01 | n.d. |
| 27 | <0.01 | <0.01 | n.d. |
| 28 | <0.01 | <0.01 | n.d. |
| 29 | <0.01 | 0.11 ± 0.01 | n.d. |
| 30 | <0.01 | 0.14 ± 0.03 | n.d. |
| 31 | <0.01 | <0.01 | n.d. |
| 32 | <0.01 | 0.21 ± 0.02 | n.d. |
| 33 | <0.01 | 0.04 ± 0.01 | n.d. |
| 34 | 0.16 ± 0.02 | 1.22 ± 0.12 | n.d. |
| 35 | 0.02 ± 0.01 | 0.02 ± 0.01 | n.d. |
| 36 | <0.01 | <0.01 | n.d. |
| 37 | <0.01 | <0.01 | n.d. |
| 38 | 0.12 ± 0.01 | 1.46 ± 0.08 | n.d. |
| 39 | <0.01 | 0.12 ± 0.01 | n.d. |
| 40 | <0.01 | 0.10 ± 0.02 | n.d. |
| 41 | <0.01 | <0.01 | n.d. |
| 42 | <0.01 | 0.03 ± 0.01 | n.d. |
| 43 | <0.01 | <0.01 | n.d. |
| 44 | 0.22 ± 0.01 | 5.4 ± 0.3 | C261aK261bL261c |
| 45 | 0.23 ± 0.01 | 0.32 ± 0.02 | n.d. |
| 46 | <0.01 | 0.02 ± 0.01 | n.d. |
| 47 | <0.01 | 0.13 ± 0.02 | n.d. |
| 48 | <0.01 | <0.01 | n.d. |
| 49 | <0.01 | <0.01 | n.d. |
| 50 | <0.01 | <0.01 | n.d. |
| 51 | <0.01 | <0.01 | n.d. |
| 52 | <0.01 | <0.01 | n.d. |
| 53 | <0.01 | <0.01 | n.d. |
| 54 | <0.01 | 0.07 ± 0.01 | n.d. |
| 55 | <0.01 | <0.01 | n.d. |
| 56 | <0.01 | <0.01 | n.d. |
| 57 | <0.01 | <0.01 | n.d. |
| 58 | <0.01 | <0.01 | n.d. |
| 59 | <0.01 | 0.05 ± 0.01 | n.d. |
| 60 | <0.01 | <0.01 | n.d. |
| 61 | <0.01 | 1.01 ± 0.06 | n.d. |
| 62 | <0.01 | <0.01 | n.d. |
| 63 | <0.01 | <0.01 | n.d. |

(Table S9 continued)

| Variant | Relative activity | | Protein mutation |
|---|---|---|---|
| | Paraoxon | 4-NPB | |
| 64 | 0.12 ± 0.01 | 0.20 ± 0.02 | n.d. |
| 65 | <0.01 | 0.08 ± 0.02 | n.d. |
| 66 | 1.91 ± 0.05 | 2.01 ± 0.01 | E36aY36bK36c |
| 67 | <0.01 | <0.01 | n.d. |
| 68 | <0.01 | <0.01 | n.d. |
| 69 | <0.01 | 0.15 ± 0.04 | n.d. |
| 70 | 0.28 ± 0.01 | 2.09 ± 0.08 | A309aA309bA309c |
| 71 | <0.01 | 0.20 ± 0.03 | n.d. |
| 72 | <0.01 | 0.02 ± 0.01 | n.d. |
| 73 | <0.01 | 0.21 ± 0.05 | n.d. |
| 74 | 0.04 ± 0.01 | <0.01 | n.d. |
| 75 | <0.01 | <0.01 | n.d. |
| 76 | <0.01 | <0.01 | n.d. |
| 77 | <0.01 | <0.01 | n.d. |
| 78 | <0.01 | <0.01 | n.d. |
| 79 | <0.01 | <0.01 | n.d. |
| 80 | <0.01 | <0.01 | n.d. |
| 81 | <0.01 | 0.04 ± 0.01 | n.d. |
| 82 | <0.01 | <0.01 | n.d. |
| 83 | <0.01 | <0.01 | n.d. |
| 84 | <0.01 | <0.01 | n.d. |
| 85 | 1.51 ± 0.06 | 1.06 ± 0.06 | L205aG205b |
| 86 | 2.78 ± 0.05 | 2.40 ± 0.09 | P262aT262bT262c |
| 87 | <0.01 | <0.01 | n.d. |
| 88 | <0.01 | 0.04 ± 0.01 | n.d. |
| 89 | 0.02 ± 0.01 | 0.13 ± 0.01 | n.d. |
| 90 | <0.01 | 0.17 ± 0.04 | n.d. |
| 91 | <0.01 | <0.01 | n.d. |
| 92 | <0.01 | 0.02 ± 0.01 | n.d. |
| 93 | 0.01 ± 0.01 | 0.04 ± 0.01 | n.d. |
| 94 | <0.01 | 0.04 ± 0.01 | n.d. |
| 95 | <0.01 | 0.04 ± 0.01 | n.d. |
| 96 | <0.01 | <0.01 | n.d. |
| 97 | <0.01 | <0.01 | G69aN69bP69c |
| 98 | <0.01 | 0.05 ± 0.02 | P197L/E197aW197bA197c |
| 99 | <0.01 | 0.14 ± 0.02 | M317I/F317aF317bL317c |
| 100 | <0.01 | <0.01 | E181D/Y181aP181bQ181c |
| 101 | <0.01 | <0.01 | H230R/R230aV230bG230c |
| 102 | 0.60 ± 0.01 | 0.54 ± 0.01 | n.d. |
| 103 | 0.05 ± 0.01 | 0.42 ± 0.01 | n.d. |
| 104 | <0.01 | <0.01 | n.d. |
| 105 | <0.01 | <0.01 | n.d. |
| 106 | <0.01 | 0.10 ± 0.03 | n.d. |
| 107 | <0.01 | 0.19 ± 0.01 | n.d. |

(Table S9 continued)

| Variant | Relative activity | | Protein mutation |
|---|---|---|---|
| | Paraoxon | 4-NPB | |
| 108 | 0.16 ± 0.02 | 0.19 ± 0.03 | n.d. |
| 109 | <0.01 | <0.01 | n.d. |
| 110 | 1.87 ± 0.09 | 1.69 ± 0.06 | S35aC35bP35c |
| 111 | <0.01 | 0.05 ± 0.01 | n.d. |
| 112 | <0.01 | 0.08 ± 0.01 | n.d. |
| 113 | <0.01 | <0.01 | n.d. |
| 114 | <0.01 | 0.05 ± 0.01 | n.d. |
| 115 | <0.01 | <0.01 | n.d. |
| 116 | <0.01 | 0.07 ± 0.01 | n.d. |
| 117 | <0.01 | 0.09 ± 0.02 | n.d. |
| 118 | <0.01 | 0.08 ± 0.03 | n.d. |
| 119 | <0.01 | <0.01 | n.d. |
| 120 | <0.01 | 0.03 ± 0.01 | n.d. |
| 121 | <0.01 | 0.03 ± 0.01 | n.d. |
| 122 | <0.01 | 0.09 ± 0.03 | n.d. |
| 123 | <0.01 | 0.09 ± 0.01 | n.d. |
| 124 | <0.01 | <0.01 | n.d. |
| 125 | 0.13 ± 0.01 | 1.26 ± 0.01 | n.d. |
| 126 | <0.01 | 0.25 ± 0.03 | n.d. |
| 127 | <0.01 | 0.04 ± 0.01 | n.d. |
| 128 | <0.01 | <0.01 | n.d. |
| 129 | <0.01 | <0.01 | n.d. |
| 130 | <0.01 | 0.04 ± 0.01 | n.d. |
| 131 | <0.01 | <0.01 | n.d. |
| 132 | <0.01 | 0.09 ± 0.02 | n.d. |
| 133 | <0.01 | <0.01 | n.d. |
| 134 | <0.01 | 0.03 ± 0.01 | n.d. |
| 135 | <0.01 | 0.03 ± 0.01 | n.d. |
| 136 | <0.01 | <0.01 | n.d. |
| 137 | <0.01 | 0.07 ± 0.01 | n.d. |
| 138 | 1.18 ± 0.1 | 3.29 ± 0.05 | D261aT261bS261c |
| 139 | <0.01 | 0.12 ± 0.02 | n.d. |
| 140 | <0.01 | <0.01 | n.d. |
| 141 | <0.01 | 0.03 ± 0.01 | n.d. |
| 142 | <0.01 | <0.01 | n.d. |
| 143 | <0.01 | 0.14 ± 0.02 | n.d. |
| 144 | <0.01 | 0.10 ± 0.02 | n.d. |
| 145 | <0.01 | <0.01 | n.d. |
| 146 | <0.01 | <0.01 | n.d. |
| 147 | <0.01 | <0.01 | n.d. |
| 148 | <0.01 | 0.20 ± 0.04 | n.d. |
| 149 | <0.01 | <0.01 | n.d. |
| 150 | 0.43 ± 0.01 | 2.85 ± 0.03 | D261aW261bK261c |
| 151 | <0.01 | 0.04 ± 0.01 | n.d. |

(Table S9 continued)

| Variant | Relative activity | | Protein mutation |
|---|---|---|---|
| | Paraoxon | 4-NPB | |
| 152 | <0.01 | 0.12 ± 0.01 | n.d. |
| 153 | <0.01 | 0.12 ± 0.01 | n.d. |
| 154 | <0.01 | <0.01 | n.d. |
| 155 | <0.01 | <0.01 | n.d. |
| 156 | <0.01 | 0.08 ± 0.01 | n.d. |
| 157 | <0.01 | 0.13 ± 0.02 | n.d. |
| 158 | <0.01 | 0.08 ± 0.01 | n.d. |
| 159 | <0.01 | 0.08 ± 0.01 | n.d. |
| 160 | <0.01 | 0.30 ± 0.06 | n.d. |
| 161 | <0.01 | 0.12 ± 0.02 | n.d. |
| 162 | <0.01 | 0.04 ± 0.01 | n.d. |
| 163 | <0.01 | <0.01 | n.d. |
| 164 | <0.01 | 0.18 ± 0.03 | n.d. |
| 165 | <0.01 | <0.01 | n.d. |
| 166 | <0.01 | 0.05 ± 0.01 | n.d. |
| 167 | 0.17 ± 0.02 | 0.08 ± 0.03 | n.d. |
| 168 | <0.01 | 0.20 ± 0.01 | n.d. |
| 169 | <0.01 | <0.01 | n.d. |
| 170 | <0.01 | <0.01 | n.d. |
| 171 | <0.01 | 0.12 ± 0.04 | n.d. |
| 172 | <0.01 | 0.22 ± 0.03 | n.d. |
| 173 | <0.01 | 0.11 ± 0.04 | n.d. |
| 174 | <0.01 | <0.01 | n.d. |
| 175 | <0.01 | <0.01 | n.d. |
| 176 | <0.01 | 0.30 ± 0.05 | n.d. |
| 177 | 2.70 ± 0.01 | 2.34 ± 0.05 | wt |
| 178 | <0.01 | 0.05 ± 0.02 | n.d. |
| **Triplet nucleotide exchange** | | | |
| 1 | 0.93 ± 0.16 | 1.00 ± 0.11 | D109C |
| 2 | 1.13 ± 0.13 | 1.19 ± 0.06 | S47Y |
| 3 | <0.01 | <0.01 | n.d. |
| 4 | 0.05 ± 0.01 | 0.04 ± 0.01 | I250M/G251L |
| 5 | 0.03 ± 0.01 | <0.01 | L262Q |
| 6 | <0.01 | <0.01 | frameshift |
| 7 | 0.62 ± 0.04 | 0.63 ± 0.05 | n.d. |
| 8 | 0.63 ± 0.02 | 0.75 ± 0.06 | n.d. |
| 9 | 0.07 ± 0.02 | 0.07 ± 0.02 | n.d. |
| 10 | 1.06 ± 0.18 | 1.24 ± 0.08 | Q343L |
| 11 | 0.12 ± 0.03 | 0.14 ± 0.03 | F72S/F73L |
| 12 | 0.13 ± 0.03 | 0.13 ± 0.03 | S75R/R76F |
| 13 | 0.02 ± 0.01 | <0.01 | G50P |
| 14 | <0.01 | <0.01 | frameshift |
| 15 | 0.05 ± 0.01 | 0.02 ± 0.01 | P256STOP |
| 16 | 0.10 ± 0.02 | 0.07 ± 0.01 | G229N |

(Table S9 continued)

| Variant | Relative activity | | Protein mutation |
|---|---|---|---|
| | Paraoxon | 4-NPB | |
| 17 | 0.58 ± 0.07 | 0.90 ± 0.03 | n.d. |
| 18 | <0.01 | <0.01 | n.d. |
| 19 | 0.95 ± 0.07 | 1.00 ± 0.10 | R363G |
| 20 | 0.54 ± 0.06 | 0.68 ± 0.09 | n.d. |
| 21 | 0.31 ± 0.03 | 0.33 ± 0.01 | G209C |
| 22 | <0.01 | <0.01 | P70R/E71STOP |
| 23 | 0.70 ± 0.08 | 0.62 ± 0.01 | n.d. |
| 24 | <0.01 | <0.01 | L336P |
| 25 | <0.01 | <0.01 | n.d. |
| 26 | 0.74 ± 0.09 | 0.84 ± 0.09 | n.d. |
| 27 | <0.01 | <0.01 | E153A/I154L |
| 28 | <0.01 | <0.01 | S307L/S308G |
| 29 | <0.01 | <0.01 | frameshift |
| 30 | 0.83 ± 0.08 | 1.37 ± 0.11 | Q155L |
| 31 | 0.12 ± 0.02 | 0.48 ± 0.08 | L330P |
| 32 | 0.82 ± 0.08 | 0.79 ± 0.08 | Y292G |
| 33 | 0.97 ± 0.09 | 1.27 ± 0.09 | E263Q |
| 34 | 0.69 ± 0.09 | 0.78 ± 0.13 | n.d. |
| 35 | 0.15 ± 0.09 | 0.15 ± 0.08 | T199V |
| 36 | 0.66 ± 0.10 | 0.75 ± 0.12 | n.d. |
| 37 | 0.16 ± 0.04 | 0.16 ± 0.04 | V183E |
| 38 | <0.01 | <0.01 | W69A |
| 39 | 1.16 ± 0.14 | 1.23 ± 0.08 | R67G |
| 40 | 0.89 ± 0.01 | 0.94 ± 0.03 | wtPTE |
| 41 | 0.79 ± 0.05 | 0.85 ± 0.05 | n.d. |
| 42 | 0.87 ± 0.06 | 0.87 ± 0.08 | wtPTE |
| 43 | <0.01 | <0.01 | n.d. |
| 44 | <0.01 | <0.01 | n.d. |
| 45 | 0.45 ± 0.02 | 0.59 ± 0.09 | n.d. |
| 46 | 0.08 ± 0.02 | 0.28 ± 0.05 | S276F/W277R |
| 47 | 0.07 ± 0.02 | 0.01 ± 0.01 | Y248T |
| 48 | 0.28 ± 0.02 | 0.72 ± 0.06 | n.d. |
| 49 | <0.01 | <0.01 | S62F/A63T |
| 50 | 0.81 ± 0.04 | 0.91 ± 0.08 | n.d. |
| 51 | <0.01 | <0.01 | n.d. |
| 52 | 1.05 ± 0.32 | 0.77 ± 0.05 | L282S |
| 53 | <0.01 | <0.01 | n.d. |
| 54 | 0.18 ± 0.03 | 0.10 ± 0.01 | G229A |
| 55 | 1.01 ± 0.12 | 1.08 ± 0.02 | Q343G |
| 56 | <0.01 | <0.01 | G60C |
| 57 | 0.20 ± 0.04 | 0.11 ± 0.02 | L151D |
| 58 | <0.01 | <0.01 | E48STOP |
| 59 | 0.68 ± 0.11 | 0.64 ± 0.07 | n.d. |
| 60 | 0.65 ± 0.05 | 0.60 ± 0.08 | n.d. |

(Table S9 continued)

| Variant | Relative activity | | Protein mutation |
|---|---|---|---|
| | Paraoxon | 4-NPB | |
| 61 | 0.69 ± 0.10 | 0.87 ± 0.06 | n.d. |
| 62 | 0.99 ± 0.21 | 0.94 ± 0.05 | A93V |
| 63 | <0.01 | <0.01 | V143A/E144STOP |
| 64 | 0.02 ± 0.01 | 0.30 ± 0.05 | n.d. |
| 65 | 0.89 ± 0.05 | 0.85 ± 0.02 | A92G/A93T |
| 66 | <0.01 | <0.01 | n.d. |
| 67 | <0.01 | <0.01 | D100Y |
| 68 | 0.77 ± 0.11 | 0.77 ± 0.02 | R96T |
| 69 | 0.93 ± 0.19 | 0.69 ± 0.04 | S102G |
| 70 | 0.19 ± 0.08 | 0.69 ± 0.13 | n.d. |
| 71 | 0.98 ± 0.14 | 0.89 ± 0.05 | R189 silent |
| 72 | 0.81 ± 0.12 | 0.79 ± 0.03 | n.d. |
| 73 | 0.35 ± 0.16 | 0.44 ± 0.12 | n.d. |
| 74 | 0.92 ± 0.04 | 0.81 ± 0.01 | n.d. |
| 75 | <0.01 | <0.01 | n.d. |
| 76 | 0.22 ± 0.09 | 0.12 ± 0.05 | n.d. |
| 77 | 0.41 ± 0.10 | 0.75 ± 0.08 | n.d. |
| 78 | 0.98 ± 0.20 | 1.02 ± 0.07 | V143A |
| 79 | <0.01 | <0.01 | n.d. |
| 80 | 0.93 ± 0.09 | 0.94 ± 0.04 | A266D |
| 81 | 0.60 ± 0.05 | 0.56 ± 0.02 | n.d. |
| 82 | 0.83 ± 0.01 | 0.79 ± 0.03 | R118K |
| 83 | 0.81 ± 0.13 | 0.74 ± 0.03 | n.d. |
| 84 | 0.31 ± 0.09 | 0.32 ± 0.1 | L336A |
| 85 | 0.06 ± 0.01 | 1.15 ± 0.1 | L271T |
| 86 | <0.01 | <0.01 | n.d. |
| 87 | 1.02 ± 0.4 | 0.99 ± 0.2 | n.d. |
| 88 | 1.06 ± 0.2 | 1.21 ± 0.07 | E144Q |
| 89 | 0.67 ± 0.08 | 0.63 ± 0.02 | n.d. |
| 90 | 0.88 ± 0.19 | 0.92 ± 0.08 | E144I |
| 91 | <0.01 | <0.01 | n.d. |
| 92 | 0.83 ± 0.06 | 0.76 ± 0.03 | n.d. |
| 93 | <0.01 | <0.01 | n.d. |
| 94 | 0.47 ± 0.05 | 0.55 ± 0.04 | n.d. |
| 95 | 0.55 ± 0.02 | 0.49 ± 0.01 | n.d. |
| 96 | <0.01 | <0.01 | n.d. |
| 97 | <0.01 | <0.01 | n.d. |
| 98 | 1.04 ± 0.21 | 1.00 ± 0.03 | A80G |
| 99 | 0.03 ± 0.01 | 1.11 ± 0.09 | n.d. |
| 100 | 0.61 ± 0.06 | 0.56 ± 0.01 | n.d. |
| 101 | 0.06 ± 0.01 | 0.05 ± 0.01 | n.d. |
| 102 | <0.01 | <0.01 | n.d. |
| 103 | <0.01 | <0.01 | n.d. |
| 104 | <0.01 | <0.01 | n.d. |

(Table S9 continued)

| Variant | Relative activity | | Protein mutation |
|---|---|---|---|
| | Paraoxon | 4-NPB | |
| 105 | <0.01 | 0.04 ± 0.02 | n.d. |
| 106 | 0.66 ± 0.14 | 0.46 ± 0.02 | n.d. |
| 107 | 1.04 ± 0.22 | 0.83 ± 0.02 | F73C |
| 108 | 0.04 ± 0.04 | 0.02 ± 0.01 | n.d. |
| 109 | <0.01 | <0.01 | n.d. |
| 110 | <0.01 | <0.01 | n.d. |
| 111 | 0.02 ± 0.01 | <0.01 | n.d. |
| 112 | <0.01 | <0.01 | n.d. |
| 113 | 0.39 ± 0.08 | 0.31 ± 0.02 | n.d. |
| 114 | <0.01 | <0.01 | n.d. |
| 115 | 0.13 ± 0.11 | 0.10 ± 0.06 | n.d. |
| 116 | <0.01 | <0.01 | n.d. |
| 117 | 1.11 ± 0.12 | 1.09 ± 0.04 | R118G |
| 118 | 0.41 ± 0.02 | 0.65 ± 0.13 | n.d. |
| 119 | 0.50 ± 0.01 | 0.42 ± 0.02 | n.d. |
| 120 | 0.09 ± 0.03 | 1.97 ± 0.12 | L272R/G273R |
| 121 | 0.60 ± 0.04 | 0.55 ± 0.05 | n.d. |
| 122 | 0.88 ± 0.06 | 1.07 ± 0.01 | R331G |
| 123 | <0.01 | <0.01 | n.d. |
| 124 | 0.12 ± 0.03 | 0.10 ± 0.02 | n.d. |
| 125 | 1.02 ± 0.07 | 0.95 ± 0.05 | N312D |
| 126 | <0.01 | <0.01 | n.d. |
| 127 | 0.80 ± 0.10 | 0.72 ± 0.02 | n.d. |
| 128 | <0.01 | <0.01 | n.d. |
| 129 | <0.01 | <0.01 | n.d. |
| 130 | 0.12 ± 0.01 | 0.07 ± 0.01 | n.d. |
| 131 | <0.01 | <0.01 | n.d. |
| 132 | 0.71 ± 0.06 | 0.65 ± 0.04 | n.d. |
| 133 | 1.05 ± 0.13 | 0.97 ± 0.05 | L330S |
| 134 | <0.01 | <0.01 | n.d. |
| 135 | 0.24 ± 0.11 | 0.20 ± 0.05 | n.d. |
| 136 | 1.04 ± 0.10 | 1.03 ± 0.01 | M293T |
| 137 | 0.98 ± 0.09 | 1.04 ± 0.01 | R118G |
| 138 | 0.85 ± 0.10 | 1.08 ± 0.02 | S75G |
| 139 | 0.85 ± 0.02 | 0.81 ± 0.04 | T147S |
| 140 | 0.19 ± 0.15 | 0.14 ± 0.08 | n.d. |
| 141 | <0.01 | <0.01 | n.d. |
| 142 | 1.08 ± 0.18 | 0.99 ± 0.07 | L330I |
| 143 | <0.01 | <0.01 | n.d. |
| 144 | <0.01 | <0.01 | n.d. |
| 145 | <0.01 | <0.01 | n.d. |
| 146 | 0.76 ± 0.03 | 0.70 ± 0.04 | n.d. |
| 147 | 1.05 ± 0.02 | 0.98 ± 0.04 | P342L/Q343E |
| 148 | 0.26 ± 0.01 | 0.25 ± 0.01 | n.d. |

(Table S9 continued)

| Variant | Relative activity | | Protein mutation |
|---|---|---|---|
| | Paraoxon | 4-NPB | |
| 149 | 0.59 ± 0.04 | 0.53 ± 0.04 | n.d. |
| 150 | 0.77 ± 0.11 | 0.82 ± 0.07 | n.d. |
| 151 | 0.88 ± 0.10 | 0.85 ± 0.04 | n.d. |
| 152 | <0.01 | <0.01 | n.d. |
| 153 | 0.85 ± 0.03 | 0.80 ± 0.03 | L262H |
| 154 | 1.07 ± 0.08 | 0.98 ± 0.03 | Q343A |
| 155 | 0.07 ± 0.01 | 1.80 ± 0.07 | L272H/G273R |
| 156 | 0.64 ± 0.06 | 0.60 ± 0.02 | n.d. |
| 157 | 0.85 ± 0.08 | 0.73 ± 0.05 | I44K/T45A |
| 158 | 0.41 ± 0.02 | 0.34 ± 0.02 | n.d. |
| 159 | <0.01 | <0.01 | n.d. |
| 160 | 1.20 ± 0.10 | 1.11 ± 0.07 | S47Y |
| 161 | 0.21 ± 0.10 | 0.15 ± 0.04 | n.d. |
| 162 | 0.08 ± 0.01 | <0.01 | n.d. |
| 163 | 0.69 ± 0.11 | 0.68 ± 0.06 | n.d. |
| 164 | <0.01 | 0.07 ± 0.01 | I313K |
| 165 | <0.01 | <0.01 | n.d. |
| 166 | 1.13 ± 0.22 | 0.99 ± 0.08 | S117 silent |
| 167 | 0.07 ± 0.04 | 0.02 ± 0.01 | n.d. |
| 168 | 0.03 ± 0.01 | 0.03 ± 0.01 | M314K/D315Y |
| 169 | 0.03 ± 0.01 | <0.01 | n.d. |
| 170 | 0.96 ± 0.07 | 1.03 ± 0.08 | n.d. |
| 171 | 1.01 ± 0.07 | 0.94 ± 0.07 | n.d. |
| 172 | 1.11 ± 0.10 | 0.91 ± 0.06 | Y292R |
| 173 | 1.02 ± 0.17 | 0.53 ± 0.05 | D323E/G324R |
| 174 | 0.97 ± 0.19 | 1.14 ± 0.13 | T54S |
| 175 | 1.17 ± 0.12 | 1.18 ± 0.04 | Y292F |
| 176 | 0.72 ± 0.14 | 0.73 ± 0.05 | n.d. |
| 177 | 0.86 ± 0.08 | 0.91 ± 0.06 | L330A |
| 178 | <0.01 | 0.02 ± 0.01 | n.d. |
| 179 | <0.01 | 0.02 ± 0.01 | n.d. |
| 180 | <0.01 | 0.02 ± 0.01 | n.d. |
| 181 | 0.69 ± 0.04 | 0.94 ± 0.07 | n.d. |
| 182 | 0.78 ± 0.04 | 0.88 ± 0.05 | L262M |
| 183 | 0.91 ± 0.08 | 0.92 ± 0.06 | G348 silent |
| 184 | 0.07 ± 0.03 | 0.08 ± 0.03 | n.d. |
| 185 | 0.79 ± 0.06 | 0.81 ± 0.05 | A242P |
| 186 | 0.71 ± 0.10 | 0.87 ± 0.07 | n.d. |
| 187 | 0.64 ± 0.06 | 0.95 ± 0.06 | n.d. |
| 188 | 0.72 ± 0.02 | 0.77 ± 0.05 | n.d. |
| 189 | 1.00 ± 0.11 | 0.85 ± 0.06 | D109S |
| 190 | <0.01 | 0.02 ± 0.01 | n.d. |
| 191 | 0.95 ± 0.05 | 1.05 ± 0.03 | R89F |
| 192 | 0.06 ± 0.01 | 0.05 ± 0.02 | n.d. |

(Table S9 continued)

| Variant | Relative activity | | Protein mutation |
|---|---|---|---|
| | Paraoxon | 4-NPB | |
| 193 | 0.01 ± 0.01 | 0.03 ± 0.02 | n.d. |
| 194 | 0.75 ± 0.09 | 0.76 ± 0.05 | n.d. |
| 195 | 0.88 ± 0.05 | 0.84 ± 0.06 | P342L |
| 196 | 0.06 ± 0.01 | 1.35 ± 0.11 | L272Q/G273S |
| 197 | <0.01 | 0.02 ± 0.01 | n.d. |
| 198 | <0.01 | 0.02 ± 0.01 | n.d. |
| 199 | 0.73 ± 0.03 | 0.75 ± 0.02 | n.d. |
| 200 | 0.67 ± 0.09 | 0.70 ± 0.07 | n.d. |
| 201 | 0.29 ± 0.05 | 0.29 ± 0.04 | K339I/G340R |
| 202 | 0.02 ± 0.01 | 0.04 ± 0.01 | n.d. |
| 203 | 0.63 ± 0.04 | 0.68 ± 0.02 | n.d. |
| 204 | <0.01 | 0.02 ± 0.01 | n.d. |
| 205 | <0.01 | 0.02 ± 0.01 | n.d. |
| 206 | <0.01 | 0.15 ± 0.07 | n.d. |
| 207 | <0.01 | 0.02 ± 0.01 | n.d. |
| 208 | 0.09 ± 0.01 | 0.08 ± 0.02 | n.d. |
| 209 | <0.01 | 0.03 ± 0.02 | n.d. |
| 210 | <0.01 | 0.02 ± 0.01 | n.d. |
| 211 | 0.52 ± 0.10 | 0.61 ± 0.04 | n.d. |
| 212 | 0.36 ± 0.04 | 0.45 ± 0.04 | n.d. |
| 213 | 0.91 ± 0.13 | 1.07 ± 0.07 | L151R |
| 214 | <0.01 | 0.02 ± 0.01 | n.d. |
| 215 | <0.01 | 0.02 ± 0.01 | n.d. |
| 216 | 0.22 ± 0.01 | 0.22 ± 0.02 | n.d. |
| 217 | 0.60 ± 0.07 | 0.66 ± 0.03 | n.d. |
| 218 | 1.08 ± 0.12 | 0.82 ± 0.02 | A80G/E81K |
| 219 | 0.96 ± 0.06 | 0.80 ± 0.02 | wtPTE |
| 220 | 0.03 ± 0.01 | 0.30 ± 0.02 | n.d. |
| 221 | 0.33 ± 0.01 | 0.26 ± 0.02 | n.d. |
| 222 | 0.21 ± 0.04 | 0.14 ± 0.02 | n.d. |
| 223 | 0.75 ± 0.08 | 0.71 ± 0.02 | n.d. |
| 224 | 0.26 ± 0.08 | 0.47 ± 0.08 | n.d. |
| 225 | 1.01 ± 0.15 | 0.91 ± 0.04 | wtPTE |
| 226 | 0.30 ± 0.03 | 0.22 ± 0.02 | n.d. |
| 227 | 1.07 ± 0.12 | 0.84 ± 0.02 | n.d. |
| 228 | <0.01 | 0.02 ± 0.01 | n.d. |
| 229 | 0.94 ± 0.04 | 0.88 ± 0.04 | P334N |
| 230 | 1.06 ± 0.09 | 0.98 ± 0.04 | P70 silent |
| 231 | <0.01 | 0.02 ± 0.01 | n.d. |
| 232 | 0.03 ± 0.01 | 0.11 ± 0.01 | M317S |
| 233 | 0.73 ± 0.12 | 0.61 ± 0.04 | n.d. |
| 234 | <0.01 | 0.02 ± 0.01 | n.d. |
| 235 | 0.41 ± 0.09 | 0.32 ± 0.05 | n.d. |
| 236 | 0.35 ± 0.11 | 0.36 ± 0.08 | n.d. |

(Table S9 continued)

| Variant | Relative activity | | Protein mutation |
|---|---|---|---|
| | Paraoxon | 4-NPB | |
| 237 | 0.25 ± 0.04 | 0.16 ± 0.01 | n.d. |
| 238 | <0.01 | 0.03 ± 0.02 | n.d. |
| 239 | <0.01 | 0.12 ± 0.01 | n.d. |
| 240 | 0.85 ± 0.04 | 0.77 ± 0.02 | G162R |
| 241 | 1.14 ± 0.26 | 0.92 ± 0.03 | R356E |
| 242 | 0.26 ± 0.03 | 0.34 ± 0.03 | n.d. |
| 243 | 0.82 ± 0.23 | 0.74 ± 0.11 | K339T |
| 244 | 1.20 ± 0.24 | 0.92 ± 0.08 | R118Q |
| 245 | <0.01 | 0.02 ± 0.01 | n.d. |
| 246 | 0.02 ± 0.01 | 0.22 ± 0.03 | n.d. |
| 247 | 0.63 ± 0.08 | 0.48 ± 0.03 | n.d. |
| 248 | 1.10 ± 0.09 | 0.95 ± 0.02 | R36N |
| 249 | <0.01 | 0.02 ± 0.01 | n.d. |
| 250 | 0.99 ± 0.21 | 0.84 ± 0.05 | G162V |
| 251 | <0.01 | 0.02 ± 0.01 | n.d. |
| 252 | 1.01 ± 0.22 | 0.80 ± 0.07 | A63T |
| 253 | 0.74 ± 0.17 | 0.59 ± 0.04 | n.d. |
| 254 | 0.04 ± 0.01 | 0.18 ± 0.02 | n.d. |
| 255 | 1.18 ± 0.05 | 0.93 ± 0.03 | S47C/E48Q |
| 256 | 0.40 ± 0.03 | 0.22 ± 0.01 | n.d. |
| 257 | 0.04 ± 0.01 | 0.49 ± 0.02 | n.d. |
| 258 | 0.72 ± 0.10 | 0.60 ± 0.01 | n.d. |
| 259 | 0.03 ± 0.01 | 0.02 ± 0.01 | n.d. |
| 260 | 1.14 ± 0.23 | 0.97 ± 0.06 | R337Y |
| 261 | 1.35 ± 0.46 | 0.92 ± 0.10 | n.d. |
| 262 | 0.24 ± 0.14 | 0.62 ± 0.29 | n.d. |
| 263 | 0.98 ± 0.04 | 1.08 ± 0.04 | L262 silent |
| 264 | 0.51 ± 0.09 | 0.58 ± 0.01 | T311A |
| 265 | <0.01 | <0.01 | n.d. |
| 266 | 0.99 ± 0.01 | 1.17 ± 0.03 | Q206L |
| 267 | 0.55 ± 0.10 | 0.74 ± 0.07 | n.d. |
| 268 | <0.01 | 0.31 ± 0.03 | Y309R |
| 269 | 0.63 ± 0.09 | 0.74 ± 0.03 | n.d. |
| 270 | 1.04 ± 0.18 | 1.16 ± 0.04 | wtPTE |
| 271 | 1.13 ± 0.19 | 0.99 ± 0.05 | V351D |
| 272 | 0.95 ± 0.08 | 1.08 ± 0.03 | L182 silent |
| 273 | <0.01 | <0.01 | n.d. |
| 274 | 0.03 ± 0.01 | <0.01 | n.d. |
| 275 | <0.01 | <0.01 | n.d. |
| 276 | 1.07 ± 0.05 | 1.25 ± 0.08 | S238G |
| 277 | 0.88 ± 0.09 | 1.00 ± 0.02 | A63S |
| 278 | 0.93 ± 0.13 | 0.88 ± 0.06 | L262I |
| 279 | <0.01 | <0.01 | n.d. |
| 280 | 0.86 ± 0.11 | 0.93 ± 0.06 | V198L |

(Table S9 continued)

| Variant | Relative activity | | Protein mutation |
|---|---|---|---|
| | Paraoxon | 4-NPB | |
| 281 | 0.72 ± 0.08 | 0.86 ± 0.04 | n.d. |
| 282 | 0.39 ± 0.13 | 0.51 ± 0.10 | n.d. |
| 283 | <0.01 | <0.01 | n.d. |
| 284 | <0.01 | <0.01 | n.d. |
| 285 | 0.95 ± 0.11 | 1.07 ± 0.03 | L262 silent |
| 286 | <0.01 | <0.01 | n.d. |
| 287 | 0.04 ± 0.01 | 0.19 ± 0.02 | n.d. |
| 288 | <0.01 | <0.01 | n.d. |
| 289 | 0.51 ± 0.11 | 0.84 ± 0.07 | n.d. |
| 290 | 0.38 ± 0.02 | 0.53 ± 0.03 | n.d. |
| 291 | 0.09 ± 0.02 | 0.06 ± 0.01 | n.d. |
| 292 | 1.11 ± 0.15 | 0.75 ± 0.04 | S102T |
| 293 | <0.01 | <0.01 | n.d. |
| 294 | 0.04 ± 0.01 | <0.01 | n.d. |
| 295 | 0.37 ± 0.16 | 0.73 ± 0.05 | n.d. |
| 296 | <0.01 | <0.01 | n.d. |
| 297 | 0.48 ± 0.13 | 0.66 ± 0.05 | n.d. |
| 298 | 0.55 ± 0.22 | 0.97 ± 0.07 | n.d. |
| 299 | 0.47 ± 0.06 | 0.76 ± 0.04 | L262E |
| 300 | 0.87 ± 0.28 | 1.15 ± 0.04 | R67T |
| 301 | <0.01 | <0.01 | n.d. |
| 302 | <0.01 | <0.01 | n.d. |
| 303 | 0.17 ± 0.06 | 0.23 ± 0.03 | n.d. |
| 304 | <0.01 | <0.01 | n.d. |
| 305 | 0.08 ± 0.01 | 0.79 ± 0.07 | R280S |
| 306 | 0.19 ± 0.04 | 0.08 ± 0.01 | n.d. |
| 307 | 0.70 ± 0.11 | 0.79 ± 0.01 | n.d. |
| 308 | <0.01 | <0.01 | n.d. |
| 309 | 0.73 ± 0.09 | 0.60 ± 0.05 | n.d. |
| 310 | <0.01 | 0.09 ± 0.01 | n.d. |
| 311 | 0.84 ± 0.19 | 0.88 ± 0.06 | Q206H |
| 312 | 0.54 ± 0.11 | 0.58 ± 0.06 | n.d. |
| 313 | 0.91 ± 0.14 | 0.92 ± 0.06 | P134 silent |
| 314 | <0.01 | <0.01 | n.d. |
| 315 | 0.02 ± 0.01 | <0.01 | n.d. |
| 316 | <0.01 | <0.01 | n.d. |
| 317 | <0.01 | <0.01 | n.d. |
| 318 | 0.80 ± 0.11 | 0.86 ± 0.02 | S365R |
| 319 | 0.81 ± 0.18 | 0.80 ± 0.04 | D100A |
| 320 | 0.87 ± 0.19 | 1.26 ± 0.05 | G208A |
| 321 | <0.01 | <0.01 | n.d. |
| 322 | 0.91 ± 0.17 | 0.87 ± 0.04 | Y292R |
| 323 | <0.01 | <0.01 | n.d. |
| 324 | 0.14 ± 0.03 | 0.10 ± 0.02 | n.d. |

(Table S9 continued)

| Variant | Relative activity | | Protein mutation |
|---|---|---|---|
| | Paraoxon | 4-NPB | |
| 325 | 0.34 ± 0.13 | 0.27 ± 0.06 | n.d. |
| 326 | <0.01 | <0.01 | n.d. |
| 327 | 0.91 ± 0.08 | 0.92 ± 0.03 | Q343I |
| 328 | 0.78 ± 0.09 | 0.78 ± 0.05 | I44T/T45A |
| 329 | 0.58 ± 0.13 | 0.63 ± 0.03 | n.d. |
| 330 | <0.01 | <0.01 | n.d. |
| 331 | 0.35 ± 0.06 | 0.28 ± 0.04 | n.d. |
| 332 | 0.15 ± 0.07 | 0.12 ± 0.03 | n.d. |
| 333 | <0.01 | <0.01 | n.d. |
| 334 | 0.64 ± 0.12 | 0.68 ± 0.05 | n.d. |
| 335 | 0.89 ± 0.32 | 0.74 ± 0.01 | P342W |
| 336 | 0.83 ± 0.15 | 0.87 ± 0.02 | K339R |
| 337 | 0.11 ± 0.02 | 0.05 ± 0.01 | n.d. |
| 338 | 0.47 ± 0.09 | 1.06 ± 0.06 | n.d. |
| 339 | 0.05 ± 0.01 | <0.01 | n.d. |
| 340 | 0.71 ± 0.04 | 0.76 ± 0.03 | n.d. |
| 341 | 0.29 ± 0.03 | 0.20 ± 0.01 | n.d. |
| 342 | 0.19 ± 0.06 | 0.35 ± 0.02 | n.d. |
| 343 | 0.95 ± 0.20 | 1.01 ± 0.03 | Y292F |
| 344 | 0.17 ± 0.05 | 0.22 ± 0.03 | n.d. |
| 345 | 0.93 ± 0.08 | 0.92 ± 0.01 | L221 silent |
| 346 | 0.43 ± 0.13 | 0.48 ± 0.08 | K339T/G340C |
| 347 | <0.01 | <0.01 | n.d. |
| 348 | <0.01 | <0.01 | n.d. |
| 349 | <0.01 | 0.35 ± 0.05 | n.d. |
| 350 | 0.11 ± 0.01 | 1.94 ± 0.11 | R280T |
| 351 | 1.21 ± 0.04 | 1.61 ± 0.07 | S269H |
| 352 | <0.01 | 0.03 ± 0.01 | ΔM317 |
| 353 | <0.01 | <0.01 | n.d. |
| 354 | <0.01 | 0.02 ± 0.01 | n.d. |
| 355 | 0.14 ± 0.03 | 0.13 ± 0.01 | n.d. |
| 356 | 0.79 ± 0.06 | 0.78 ± 0.02 | Q343W |
| 357 | <0.01 | <0.01 | n.d. |
| 358 | 1.11 ± 0.17 | 1.24 ± 0.08 | E263Q |
| 359 | 1.21 ± 0.23 | 1.10 ± 0.05 | Y292L |
| 360 | 1.03 ± 0.18 | 1.01 ± 0.10 | V143 silent |
| 361 | <0.01 | <0.01 | n.d. |
| 362 | <0.01 | <0.01 | n.d. |
| 363 | 1.13 ± 0.16 | 1.10 ± 0.03 | Q343L |
| 364 | 1.01 ± 0.04 | 0.99 ± 0.03 | P342R |
| 365 | 0.95 ± 0.12 | 0.92 ± 0.03 | A364 silent |
| 366 | 0.38 ± 0.03 | 0.27 ± 0.02 | n.d. |
| 367 | 0.07 ± 0.07 | 0.05 ± 0.03 | n.d. |
| 368 | 1.12 ± 0.29 | 0.84 ± 0.03 | T241I |

(Table S9 continued)

| Variant | Relative activity | | Protein mutation |
|---|---|---|---|
| | Paraoxon | 4-NPB | |
| 369 | 0.23 ± 0.03 | 0.32 ± 0.04 | n.d. |
| 370 | 0.44 ± 0.07 | 0.75 ± 0.08 | n.d. |
| 371 | 0.23 ± 0.01 | 0.27 ± 0.01 | V310A/ΔT311 |
| 372 | 0.72 ± 0.06 | 0.85 ± 0.05 | n.d. |
| 373 | 0.50 ± 0.09 | 0.45 ± 0.04 | n.d. |
| 374 | 0.83 ± 0.09 | 0.75 ± 0.03 | E219 silent |
| 375 | 0.18 ± 0.04 | 0.10 ± 0.01 | n.d. |
| 376 | <0.01 | 2.98 ± 0.26 | H254P |
| 377 | <0.01 | <0.01 | n.d. |
| 378 | <0.01 | <0.01 | n.d. |
| 379 | <0.01 | <0.01 | n.d. |
| 380 | 0.74 ± 0.06 | 0.85 ± 0.03 | n.d. |
| 381 | 0.86 ± 0.02 | 0.93 ± 0.02 | P334T |
| 382 | 0.02 ± 0.02 | 0.09 ± 0.05 | n.d. |
| 383 | 0.27 ± 0.08 | 0.29 ± 0.07 | n.d. |
| 384 | <0.01 | <0.01 | n.d. |
| 385 | 0.18 ± 0.01 | 1.45 ± 0.05 | L272C |
| 386 | 0.80 ± 0.10 | 1.68 ± 0.09 | W302F |
| 387 | 0.04 ± 0.01 | 0.03 ± 0.01 | n.d. |
| 388 | 1.07 ± 0.10 | 1.04 ± 0.02 | Q343P |
| 389 | <0.01 | <0.01 | n.d. |
| 390 | <0.01 | <0.01 | n.d. |
| 391 | 0.06 ± 0.01 | 1.20 ± 0.09 | L271A |
| 392 | <0.01 | 0.02 ± 0.01 | n.d. |
| 393 | 1.12 ± 0.20 | 0.91 ± 0.07 | P223A |
| 394 | 1.22 ± 0.19 | 0.99 ± 0.04 | n.d. |
| 395 | 0.62 ± 0.09 | 0.59 ± 0.08 | n.d. |
| 396 | <0.01 | <0.01 | n.d. |
| 397 | 1.03 ± 0.07 | 0.91 ± 0.03 | P329 silent |
| 398 | 0.41 ± 0.03 | 0.36 ± 0.03 | n.d. |
| 399 | <0.01 | <0.01 | n.d. |
| 400 | 0.61 ± 0.12 | 0.52 ± 0.07 | n.d. |
| 401 | 0.93 ± 0.05 | 0.96 ± 0.06 | P334A |
| 402 | <0.01 | <0.01 | n.d. |
| 403 | 1.14 ± 0.32 | 0.83 ± 0.04 | n.d. |
| 404 | <0.01 | <0.01 | n.d. |
| 405 | <0.01 | <0.01 | n.d. |
| 406 | 0.18 ± 0.05 | 0.13 ± 0.03 | n.d. |
| 407 | 0.43 ± 0.12 | 0.55 ± 0.10 | n.d. |
| 408 | 0.07 ± 0.01 | 1.44 ± 0.13 | L272P |
| 409 | <0.01 | 0.04 ± 0.01 | n.d. |
| 410 | 1.50 ± 0.31 | 1.03 ± 0.02 | Q343E |
| 411 | 0.02 ± 0.01 | 0.05 ± 0.01 | n.d. |
| 412 | 0.92 ± 0.06 | 1.31 ± 0.02 | D235T |

(Table S9 continued)

| Variant | Relative activity | | Protein mutation |
| --- | --- | --- | --- |
| | Paraoxon | 4-NPB | |
| 413 | <0.01 | <0.01 | n.d. |
| 414 | 0.76 ± 0.04 | 0.67 ± 0.06 | n.d. |
| 415 | 1.26 ± 0.36 | 1.34 ± 0.23 | S308C |
| 416 | 0.98 ± 0.19 | 0.72 ± 0.09 | S269A |
| 417 | <0.01 | <0.01 | n.d. |
| 418 | <0.01 | <0.01 | n.d. |
| 419 | <0.01 | <0.01 | n.d. |
| 420 | 1.03 ± 0.24 | 1.07 ± 0.04 | A266G |
| 421 | 0.01 ± 0.01 | <0.01 | n.d. |
| 422 | <0.01 | <0.01 | n.d. |
| 423 | 1.14 ± 0.21 | 0.88 ± 0.04 | A49M |
| 424 | 0.73 ± 0.18 | 0.52 ± 0.09 | n.d. |
| 425 | 1.10 ± 0.21 | 0.92 ± 0.07 | G348A |
| 426 | <0.01 | <0.01 | n.d. |
| 427 | 1.26 ± 0.22 | 1.00 ± 0.06 | T117N/P118A |
| 428 | <0.01 | <0.01 | n.d. |
| 429 | 1.35 ± 0.18 | 1.03 ± 0.06 | n.d. |
| 430 | 0.36 ± 0.07 | 0.23 ± 0.04 | n.d. |
| 431 | <0.01 | <0.01 | n.d. |
| 432 | 0.82 ± 0.19 | 0.68 ± 0.11 | V143A/E144K |
| 433 | 1.25 ± 0.08 | 1.06 ± 0.08 | R67H |
| 434 | 0.01 ± 0.01 | <0.01 | n.d. |
| 435 | 1.49 ± 0.23 | 1.11 ± 0.07 | n.d. |

**Supplementary Table S10. Analysis of solvent-accessible surface area of mutated residues in *wt*PTE variants retaining ≥50% of the parental paraoxonase activity.**

The solvent accessible surface area (SASA) of residues mutated (either InDel or substitution) in *wt*PTE variants retaining ≥50% of the parental paraoxonase activity was calculated from the structure of *wt*PTE (PDB code: 4PCP) using the PISA web server at the European Bioinformatics Institute (http://www.ebi.ac.uk/pdbe/prot_int/pistart.html) [14]. Relative accessible surface area (RSA) was defined as the ratio of the SASA for a given residue within the structured protein *vs.* in the free residue [15]. Residues were classified as core for RSA < 0.25, and surface for RSA ≥ 0.25 [16].

**Supplementary Table S10a. List of residues in *wt*PTE InDel variants retaining ≥50% of the parental paraoxonase activity.**

| Residue no. | SASA (Å$^2$) | RSA | Location | Corresponding variants |
|---|---|---|---|---|
| 34 | 79.4 | 0.93 | surface | F34aN34b |
| 35 | 88.8 | 0.59 | surface | ∆D35R36, P35a, F35a, K35aC35b, F35aH35b, S35aC35bP35c |
| 36 | 124.2 | 0.52 | surface | ∆R36, E36aY36bK36c |
| 37 | 4.5 | 0.02 | core | T37aI37b |
| 42 | 24.2 | 0.28 | surface | H42a |
| 43 | 77.6 | 0.54 | surface | P43H/A43a |
| 45 | 48.6 | 0.33 | surface | C45aP45b |
| 49 | 1.6 | 0.01 | core | ∆A49 |
| 75 | 42.7 | 0.35 | surface | ∆S75R76K77 |
| 155 | 96.2 | 0.51 | surface | ∆Q155 |
| 161 | 48.7 | 0.33 | surface | T161K/P161a |
| 173 | 66.1 | 0.45 | surface | ∆T173 |
| 174 | 8.0 | 0.09 | core | ∆G174 |
| 203 | 34.7 | 0.31 | surface | ∆A203 |
| 205 | 85.0 | 0.70 | surface | L205aG205b |
| 206 | 102.0 | 0.54 | surface | Q206H/∆R207G208 |
| 259 | 10.4 | 0.09 | core | P259a |
| 261 | 56.0 | 0.66 | surface | S261a, P261a, G261aR261b, D261aT261bS261c |
| 262 | 41.6 | 0.23 | core | ∆L262, I262a, P262aT262bT262c |
| 263 | 126.4 | 0.69 | surface | ∆E263, E263V/Q263a |
| 266 | 97.8 | 0.87 | surface | ∆A266, G266aR266b |
| 267 | 31.8 | 0.26 | surface | A266G/∆S267 |
| 269 | 50.3 | 0.41 | surface | ∆S269 |
| 292 | 56.5 | 0.25 | core | ∆Y292 |
| 293 | 45.9 | 0.22 | core | ∆M293, M293I/∆K294 |
| 319 | 84.7 | 0.69 | surface | S319a |

(Table S10 continued)

| Residue no. | SASA (Å$^2$) | RSA | Location | Corresponding variants |
|---|---|---|---|---|
| 337 | 109.0 | 0.45 | surface | ΔR337 |
| 338 | 126.9 | 0.69 | surface | ΔE338 |
| 339 | 91.4 | 0.43 | surface | Q339a, K339M/Q339a, V339a |
| 362 | 122.6 | 0.68 | surface | ΔL362, K362a |
| 363 | 178.5 | 0.74 | surface | K363aR363bR363c |

**Supplementary Table S10b. List of residues in wtPTE substitution variants retaining ≥50% of the parental paraoxonase activity.**

| Residue no. | SASA (Å$^2$) | RSA | Location | Corresponding variants |
|---|---|---|---|---|
| 36 | 124.2 | 0.52 | surface | R36N |
| 44 | 18.0 | 0.10 | core | I44K/T45A, I44T/T45A |
| 45 | 48.6 | 0.33 | surface | I44K/T45A, I44T/T45A |
| 47 | 98.0 | 0.80 | surface | S47Y, S47C/E48Q |
| 48 | 96.6 | 0.53 | surface | S47C/E48Q |
| 49 | 1.6 | 0.01 | core | A49M |
| 54 | 9.0 | 0.06 | core | T54S |
| 63 | 41.2 | 0.36 | surface | A63T, A63S |
| 67 | 183.9 | 0.76 | surface | R67G, R67T, R67H |
| 73 | 29.3 | 0.13 | core | F73C |
| 75 | 42.7 | 0.35 | surface | S75G |
| 80 | 8.1 | 0.07 | core | A80G, A80G/E81K |
| 81 | 75.9 | 0.41 | surface | A80G/E81K |
| 89 | 124.0 | 0.51 | surface | R89F |
| 92 | 94.9 | 0.84 | surface | A92G/A93T |
| 93 | 27.5 | 0.24 | core | A93V |
| 96 | 85.8 | 0.36 | surface | R96T |
| 100 | 0.9 | 0.01 | core | D100A |
| 102 | 1.3 | 0.01 | core | S102G, S102T |
| 109 | 20.8 | 0.14 | core | D109C, D109S |
| 117 | 0.1 | 0.00 | core | T117N/P118A |
| 118 | 148.9 | 0.62 | surface | R118K, R118G, R118Q, T117N/P118A |
| 143 | 19.7 | 0.12 | core | V143A, V143A/E144K |
| 144 | 109.2 | 0.60 | surface | E144Q, E144I |
| 147 | 13.6 | 0.09 | core | T147S |
| 151 | 22.0 | 0.12 | core | L151R |
| 155 | 96.2 | 0.51 | surface | Q155L |
| 162 | 53.2 | 0.63 | surface | G162R, G162V |

(Table S10 continued)

| Residue no. | SASA (Å$^2$) | RSA | Location | Corresponding variants |
|---|---|---|---|---|
| 198 | 0.0 | 0.00 | core | V198L |
| 206 | 102.0 | 0.54 | surface | Q206L, Q206H |
| 208 | 1.1 | 0.01 | core | G208A |
| 223 | 27.3 | 0.19 | core | P223A |
| 235 | 85.3 | 0.57 | surface | D235T |
| 238 | 84.0 | 0.69 | surface | S238G |
| 241 | 25.4 | 0.17 | core | T241I |
| 242 | 53.0 | 0.47 | surface | A242P |
| 262 | 41.6 | 0.23 | core | L262H, L262M, L262I |
| 263 | 126.4 | 0.69 | surface | E263Q |
| 266 | 97.8 | 0.87 | surface | A266D, A266G |
| 269 | 50.3 | 0.41 | surface | S269H, S269A |
| 282 | 77.9 | 0.43 | surface | L282S |
| 292 | 56.5 | 0.25 | core | Y292G, Y292R, Y292F, Y292L |
| 293 | 45.9 | 0.22 | core | M293T |
| 302 | 3.1 | 0.01 | core | W302F |
| 308 | 38.6 | 0.32 | surface | S308C |
| 311 | 94.8 | 0.65 | surface | T311A |
| 312 | 97.9 | 0.62 | surface | N312D |
| 323 | 26.4 | 0.18 | core | D323E/G324R |
| 324 | 10.1 | 0.12 | core | D323E/G324R |
| 330 | 75.1 | 0.42 | surface | L330S, L330I, L330A |
| 331 | 116.1 | 0.48 | surface | R331G |
| 334 | 56.3 | 0.39 | surface | P334N, P334T, P334A |
| 337 | 109.0 | 0.45 | surface | R337Y |
| 339 | 91.4 | 0.43 | surface | K339T, K339R |
| 342 | 75.3 | 0.53 | surface | P342L/Q343E, P342L, P342W, P342R |
| 343 | 98.0 | 0.52 | surface | Q343L, Q343G, P342L/Q343E, Q343A, Q343I, Q343W, Q343P, Q343E |
| 348 | 7.9 | 0.09 | core | G348A |
| 351 | 57.4 | 0.36 | surface | V351D |
| 356 | 102.8 | 0.43 | surface | R356E |
| 363 | 178.5 | 0.74 | surface | R363G |

**Supplementary Table S11. Cell lysates activity levels of InDel variants of *wt*PTE improved in arylesterase activity.**

Changes in phosphotriesterase (native substrate: paraoxon) and arylesterase (promiscuous substrates: 4-NPB or 2-NH) activities are determined relative to those of *wt*PTE by comparing the initial rates in cell lysates measured under identical conditions with 200 μM of the respective substrates (see Methods). Data (also plotted on Figure 6) are averages of triplicate values from three independent experiments and error values represent +/- 1 SEM.

**Supplementary Table S11A: Promiscuous activity against 4-NPB.**

| | | **Activity relative to *wt*PTE** | |
|---|---|---|---|
| | | **Arylesterase** | **Phosphotriesterase** |
| **Library** | **Protein mutation [a]** | **4-NPB** | **Paraoxon** |
| **-3 bp** | ΔQ206 | 2.3 ± 0.2 | 0.7 ± 0.2 |
| | ΔD232 | 2.2 ± 0.1 | 0.06 ± 0.01 |
| | ΔT234 | 2.3 ± 0.1 | 0.41 ± 0.03 |
| | ΔH254 | 2.4 ± 0.2 | < 0.01 |
| | ΔS269 | 2.01 ± 0.01 | 0.86 ± 0.04 |
| | L272R/ΔG273 | 2.37 ± 0.01 | 0.16 ± 0.01 |
| **-6 bp** | ΔH257S258 | 5.2 ± 0.8 | 0.06 ± 0.01 |
| | ΔS258A259 | 2.2 ± 0.3 | 0.07 ± 0.01 |
| | ΔL262E263/D264H | 2.1 ± 0.8 | 0.08 ± 0.01 |
| | ΔS267A268 or ΔS269A270 | 2.3 ± 0.1 | 0.25 ± 0.04 |
| | ΔA270L271 | 3.0 ± 0.4 | 0.18 ± 0.01 |
| | ΔL271L272 | 3 ± 0.3 | 0.16 ± 0.02 |
| | ΔG273I274 | 3.2 ± 0.3 | 0.14 ± 0.01 |
| **-9 bp** | ΔG261L262E263 [b] | 5.1 ± 0.1 | 0.17 ± 0.01 |
| | ΔA268S269A270 | 2.6 ± 0.1 | 0.09 ± 0.01 |
| | ΔL272G273I274 | 2.4 ± 0.1 | 0.24 ± 0.01 |
| **+3 bp** | Y255a | 2.3 ± 0.5 | 0.01 ± 0.01 |
| | L256a | 14.4 ± 1.5 | 0.01 ± 0.01 |
| | H257a | 2.9 ± 0.4 | 0.17 ± 0.02 |
| | D258a, A259T | 3.6 ± 0.8 | 0.34 ± 0.08 |
| | V258a, A259P | 3.2 ± 1.3 | 0.8 ± 0.3 |
| | I258a | 2.8 ± 0.9 | 0.5 ± 0.2 |
| | P259a | 3.4 ± 0.2 | 0.95 ± 0.1 |
| | A259G/S259a | 4.1 ± 0.1 | 0.37 ± 0.03 |
| | S261a | 2.1 ± 0.1 | 0.78 ± 0.01 |
| | P261a | 2.7 ± 0.1 | 0.95 ± 0.01 |
| | W261a | 2.4 ± 0.4 | 0.11 ± 0.02 |
| | S319a | 2.5 ± 0.1 | 1.24 ± 0.03 |
| **+6 bp** | G255aA255b | 9.4 ± 1 | 0.05 ± 0.01 |
| | D258aR258b/A259S | 4 ± 0.1 | 0.21 ± 0.01 |
| | P261aC261b | 2.6 ± 0.1 | 0.21 ± 0.01 |
| | I261aG261b | 3.3 ± 0.1 | 1.4 ± 0.1 |

(Table S11 continued)

| | | | |
|---|---|---|---|
| | E261aS261b | 4 ± 0.2 | 0.7 ± 0.1 |
| | V262aL262b | 4.8 ± 0.2 | 1.25 ± 0.03 |
| | W262aK262b | 4.3 ± 0.1 | 0.46 ± 0.05 |
| | L265aP265b | 2.8 ± 0.1 | 1.03 ± 0.06 |
| | G265aY265b/A266S | 2.9 ± 0.1 | 1.2 ± 0.1 |
| | G266aR266b | 2.6 ± 0.1 | 0.8 ± 0.1 |
| | Y268aR268b | 2.9 ± 0.1 | 0.16 ± 0.01 |
| | Q271aR271b | 3 ± 0.1 | 0.13 ± 0.01 |
| | R271aC271b | 2.8 ± 0.1 | 0.12 ± 0.01 |
| | Y275aG275b | 3.1 ± 0.1 | 0.09 ± 0.01 |
| **+9 bp** | T121aS121bD121c | 2.8 ± 0.8 | 0.21 ± 0.06 |
| | K257aH257bG257c | 4.5 ± 0.2 | 0.17 ± 0.01 |
| | S258aG258bF258c | 2.3 ± 0.3 | 0.25 ± 0.01 |
| | C261aK261bL261c | 5.4 ± 0.3 | 0.22 ± 0.01 |
| | D261aT261bS261c | 3.3 ± 0.1 | 1.2 ± 0.1 |
| | D261aW261bK261c | 2.9 ± 0.1 | 0.43 ± 0.01 |
| | H261aI261bL261c | 4.9 ± 0.2 | 0.22 ± 0.01 |
| | V261aN261bG261c | 2.6 ± 0.1 | 0.18 ± 0.01 |
| | G262aL262bE262c/E263K | 2.3 ± 0.4 | 1 ± 0.2 |
| | L268aG268bC268c/S269P | 2.1 ± 0.1 | 0.13 ± 0.01 |
| | S269aG269bS269c | 2.4 ± 0.1 | 0.34 ± 0.02 |
| | T269aS269bG269c | 2.2 ± 0.2 | 0.34 ± 0.03 |
| | E276aG276bM276c | 2.4 ± 0.1 | 0.12 ± 0.01 |
| | A309aA309bA309c | 2.1 ± 0.1 | 0.28 ± 0.01 |

**Supplementary Table S11B: Promiscuous activity against 2-NH**

| | | Activity relative to *wt*PTE | |
|---|---|---|---|
| | | **Arylesterase** | **Phosphotriesterase** |
| **Library** | **Protein mutation [a]** | **2-NH** | **Paraoxon** |
| **-3 bp** | ΔG273 | 3.1 ± 0.3 | 0.21 ± 0.01 |
| | ΔL303 | 3.3 ± 0.5 | < 0.01 |
| | ΔS308 | 8.7 ± 1.2 | 0.41 ± 0.04 |
| | ΔT311 | 6.3 ± 0.5 | 0.48 ± 0.03 |
| | ΔV316 | 4.1 ± 0.5 | 0.04 ± 0.01 |
| | ΔM317 | 9.5 ± 0.7 | < 0.01 |
| **-6 bp** | ΔI260G261 | 2.6 ± 0.1 | 0.02 ± 0.01 |
| | I313N/ΔM314D315 | 5.2 ± 0.2 | < 0.01 |
| **-9 bp** | ΔP256H257S258 | 8.7 ± 0.5 | < 0.01 |
| | ΔG261L262E263 [b] | 3.6 ± 0.3 | 0.17 ± 0.01 |
| | ΔA270L271L272G273 | 10.5 ± 2 | 0.09 ± 0.01 |
| **+3 bp** | T311a | 8.1 ± 0.1 | 0.93 ± 0.06 |
| | G311a | 15.7 ± 3.2 | 0.28 ± 0.04 |
| | P311a | 5.7 ± 0.3 | 0.24 ± 0.02 |
| | I313R/F313a | 14.3 ± 0.5 | 0.11 ± 0.03 |

(Table S11 continued)

| | | | |
|---|---|---|---|
| | I313M/F313a | 5 ± 0.2 | < 0.01 |
| **+6 bp** | V99G/Q99aI99b | 6.5 ± 0.5 | 0.5 ± 0.1 |
| | P256R/G256aA256b | 138.6 ± 11.3 | < 0.01 |
| | S256aG256b | 34.5 ± 1.1 | 0.7 ± 0.1 |
| | V256aW256b | 9.0 ± 1.7 | 0.07 ± 0.04 |
| | H257Q/T257aY257b | 4.9 ± 1 | 1.5 ± 0.5 |
| | I313K/V313aV313b | 10.6 ± 1 | 0.04 ± 0.01 |
| | I313S/S313aL313b | 10 ± 1 | 0.02 ± 0.01 |
| **+9 bp** | V310A/S310aD310bI310c | 5.2 ± 0.5 | 0.31 ± 0.02 |
| | T311K/P311aE311bA311c | 3.9 ± 0.1 | < 0.01 |
| | T311S/M311aV311bS311c | 3 ± 0.1 | 0.36 ± 0.02 |

[a] The symbol Δ before a residue (or a group of residues) signifies that this (or these) residue(s) have been deleted. Inserted residues are labelled using the number of the position after which they are inserted and alphabetical order (e.g., glutamine and tyrosine residues inserted in this order after the residues at position 230 would be labelled Q230aY230b).
[b] This variant (ΔG261L262E263) was found in both screening campaign against 4-NPB and 2-NH.

**Supplementary Table S12. Sequence analysis of naïve TRIAD libraries focused on Loop 7 of *wt*PTE.**

Sequences were determined from randomly chosen variants upon generation of the libraries. Residues are numbered according to the crystal structure of *wt*PTE (PDB: 4PCP). *Occurrence* refers to the number of times that a specific mutation was observed among the sequenced variants.

| Library | Variant number | DNA mutation | Length change (bp) | Protein mutation | Occurrence |
|---|---|---|---|---|---|
| -3 bp | 1 | GGT(CTAG)AAG | -4 bp | frameshift | n.a. |
| | 2 | C(TGG)GT | -3 bp | L272R/ΔG273 | 8 |
| | 3 | AG(TGCG)A | -4 bp | frameshift | n.a. |
| | 4 | TC(AGC)C | -3 bp | ΔA270 | 7 |
| | 5 | CT(GGGT)A | -4 bp | frameshift | n.a. |
| | 6 | GA(CCA)T | -3 bp | ΔH254 | 3 |
| | 7 | C(TGG)GT | -3 bp | L272R/ΔG273 | 8 |
| | 8 | GG(TCT)A | -3 bp | ΔL262 | 9 |
| | 9 | ATT(CCG)CAC | -3 bp | ΔP256 | 4 |
| | 10 | CT(AGA)C | -3 bp | ΔD253 | 2 |
| | 11 | GA(CCA)T | -3 bp | ΔH254 | 3 |
| | 12 | ATT(CCG)CAC | -3 bp | ΔP256 | 4 |
| | 13 | A(TTG)GT | -3 bp | I260S/ΔG261 | 2 |
| | 14 | C(TGG)GT | -3 bp | L272R/ΔG273 | 8 |
| | 15 | CGT(TCG)TGG | -3 bp | ΔS276 | 3 |
| | 16 | C(TGG)GT | -3 bp | L272R/ΔG273 | 8 |
| | 17 | A(TTG)GT | -3 bp | I260S/ΔG261 | 2 |
| | 18 | TC(AGC)C | -3 bp | ΔA270 | 7 |
| | 19 | TC(AGC)C | -3 bp | ΔA270 | 7 |
| | 20 | GGT(CTA)GAA | -3 bp | ΔL262 | 9 |
| | 21 | ATT(GGT)CTA | -3 bp | ΔG261 | 1 |
| | 22 | CT(CCT)G | -3 bp | ΔL271/272 | 1 |
| | 23 | GGT(CTA)GAA | -3 bp | ΔL262 | 9 |
| | 24 | CAC(AGT)GCG | -3 bp | ΔS258 | 2 |
| | 25 | GG(TCT)A | -3 bp | ΔL262 | 9 |
| | 26 | TCG(TGG)CAA | -3 bp | ΔW277 | 2 |
| | 27 | TCA(GCC)CTC | -3 bp | ΔA270 | 7 |
| | 28 | TC(AGC)C | -3 bp | ΔA270 | 7 |
| | 29 | CTA(GAA)GAT | -3 bp | ΔE263 | 2 |
| | 30 | CAC(AGT)GCG | -3 bp | ΔS258 | 2 |
| | 31 | GCA(TCAG)CCC | -4 bp | frameshift | n.a. |
| | 32 | ATT(CCG)CAC | -3 bp | ΔP256 | 4 |
| | 33 | C(TGG)GT | -3 bp | L272R/ΔG273 | 8 |
| | 34 | CTA(GAA)GAT | -3 bp | ΔE263 | 2 |
| | 35 | T(CGT)GG | -3 bp | ΔS276 | 3 |
| | 36 | G(GTA)TT | -3 bp | G273V/ΔI274 | 1 |
| | 37 | TC(AGC)C | -3 bp | ΔA270 | 7 |
| | 38 | C(TGG)GT | -3 bp | L272R/ΔG273 | 8 |

(Table S12 continued)

| Library | Variant number | DNA mutation | Length change (bp) | Protein mutation | Occurrence |
|---|---|---|---|---|---|
| | 39 | TCG(TGG)CAA | -3 bp | ΔW277 | 2 |
| | 40 | AG(TGCG)A | -4 bp | Frameshift | n.a. |
| | 41 | C(TAG)AA | -3 bp | L262Q/ΔE263 | 2 |
| | 42 | CGT(TCG)TGG | -3 bp | ΔS276 | 3 |
| | 43 | CTA(GAC)CAT | -3 bp | ΔD253 | 2 |
| | 44 | AT(TCCG)C | -4 bp | frameshift | n.a. |
| | 45 | GG(TCT)A | -3 bp | ΔL262 | 9 |
| | 46 | C(TAG)AA | -3 bp | L262Q/ΔE263 | 2 |
| | 47 | TCA(GCC)CTC | -3 bp | ΔA270 | 7 |
| | 48 | C(TGG)GT | -3 bp | L272R/ΔG273 | 8 |
| | 49 | GG(TCT)A | -3 bp | ΔL262 | 9 |
| | 50 | GGT(CTAG)AAG | -4 bp | frameshift | n.a. |
| | 51 | GG(TCT)A | -3 bp | ΔL262 | 9 |
| | 52 | AAT(GCG)AGT | -3 bp | ΔA266 | 1 |
| | 53 | C(TGG)GT | -3 bp | L272R/ΔG273 | 8 |
| | 54 | GG(TCT)A | -3 bp | ΔL262 | 9 |
| | 55 | ATT(CCG)CAC | -3 bp | ΔP256 | 4 |
| | 56 | GA(CCA)T | -3 bp | ΔH254 | 3 |
| | 57 | AG(TGCG)A | -4 bp | frameshift | n.a. |
| | 58 | GG(TCT)A | -3 bp | ΔL262 | 9 |
| | 59 | GCC(CTCC)TGG | -4 bp | frameshift | n.a. |
| -6 bp | 1 | CTC(CTGGGT)ATT | -6 bp | ΔL272G273 | 2 |
| | 2 | GC(GAGTGC)ATCA | -6 bp | ΔS267A268 | 4 |
| | 3 | ATC(GGTCTA)GAC | -6 bp | ΔG251L252 | 2 |
| | 4 | G(GTCTAG)AA | -6 bp | ΔG261L262 | 5 |
| | 5 | CT(CCTGGGT)ATTC | -7 bp | frameshift | n.a. |
| | 6 | ATC(GGTCTAGA)CCA | -8 bp | frameshift | n.a. |
| | 7 | GG(TCTAGA)AGAT | -6 bp | ΔL262E263 | 8 |
| | 8 | T(CGTGGC)AA | -6 bp | S276stop | 1 |
| | 9 | ATT(CGTTCG)TGG | -6 bp | ΔR275S276 | 1 |
| | 10 | AT(TGGTCT)A | -6 bp | ΔG261L262 | 5 |
| | 11 | GG(TCTAGA)A | -6 bp | ΔL262E263 | 8 |
| | 12 | AT(TGGTCTAG)A | -8 bp | frameshift | n.a. |
| | 13 | AT(TGGTCT)A | -6 bp | ΔG261L262 | 5 |
| | 14 | GC(GAGTGC)A | -6 bp | ΔS267A268 | 4 |
| | 15 | CGT(TCGTGG)CAA | -6 bp | ΔS276W277 | 4 |
| | 16 | CTC(CTGGGTA)TTC | -7 bp | frameshift | n.a. |
| | 17 | GGT(CTAGAA)GAT | -6 bp | ΔL262E263 | 8 |
| | 18 | CT(GGGTAT)T | -6 bp | ΔG273I274 | 3 |
| | 19 | GG(TCTAGA)A | -6 bp | ΔL262E263 | 8 |
| | 20 | C(TAGAAG)AT | -6 bp | L262H/ΔE263D264 | 6 |
| | 21 | C(TCCTGG)GT | -6 bp | L271R/ΔL272G273 | 1 |
| | 22 | CT(GGGTAT)T | -6 bp | ΔG273I274 | 3 |
| | 23 | GG(TCTAGA)A | -6 bp | ΔL262E263 | 8 |
| | 24 | GG(TCTAGA)A | -6 bp | ΔL262E263 | 8 |

(Table S12 continued)

| Library | Variant number | DNA mutation | Length change (bp) | Protein mutation | Occurrence |
|---|---|---|---|---|---|
| | 25 | ATT(CCGCAC)AGT | -6 bp | ΔP256H257 | 7 |
| | 26 | AT(CGGTCTA)G | -7 bp | frameshift | n.a. |
| | 27 | AT(TGGTCT)A | -6 bp | ΔG261L262 | 5 |
| | 28 | C(TAGAAG)AT | -6 bp | L262H/ΔE263D264 | 6 |
| | 29 | TCG(TGGCAA)ACA | -6 bp | ΔW276Q277 | 1 |
| | 30 | C(TAGAAG)AT | -6 bp | L262H/ΔE263D264 | 6 |
| | 31 | CTC(ATCGGTC)TAG | -7 bp | frameshift | n.a. |
| | 32 | C(ACAGTG)CG | -6 bp | H257P/ΔS258A259 | 2 |
| | 33 | AT(TGGTCT)A | -6 bp | ΔG261L262 | 5 |
| | 34 | CTC(CTGGGTA)TTC | -7 bp | frameshift | n.a. |
| | 35 | AT(TCCGCA)C | -6 bp | ΔP256H257 | 7 |
| | 36 | CTC(CTGGGT)ATT | -6 bp | ΔL272G273 | 2 |
| | 37 | ATT(CCGCAC)AGT | -6 bp | ΔP256H257 | 7 |
| | 38 | ATT(GGTCTAGA)AG | -7 bp | frameshift | n.a. |
| | 39 | T(CAGCCC)TC | -6 bp | S269F/ΔA270L271 | 2 |
| | 40 | C(ACAGTG)CG | -6 bp | H257P/ΔS258A259 | 2 |
| | 41 | GG(TCTAGA)A | -6 bp | ΔL262E263 | 8 |
| | 42 | CGT(TCGTGGC)AAA | -7 bp | frameshift | n.a. |
| | 43 | CC(GCACAG)T | -6 bp | ΔH257S258 | 1 |
| | 44 | CGT(TCGTGG)CAA | -6 bp | ΔS276W277 | 4 |
| | 45 | GC(GAGTGC)A | -6 bp | ΔS267A268 | 4 |
| | 46 | TC(AGCCCTC)C | -7 bp | frameshift | n.a. |
| | 47 | GGT(CTAGAA)GAT | -6 bp | ΔL262E263 | 8 |
| | 48 | GG(TCTAGA)A | -6 bp | ΔL262E264 | 1 |
| | 49 | CTA(GACCATA)TTC | -7 bp | frameshift | n.a. |
| | 50 | AT(TCCGCA)C | -6 bp | ΔP256H257 | 7 |
| | 51 | CGT(TCGTGG)CAA | -6 bp | ΔS276W277 | 4 |
| | 52 | GCC(CTCCTG)GGT | -6 bp | ΔL271L272 | 2 |
| | 53 | ATT(CCGCAC)AGT | -6 bp | ΔP256H257 | 7 |
| | 54 | C(TAGAAG)AT | -6 bp | L262H/ΔE263D264 | 6 |
| | 55 | ATT(CCGCAC)AGT | -6 bp | ΔP256H257 | 7 |
| | 56 | CAT(ATTCCG)CAC | -6 bp | ΔI255P256 | 3 |
| | 57 | CT(AGACCA)T | -6 bp | ΔD253H254 | 2 |
| | 58 | CAT(ATTCCG)CAC | -6 bp | ΔI255P256 | 3 |
| | 59 | C(TAGAAG)AT | -6 bp | L262H/ΔE263D264 | 6 |
| | 60 | ATT(CCGCAC)AGT | -6 bp | ΔP256H257 | 7 |
| | 61 | C(TAGAAG)AT | -6 bp | L262H/ΔE263D264 | 6 |
| | 62 | GC(CCTCCT)G | -6 bp | ΔL271L272 | 2 |
| | 63 | AA(TGCGAGT)G | -7 bp | frameshift | n.a. |
| | 64 | CTA(GACCAT)ATT | -6 bp | ΔD253H254 | 2 |
| | 65 | GC(GAGTGC)A | -6 bp | ΔS267A268 | 4 |
| | 66 | CTC(CTGGGTA)TTC | -7 bp | frameshift | n.a. |
| | 67 | C(ACAGTGC)GA | -7 bp | frameshift | n.a. |
| | 68 | T(CAGCCC)TC | -6 bp | S269F/ΔA270L271 | 2 |
| | 69 | CT(GGGTAT)T | -6 bp | ΔG273I274 | 3 |

(Table S12 continued)

| Library | Variant number | DNA mutation | Length change (bp) | Protein mutation | Occurrence |
|---|---|---|---|---|---|
| | 70 | G(ACCATA)TT | -6 bp | D253V/ΔH254I255 | 1 |
| | 71 | GC(ATCAGC)C | -6 bp | ΔS269A270 | 1 |
| | 72 | CAT(ATTCCG)CAC | -6 bp | ΔI255P256 | 3 |
| | 73 | CGT(TCGTGG)CAA | -6 bp | ΔS276W277 | 4 |
| | 74 | AT(CGGTCT)A | -6 bp | ΔG251L252 | 2 |
| | 75 | G(CCCTCC)TG | -6 bp | A270V/ΔL271L272 | 1 |
| | 76 | GGT(CTAGACC)ATA | -7 bp | frameshift | n.a. |
| | 77 | TC(AGCCCT)C | -6 bp | ΔA270L271 | 1 |
| +3 bp | 1 | GGT+A+CT | +1 bp | frameshift | n.a. |
| | 2 | CA+AAA+T | +3 bp | H254Q/N254a | 1 |
| | 3 | C+GGT+TA | +3 bp | R261a | 1 |
| | 4 | GGT+TTT+CTA | +3 bp | F251a | 1 |
| | 5 | C+ATT+TG | +3 bp | H271a | 2 |
| | 6 | G+GCT+AA | +3 bp | E263G/Stop | 1 |
| | 7 | CTC+AT+C | +2 bp | frameshift | n.a. |
| | 8 | CA+CCA+A | +3 bp | H277a | 1 |
| | 9 | CTA+CTA+GAA | +3 bp | L262a | 4 |
| | 10 | C+GGC+AC | +3 bp | R256a | 1 |
| | 11 | ATT+ATG+CCG | +3 bp | M255a | 1 |
| | 12 | T+TG+CGT | +2 bp | frameshift | n.a. |
| | 13 | GA+GCT+C | +3 bp | D253E/L253a | 1 |
| | 14 | CT+TGT+A | +3 bp | V262a | 1 |
| | 15 | GGT+GTT+CTA | +3 bp | V261a | 1 |
| | 16 | GA+GA+CC | +2 bp | frameshift | n.a. |
| | 17 | GGTC+A+TAGA | +1 bp | frameshift | n.a. |
| | 18 | GA+ATA+CCAT | +3 bp | D253E/Y253a | 1 |
| | 19 | G+AT+CGA | +2 bp | frameshift | n.a. |
| | 20 | ATT+AGT+GGT | +3 bp | S260a | 1 |
| | 21 | GG+GTC+T | +3 bp | S261a | 1 |
| | 22 | CT+GTG+C | +3 bp | C271a | 1 |
| | 23 | GGT+TA+CTAG | +2 bp | frameshift | n.a. |
| | 24 | G+AGG+CG | +3 bp | E265a | 1 |
| | 25 | CTA+T+GA | +1 bp | frameshift | n.a. |
| | 26 | AT+GAT+T | +3 bp | M259a | 1 |
| | 27 | C+TA+CGC | +2 bp | frameshift | n.a. |
| | 28 | CTC+CAT+CTG | +3 bp | H271a | 2 |
| | 29 | GGT+TG+CTAG | +2 bp | frameshift | n.a. |
| | 30 | CT+CTT+C | +3 bp | F271a | 3 |
| | 31 | ATT+TCC+CCG | +3 bp | S255a | 1 |
| | 32 | C+ATC+TA | +3 bp | H261a | 1 |
| | 33 | CT+CTT+G | +3 bp | L272a | 1 |
| | 34 | CT/A+G+AGA | +1 bp | frameshift | n.a. |
| | 35 | T+GTT+CA | +3 bp | C268a | 1 |
| | 36 | GGT+TAG+CTA | +3 bp | Stop | 2 |
| | 37 | CCGC+CGC+AC | +3 bp | P256a | 1 |

(Table S12 continued)

| Library | Variant number | DNA mutation | Length change (bp) | Protein mutation | Occurrence |
|---|---|---|---|---|---|
| | 38 | AA+AG+TG | +2 bp | frameshift | n.a. |
| | 39 | CGTT+GAT+CG | +3 bp | Stop | 2 |
| | 40 | CTC+TTT+CTG | +3 bp | F271a | 3 |
| | 41 | CGT+A+TC | +1 bp | frameshift | n.a. |
| | 42 | CT+TCT+A | +3 bp | L262a | 4 |
| | 43 | C+GT+CGC | +2 bp | frameshift | n.a. |
| | 44 | CTA+TCA+GAC | +3 bp | S252a | 1 |
| | 45 | C+AGT+TG | +3 bp | Q271a | 1 |
| | 46 | GG+CAT+T | +3 bp | I261a | 1 |
| | 47 | TCA+TCA+GCC | +3 bp | S269a | 1 |
| | 48 | ATC+AT+GGTC | +2 bp | frameshift | n.a. |
| | 49 | CT+TCT+A | +3 bp | L262a | 4 |
| | 50 | GT+AC+CTAG | +2 bp | frameshift | n.a. |
| | 51 | GGT+GAA+CTA | +3 bp | E261a | 1 |
| | 52 | A+ATA+CA | +3 bp | N278a | 1 |
| | 53 | ATT+CTT+CCG | +3 bp | L255A | 1 |
| | 54 | CT+TAG+G | +3 bp | R272a | 1 |
| | 55 | GGT+GCT+CTA | +3 bp | A261a | 2 |
| | 56 | GGT+A+CT | +1 bp | frameshift | n.a. |
| | 57 | CC+C+GCA | +1 bp | frameshift | n.a. |
| | 58 | CT+CG+AG | +2 bp | frameshift | n.a. |
| | 59 | CT+TGT+A | +3 bp | V252a | 1 |
| | 60 | ATT+ATT+CCG | +3 bp | I255a | 1 |
| | 61 | CT+TCA+A | +3 bp | Q262a | 1 |
| | 62 | CT+TC+GG | +2 bp | frameshift | n.a. |
| | 63 | CT+TTT+A | +3 bp | L262a | 4 |
| | 64 | C+CTC+TA | +3 bp | P251a | 1 |
| | 65 | GAC+TC+C | +2 bp | frameshift | n.a. |
| | 66 | GGT+GG+CTAG | +2 bp | frameshift | n.a. |
| | 67 | GGTC+CAT+TA | +3 bp | P261a | 2 |
| | 68 | GCGA+CGA+TT | +3 bp | T259a | 1 |
| | 69 | CTC+TTT+CTG | +3 bp | F271a | 3 |
| | 70 | CCG+TC+C | +2 bp | frameshift | n.a. |
| | 71 | GCGAT+GAG+T | +3 bp | I260M/S260a | 1 |
| | 72 | CTC+CCC+CTG | +3 bp | P271a | 1 |
| | 73 | GG+CTA+T | +3 bp | Y261a | 1 |
| | 74 | GGT+GCT+CTA | +3 bp | A261a | 2 |
| | 75 | T+CT+GGC | +2 bp | frameshift | n.a. |
| | 76 | GGT+CCT+GTA | +3 bp | P261a | 2 |
| | 77 | GGTC+CA+TAG | +2 bp | frameshift | n.a. |

**Supplementary Table S13. Methods developed for the generation of libraries with random insertions, repeats and/or deletions.**

| Method | Principle | Mutational scope (Type of mutations/Number per target sequence) | Frameshift InDels (%) | Reference |
|---|---|---|---|---|
| **RID** | Random Insertion/Deletion mutagenesis. (1) A circular single-stranded DNA (ssDNA) corresponding to the sense chain of the target gene is produced from the linear double-stranded target gene by linker ligation, restriction digestion, circularization by self-ligation and exonuclease digestion to remove the anti-sense chain. (2) Random cleavage (linearization) of the circular ssDNA at single positions by treatment with Ce(IV)‐EDTA complex. (3) Ligation of 5'- and 3'- anchors at both ends of the ssDNA. These anchors are designed differently depending whether a deletion or an insertion is to be introduced. (4) PCR amplification of the DNAs linked to the two anchors at both ends. (5) Digestion by a type IIS restriction enzyme (*e.g.*, BciVI) removes the anchor and leaves a deletion or an insertion in the target gene (depending on how the anchors' sequences have been designed). (6) Reconstitution of the target gene by self-ligation (re-circularization) and linearization by restriction digestion. The resulting products can then be cloned in a vector to finalize the variant library. | One single InDel per variant; the procedure also generates random point substitutions presumably during the PCR step.**[a]** | ~10% | 17 |
| **Segmental mutagenesis** | (1) The vector is first linearized either at the 5′ or 3′ end of the target gene. (2) Progressive BAL-31 exonuclease action and removal of the remaining vector DNA yields two batches of either 3′ or 5′ truncated gene fragments. (3) Combinatorial assembly of these two ends to generate variants of the target gene yields the segmental mutagenesis library which is then igated into a vector and transformed into *E.coli*. | One single deletion or one tandem repeat per variant (on a defined region of the target gene) | ~66% | 18 |
| **RAISE** | RAndom Insertional-deletional Strand Exchange mutagenesis. (1) The target gene is fragmented using DNAseI. (2) The obtained fragments are extended randomly using Terminal deoxynucleotidyl transferase (TdT). (3) Assembly PCR with the TdT-extended fragment results in the shuffling of InDels and substitutions (generated by TdT or during the PCR steps) within the target gene. | Combination of region-exchanged mutations and substitutions. **[b]** | ~66% | 19 |
| **COBARDE** | Codon-based random deletion mutagenesis. (1) Chemical synthesis (based on the phosphoramidite method) generating a population of mutagenic oligonucleotides with multiple codon deletions in reference to the target gene. The mutagenic process consists of multiple successive cycles of 3-nucleotide extension as follow: (i) transient blockage of a fraction of the synthesized oligos, (ii) extension of the unblocked oligos by 3 nucleotides, (iii) removal of the blocking groups. (2) The resulting oligonucleotide mixture (corresponding the deletion variants) is purified, duplexed using a DNA polymerase and ligated into a vector. | One or multiple combined codon-based deletions per variant (usually on a defined region of the target gene). **[c]** | <5% | 20 |
| **TRINS** | Tandem repeat insertion (TRINS). (1) The target gene is fragmented using DNAseI. (2) An aliquot of the generated fragments is converted into single-stranded circular DNA using CircLigase. (3) Tandem repeats are generated by mixing linear fragments together with circularized fragments in an assembly PCR reaction involving rolling-circle polymerisation. (4) Assembly PCR products are then cloned to finalized the TRINS library. | One or multiple tandem repeats per variants. Tandem repeat size variable (depending on the size of the initial DNAseI linear fragment). **[d]** | ~66% | 21 |
| **Pentapeptide scanning** | (1) An engineered transposon is randomly inserted within the vector containing the target gene by *in vivo* or *in vitro* reaction (depending on the type of transposon | One single insertion of defined size and sequence (5 nucleotide triplets) per variant. | Not reported | |

| | | | | |
|---|---|---|---|---|
| | used). The sub-library consisting of only of the target gene with a single transposon insertion can be isolated by DNA electrophoresis and size selection. (2) Restriction digestion (*e.g.,* with NotI in the case of modified Mu transposon) leaves a 15 bp insertion after self-ligation of the target gene. | | | 22, 23 |
| **TND** | Triplet nucleotide deletion<br>1) An engineered transposon (dubbed MuDel) is randomly inserted within the vector containing the target gene using *in vitro*. The sub-library consisting of only of the target gene with a single transposon insertion can be isolated by DNA electrophoresis and size selection. (2) Digestion with type IIS restriction enzyme MlyI, results in a triplet deletion upon self-ligation of the target gene. | One single nucleotide triplet deletion per variant. **[e]** | Not reported | 4, 24 |
| **CDM** | Codon Deletion Mutagenesis<br>1) The target gene is cloned in a vector such as the resulting protein is N-terminally fused to an intein. (2) An engineered transposon (dubbed MuCDM) is then randomly inserted within the target gene using *in vitro*. MuCDM contains an intein sequence fused to an antibiotic resistance (*e.g.*, TEM1), thus enabling selection only if transposon insertion is in the reading frame of the target gene. (3) An inverse PCR reaction with primers based on the transposon's terminal sequences is performed to amplify the vector from the transposon's insertion point. These primers carry a carefully positioned type IIS restrictions site (*e.g.*, for BsgI) to remove a specific number of nucleotides from the resulting inverse PCR product. (3) Digestion by the type IIS restriction enzyme removes 1 to 5 nucleotide codons from the target gene depending on the positioning of the recognition sequence on the primers. (4) CDM libraries are generated upon self-ligation and transformation of the vector carrying the target gene variants in *E. coli*. | Deletions of one to five consecutive codons. **[f]** | <10% | 25 |
| **Extensive gene truncation** | 1) An engineered transposon (MuDel) is randomly inserted within the target gene by *in vitro* transposition. (2) 5′ and 3′ fragment sub-libraries of the target gene are amplified in two separate PCR reactions. In each reaction, one primer is complementary to the 5′ or 3′ constant regions of the target gene (adding BsaI at these ends), and the other to a sequence located in the transposon. (3) Digestion with BsaI creates unique overhangs in each sub-library complementary to unique overhangs in a DNA linker (free of MlyI sites) to favor directional ligation between these sub-libraries and the linker (4) The product of ligation was digested with MlyI removing the transposon sequence. (5) Intramolecular blunt-end ligation results in a circular product joining the 5′ and 3′ terminal fragments of the target gene. This circularized product is a library corresponding to the target genes with extensive truncation. (7) PCR on this circular library with primers complementary to the termini of the target gene results in a linear version of the extensive truncation library. (8) The final library of truncated variants of the desired size range is isolated by gel electrophoresis and cloned in a vector. | Extensive DNA truncations of desired size range. **[g]** | Not reported | 26 |
| **InDel assembly** | Assembly approach relying on successive cycles of DNA restriction and ligation to assemble a DNA library on beads. At each assembly cycle, DNA templates immobilized on beads are restricted with a type IIs endonuclease (*e.g.*, SapI) and building blocks annealed and ligated. After ligation, the cycle can be restarted. Compositional variation is achieved primarily by combining controlled pools of building blocks of various length. | One or multiple combined codon-based insertions and deletions per variant (usually on a defined region of the target gene). **[h]** | Not reported | 27 |

(Table S13 continued)

**[a]** The RID mutagenesis was validated by randomly replacing three consecutive bases by recognition sequence for BglII (AGATCT) in the GFPuv gene: 17 variants (out of 19 randomly picked variants for sequencing; ~90%) displayed the desired mutations; 2 variants out of the pool of 19 variants (~10%) were frameshifted (deletion of 4 consecutive bases instead of 3); in addition, 6 variants out of 19 (~30%) also displayed single point substitution. The RID mutagenesis has also been applied to randomly replace three consecutive bases by a mixture of 20 codons, effectively resulting in point substitution mutants.

**[b]** RAISE was validated using TEM-1 beta-lactamase as target gene. After transformation of the library (~2,000 variants), 41 colonies were randomly picked and sequenced leading to the identification of region-exchanged mutations and point substitutions. Twenty-nine region-exchanged mutations were found in 19 variants. The number of the region-exchanged mutations per variant was 1 (12 variants), 2 (6 variants), or 5 (1 variants). Approximately two-thirds of the region-exchanged mutations were frameshifts. Seventy-nine point substitutions were identified over 34 variants, among which 15 had also region-exchanged mutations. Three variants out of the 41 that sequenced were parental sequences, presumably due to vector self-ligation.

**[c]** COBARDE was validated using a sequence of 9 residues forming the omega loop in TEM-1 beta-lactamase: 4 parental sequences (presumably vector self-ligation) and 1 frame-shift (insertion of 1 bp) were found out of 34 sequenced transformants.

**[d]** TRINS was validated using three different templates: TEM-1 beta-lactamase, m.HaeIII methyltransferase and KE70 R6 (a laboratory-evolved variant of computationally designed Kemp eliminase). Out of 35 sequenced variants (from the three naïve libraries), 27 carried one insertion per gene, 4 had two and 4 had three. Two-third of the tandem repeats (23 of 35) resulted in frameshift. The sequenced variants also carried ~2 random point substitutions per variant presumably incorporated during PCR steps.

**[e]** TND was validated using TEM-1 beta-lactamase (Jones, 2005) and eGFP (Baldwin et al., 2009; Arpino et al., 2014) as templates. In the case of TEM-1, the library generation process was combined with two consecutive selection steps: (i) selection for loss of ampicillin resistance upon transposition insertion within *bla*) and (ii) selection for retention of antibiotic resistance upon triplet nucleotide deletion. In the case of eGFP, the final library consisted of ~2,500 variants and 153 variants were chosen for sequencing based on the colony phenotype (88 fluorescent and 65 nonfluorescent). This led to the identification of 87 unique triplet deletions (out of 153): 42 triplet deletions among the 88 fluorescent variants and 45 among the 65 nonfluorescent ones. No additional point substitutions or frameshifts were observed among the sequenced variants.

**[f]** CMD was validated using super folder GFP (sfGFP) as template. Five libraries, corresponding to the deletion of 1 to 5 consecutive codons, were generated and around 20 variants from each library (amounting to a total of 104 sequences) were sequenced, showing that the majority of the variants (~92%) contained the desired deletions. Eight out of 104 sequences had either no mutations or unwanted mutations, most of them due to incomplete BsgI digestion.

**[g]** The extensive gene truncation method was validated using an artificial RNA ligase enzyme (DNA size ~ 350 bp) as template and resulted in a library with truncations up to ~235 bp. Next generation sequencing analysis of the library revealed that it contained 9,006 unique deletions (~32% of the 27,730 possible unique deletions in this size range). The distribution of deletion lengths was found to range between 6 and 235 nucleotides in length. Deletions longer than 110 bp were observed at 50% or greater of the number of all possible deletions. The library was subjected to in vitro selection and functional variants with deletions of up to 18 amino acids of the parental enzyme.

**[h]** InDel assembly was validated using part of TEM-1 beta-lactamase's omega loop (5 residues, $_{164}$RWEPE$_{168}$). The library was designed in order to explore the sequence neighbourhood of a previously reported variant ($_{164}$RYYGE$_{168}$) by using biased mixes of building blocks. The resulting library was analysed by next-generation sequencing before and after selection for ceftazidime resistance and demonstrated selective enrichment of the target sequence ($_{164}$RYYGE$_{168}$) as well as variants with extensions (*e.g.*, $_{164}$RGYMKER$_{168b}$).

**Supplementary Table S14. Oligonucleotides used in this study.**

| Experiment | Oligonucleotide name and sequence |
|---|---|
| Preparation of SubsNNN by PCR using pUC57-Del2 as template | **Subs-F:** 5'-[Phos]-ATGTCGACTCGACTAGTGCTTGGATTCTCA-3'<br>**Subs-B:** 5'-[Phos]-NNNGGGATGACTCCATGGACTTCGC-3'<br>(MlyI sites underlined) |
| TransIns adapter to generate pUC57-TransIns from pUC57-TransDel | **TransIns-F:** 5'-[Phos]-AATTCAGATCTGCGGCCGCGCACGAAAAACGCGAAAGCGTTTCACGAT-AAATGCGAAAACGGA -3'<br>**TransIns-R:** 5'-[Phos]-CTAGTCCGTTTTCGCATTTATCGTGAAACGCTTTCGCGTTTTTCGTGCG-CGGCCGCAGATCTG-3'<br>(NotI sites underlined) |
| Del3 adapter to generate pUC57-Del3 from pUC57-Del2 | **Del3-F:** 5'-[Phos]-CATGGAGTCATCCCGGGA-3'<br>**Del3-R:** 5'-[Phos]-AGCTTCCCGGGATGACTC-3' |
| Ins adapter to generate pUC57-Ins from pUC57-Del2 | **Ins-F:** 5'-[Phos]-AATTCTAGATCTGCGGCCGCATCCGTCTTCAGTCGCTGCTGA-3'<br>**Ins-R:** 5'-[Phos]-CTAGTCAGCAGCGACTGAAGACGGATGCGGCCGCAGATCTAG-3' |
| Ins1/2/3 adapter to generate libraries pUC57-Ins1/2/3 from pUC57-Ins | **Ins1/2/3-F:** 5'-[Phos]-CATGGCTGAAGGCCACTCGAGATCGAT (NNN)$_{1/2/3}$ GCGCTGACTCA-3'<br>**Ins1/2/3-R:** 5'-[Phos]-AGCTTGAGTCAGCGC (NNN)$_{1/2/3}$ ATCGATCTCGAGTGGCCTTCAGC-3' |
| Removal of MlyI recognition site in the origin of replication of pUC19 by saturation mutagenesis | **Ori-MlyI-F:** 5'-ACCCGGTAAGACACGACTTATCGCCACTGGCA-3'<br>**Ori-MlyI-B:** 5'-GTGTCTTACCGGGTTGGNNTCAAGACGATAGTTACCGGA-3' |
| Removal of AcuI recognition site in the origin of replication of pUC19 by saturation mutagenesis | **Ori-AcuI-F:** 5'-TATCTGCGCTCTGNNGAAGCCAGTTACCTT-3'<br>**Ori-AcuI-B:** 5'-AGCGCAGATACCAAATACTGTTCTTCTAGTGTAGCCGTA-3' |
| Amplification of the origin of replication of pUC19 for assembly into the pID vectors | **Ori-AflII:** 5'-GGACTTAAGGAGCAAAAGGCCAGCAAAAGG-3'<br>**Ori-SpeI:** 5'-GCACACTAGTCTCATGACCAAAATCCCTTAACG-3' |
| (1) Amplification of TetR from pASK-IBA5plus (TetR-F/TetR-B)<br>(2) Amplification of AmpR from pID-T7 (mTEM1-F/mTEM1-B)<br>(3) Overlap PCR to form AmpR-TetR operon (mTEM1-F/TetR-B) | **TetR-F:** 5'-TGATTAAGCATTGGTAGGAATTAATGATGTCTCGTT-3'<br>**TetR-B:** 5'-TTAAACTAGTGAAGTTACCATCACGGA-3'<br>**mTEM1-F:** 5'-CTGAAAGGACGTCAGGTGGCAC-3'<br>**mTEM1-B:** 5'-TAATTCCTACCAATGCTTAATCAGTGAGGCA-3' |
| Amplification of Tet promoter from pASK-IBA5plus | **Tet-prom-F:** 5'-AGGCTTAAGACATGACCCGACACCATCGA-3'<br>**Tet-prom-B:** 5'-GGCTCATATGTATATCTCCTTCTTAAAG-3' |

#### 4. SUPPLEMENTARY REFERENCES

1. Lu Q. Seamless cloning and gene fusion. *Trends Biotechnol* **23**, 199-207 (2005).

2. Afriat-Jurnou L, Jackson CJ, Tawfik DS. Reconstructing a missing link in the evolution of a recently diverged phosphotriesterase by active-site loop remodeling. *Biochemistry* **51**, 6047-6055 (2012).

3. Hoque MA*, et al.* Stepwise Loop Insertion Strategy for Active Site Remodeling to Generate Novel Enzyme Functions. *ACS Chem Biol* **12**, 1188-1193 (2017).

4. Jones DD. Triplet nucleotide removal at random positions in a target gene: the tolerance of TEM-1 beta-lactamase to an amino acid deletion. *Nucleic Acids Res* **33**, e80 (2005).

5. Kaltenbach M, Emond S, Hollfelder F, Tokuriki N. Functional Trade-Offs in Promiscuous Enzymes Cannot Be Explained by Intrinsic Mutational Robustness of the Native Activity. *PLoS Genet* **12**, e1006305 (2016).

6. Ikeda RA, Ligman CM, Warshamana S. T7 promoter contacts essential for promoter activity in vivo. *Nucleic Acids Res* **20**, 2517-2524 (1992).

7. Zhang J, Kobert K, Flouri T, Stamatakis A. PEAR: a fast and accurate Illumina Paired-End reAd mergeR. *Bioinformatics* **30**, 614-620 (2014).

8. Langmead B, Salzberg SL. Fast gapped-read alignment with Bowtie 2. *Nat Methods* **9**, 357-359 (2012).

9. Li H*, et al.* The Sequence Alignment/Map format and SAMtools. *Bioinformatics* **25**, 2078-2079 (2009).

10. Rice P, Longden I, Bleasby A. EMBOSS: the European Molecular Biology Open Software Suite. *Trends Genet* **16**, 276-277 (2000).

11. Tokuriki N, Jackson CJ, Afriat-Jurnou L, Wyganowski KT, Tang R, Tawfik DS. Diminishing returns and tradeoffs constrain the laboratory optimization of an enzyme. *Nat Commun* **3**, 1257 (2012).

12. Campbell E*, et al.* The role of protein dynamics in the evolution of new enzyme function. *Nat Chem Biol* **12**, 944-950 (2016).

13. Kaltenbach M, Jackson CJ, Campbell EC, Hollfelder F, Tokuriki N. Reverse evolution leads to genotypic incompatibility despite functional and active site convergence. *Elife* **4**,  (2015).

14. Krissinel E, Henrick K. Inference of macromolecular assemblies from crystalline state. *J Mol Biol* **372**, 774-797 (2007).

15. Miller S, Janin J, Lesk AM, Chothia C. Interior and surface of monomeric proteins. *J Mol Biol* **196**, 641-656 (1987).

16. Tokuriki N, Stricher F, Schymkowitz J, Serrano L, Tawfik DS. The stability effects of protein mutations appear to be universally distributed. *J Mol Biol* **369**, 1318-1332 (2007).

17. Murakami H, Hohsaka T, Sisido M. Random insertion and deletion of arbitrary number of bases for codon-based random mutation of DNAs. *Nat Biotechnol* **20**, 76-81 (2002).

18. Pikkemaat MG, Janssen DB. Generating segmental mutations in haloalkane dehalogenase: a novel part in the directed evolution toolbox. *Nucleic Acids Res* **30**, E35-35 (2002).

19. Fujii R, Kitaoka M, Hayashi K. Random insertional-deletional strand exchange mutagenesis (RAISE): a simple method for generating random insertion and deletion mutations. *Methods Mol Biol* **1179**, 151-158 (2014).

20. Osuna J, Yanez J, Soberon X, Gaytan P. Protein evolution by codon-based random deletions. *Nucleic Acids Res* **32**, e136 (2004).

21. Kipnis Y, Dellus-Gur E, Tawfik DS. TRINS: a method for gene modification by randomized tandem repeat insertions. *Protein Eng Des Sel* **25**, 437-444 (2012).

22. Hallet B, Sherratt DJ, Hayes F. Pentapeptide scanning mutagenesis: random insertion of a variable five amino acid cassette in a target protein. *Nucleic Acids Res* **25**, 1866-1867 (1997).

23. Hayes F, Hallet B. Pentapeptide scanning mutagenesis: encouraging old proteins to execute unusual tricks. *Trends Microbiol* **8**, 571-577 (2000).

24. Arpino JA, Reddington SC, Halliwell LM, Rizkallah PJ, Jones DD. Random single amino acid deletion sampling unveils structural tolerance and the benefits of helical registry shift on GFP folding and structure. *Structure* **22**, 889-898 (2014).

25. Liu SS, Wei X, Ji Q, Xin X, Jiang B, Liu J. A facile and efficient transposon mutagenesis method for generation of multi-codon deletions in protein sequences. *J Biotechnol* **227**, 27-34 (2016).

26. Morelli A, Cabezas Y, Mills LJ, Seelig B. Extensive libraries of gene truncation variants generated by in vitro transposition. *Nucleic Acids Res* **45**, e78 (2017).

27. Tizei PAG, Harris E, Renders M, Pinheiro VB. InDel assembly: A novel framework for engineering protein loops through length and compositional variation (2017).
